# Genetic regulation of transcriptional variation in natural *Arabidopsis thaliana* accessions

**DOI:** 10.1101/037077

**Authors:** Yanjun Zan, Xia Shen, Simon K. G. Forsberg, Örjan Carlborg

**Affiliations:** Department of Clinical Sciences, Division of Computational Genetics, Swedish University of Agricultural Sciences, Uppsala, Sweden; Usher Institute for Population Health Sciences and Informatics, University of Edinburgh, Edinburgh, United Kingdom; Department of Medical Epidemiology and Biostatistics, Karolinska Institutet, Stockholm, Sweden; MRC Human Genetics Unit, MRC Institute of Genetics and Molecular Medicine, University of Edinburgh, Edinburgh, UK

**Keywords:** eQTL mapping, RNA sequencing, Gene Expression, *Arabidopsis thaliana*, structural variation

## Abstract

An increased knowledge of the genetic regulation of expression in *Arabidopsis thaliana* is likely to provide important insights about the basis of the plant’s extensive phenotypic variation. Here, we reanalysed two publicly available datasets with genome-wide data on genetic and transcript variation in large collections of natural *A. thaliana* accessions. Transcripts from more than half of all genes were detected in the leaf of all accessions, and from nearly all annotated genes in at least one accession. Thousands of genes had high transcript levels in some accessions but no transcripts at all in others and this pattern was correlated with the genome-wide genotype. In total, 2,669 eQTL were mapped in the largest population, and 717 of them were replicated in the other population. 646 cis-eQTLs regulated genes that lacked detectable transcripts in some accessions, and for 159 of these we identified one, or several, common structural variants in the populations that were shown to be likely contributors to the lack of detectable RNA-transcripts for these genes. This study thus provides new insights on the overall genetic regulation of global gene-expression diversity in the leaf of natural *A. thaliana* accessions. Further, it also shows that strong cis-acting polymorphisms, many of which are likely to be structural variations, make important contributions to the transcriptional variation in the worldwide *A. thaliana* population.

## Introduction

Several earlier studies have utilized genome-wide data to explore the link between genetic and phenotypic diversity in natural *Arabidopsis thaliana* populations [1-10]. Some of this phenotypic variation was found to be caused by regulatory genetic variants that lead to differences in gene expression (Expression Level Polymorphisms or ELPs for short), underlying, for example, semi-dwarfism [11], changes in flowering-time [5, 12], changes in seed flotation [13], and changes in self-incompatibility [14]. By extending efforts to scan for ELPs to the whole genome and transcriptome level in large collections of natural *A. thaliana* accessions, useful insights might be gained about the link between genetic and expression level variation in the worldwide population. This may ultimately reveal interesting candidate genes and mechanisms that have contributed to adaptation to the natural environment.

Earlier studies have shown that there is a considerable transcriptional variation amongst natural *A. thaliana* accessions. A comparison of expression in shoot-tissue from two accessions, Sha and Bay-0, found that (i) 15,352 genes (64% of all tested) were expressed in the shoot in the field, (ii) 3,344 genes (14% of all tested) were differentially expressed between the accessions and (iii) 53 genes were uniquely expressed in Sha/Bay-0, respectively [15]. Later, Gan *et al.* [16] studied the seedling transcriptome from 18 natural accessions and found that 75% (20,550) of the protein-coding genes, 21% of the non-coding RNAs and 21% of the pseudogenes were expressed in the seeding tissue of at least one of the accessions. Further, they also found that 46% (9,360) of the expressed protein-coding genes were differently expressed between at least one pair of accessions [16]. Kliebenstein *et al.* [17] found that, on average, 2,234 genes were differently expressed among 7 natural accessions when treated with salicylic acid and, in the same accessions, Leeuwen *et al.* [18] found that gene-network level expression responses were genotype dependent. There are also numerous other studies that have illustrated the importance of naturally occurring ELPs (e.g. [19-21]) for the transcriptional variation among natural *A.* thaliana accessions. It is therefore expected that there will be an extensive, and genetically controlled, transcriptional variation in the worldwide *A. thaliana* population, although its extent remains to be shown.

Expression quantitative trait loci (eQTL) mapping is a useful approach to link genetic and expression variation. It has been successfully applied in many organisms, including yeast [22], plants [23], as well as animals and humans [24]. Most eQTLs are detected in the close vicinity of the gene itself (cis-eQTL) and these often explain a large proportion of the observed expression variation [25-27]. Fewer eQTL (from 20-50% reported in various organisms [25, 28, 29]) are located in the remainder of the genome (trans-eQTL). Using eQTL mapping, the genetic regulation of expression variation has been dissected in *A. thaliana* using recombinant inbred lines (RIL) [20, 23, 29] and other experimental crosses [30]. These initial studies have revealed that the majority of the ELPs in *A. thaliana* are heritable, that most of the detected eQTLs are located in cis and that structural variations are a common mechanism contributing to this variation [20]. The data generated within the *A. thaliana* 1001 genomes projects now provides an opportunity to explore this expression variation also in larger collections of natural *A. thaliana* accessions that better cover the worldwide distribution of the plant [2, 32, 33].

Here, we explored the genome-wide expression variation, and the genetic regulation of gene expression, using a dataset generated from 144 natural *Arabidopsis thaliana* accessions (*SCHMITZ-data*; [32]) and replicated some of our findings in a second dataset with 107 natural *A. thaliana* accessions *(DUBIN-data;* [33]). Transcripts from a core-set of genes were present in the leaf in all accessions, but thousands of genes showed a different pattern with high transcript levels in some accessions, but no transcripts at all in others. RNA sequencing bias was a concern for lowly expressed genes, but focusing on the highly expressed genes we found a large overall contribution by genetics to this variation. Hundreds of cis-eQTL contributing to the lack of transcripts in some accessions were mapped in the larger dataset [32] and many of these replicated in the independent smaller dataset [33]. For about one quarter of the cis-eQTL genes one, or in many cases several, common structural variants were significantly associated with the lack of reads in the transcriptome analyses. This indicates that the lack of transcripts observed for many accessions in the worldwide *A.* thaliana population is often due to common deletions of transcription start sites or the whole genes. Our results thus provides an overall perspective on the transcript-level variation in natural *A. thaliana* accessions and dissect the genetics underlying the presence or absence of transcripts for individual genes in individual accessions. Overall, our results confirm that lack-of-function alleles [5, 31, 34-38], often due to structural variations [16, 20], are important contributors to the overall transcriptional variation also in the worldwide *A. thaliana* population.

## Results

### *RNA sequencing detects transcripts from nearly all TAIR10 annotated genes in the leaf of natural* A. thaliana *accessions*

We downloaded publicly available RNA-seq [32] and whole-genome SNP genotype data [2] for a population of 144 natural *A. thaliana* accessions (*SCHMITZ-data*). In this data, we first explored the variability in RNA-seq scored expression-values across 33,554 genes and 140 accessions that passed quality control (see Materials and Methods). A gene was considered as expressed if it had a normalized FPKM (Fragment Per Kilobase of exon per Million fragments mapped) value greater than zero. The available RNA-seq data was generated from a single tissue (leaf) and we therefore expected that a considerable proportion of the genes would be transcriptionally inactive due to tissue specific expression. The data, however, showed that among the 33,554 genes in the *SCHMITZ* data, only 289 lacked transcripts across all the accessions in the population (i.e. had a normalized FPKM = 0 in all accessions).

### *A core-set of genes has detectable transcripts in the leaf of all the evaluated natural* A. thaliana *accessions*

In the *SCHMIZ-data,* transcripts were detected (Normalized FPKM > 0) in all accessions for a large set of genes (18,289; Figure 1). In the *DUBIN-data,* 12,927 genes had detectable transcripts in all accessions and all of these transcripts, except 79, had detectable transcripts in all of the accession in the *SCHMITZ-data* [32] (Figure 1). Of the 10,549 genes for which transcripts were detected (RPKM > 0) only in some accessions in the DUBIN-data, transcripts were detected for 4,129 in all and 6,393 in some (RPKM > 0) accessions in the *SCHMITZ-data* (Figure 1). Hence, for only 27 genes with transcripts detected in at least one of the accessions in the DUBIN-data, no transcripts were detected in any of the accessions in the *SCHMITZ-data* (Figure 1).

**Figure 1.**
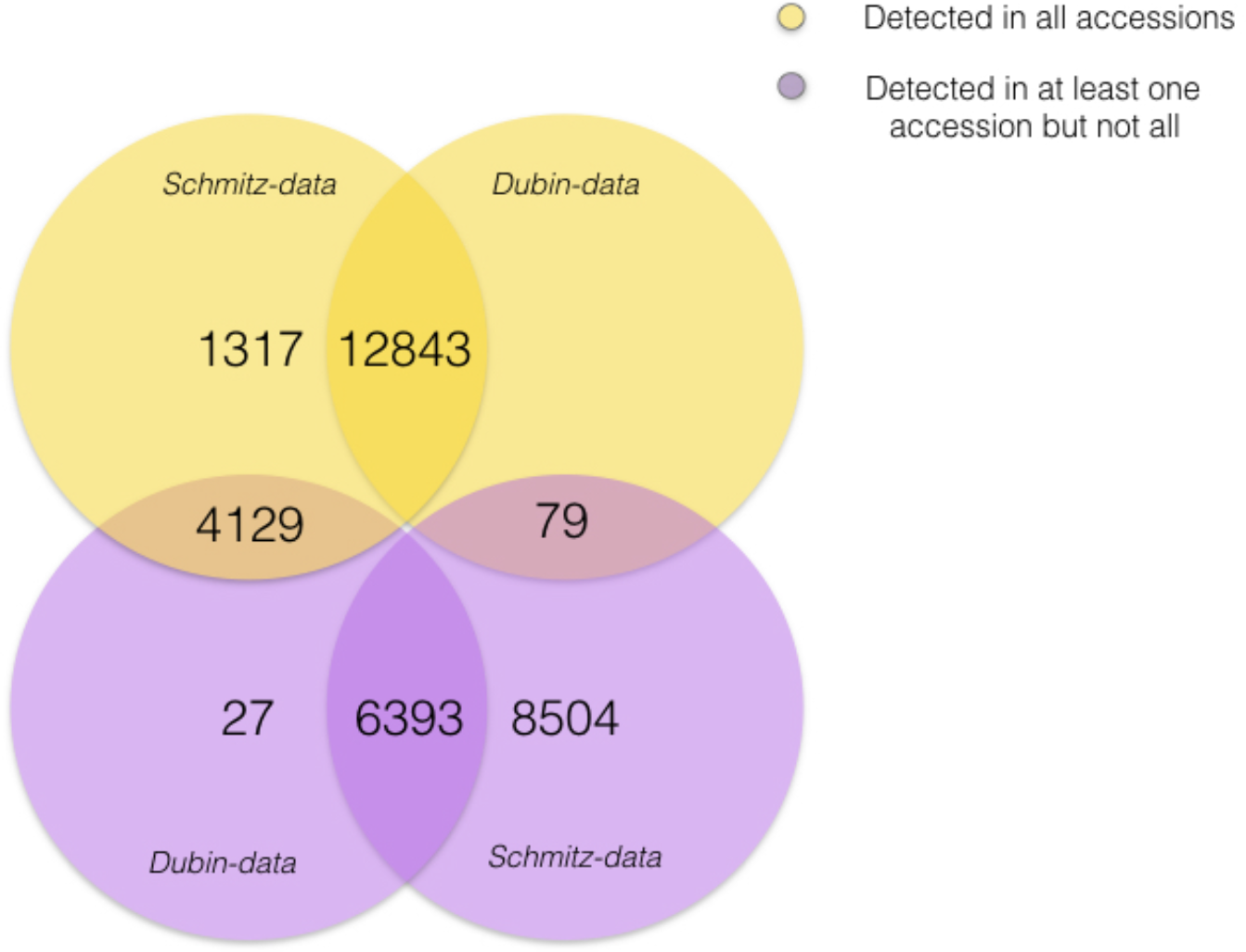
Figure 1. Overlap of RNA-seq scored transcripts in the leaf of 140 natural A. thaliana accessions [32] (SCHMITZ-data; Normalized FPKM > 0) and 107 Swedish natural A. thaliana accessions [33] (DUBIN-data; RPKM > 0). The numbers of detected transcripts in all accessions of the respective datasets are shown in yellow. The numbers of detected transcripts in at least one, but not all, of the accessions in the respective datasets are shown in purple.

The lower sequencing coverage, and more stringent filtering of the sequence reads in the *DUBIN-data* [33], likely explains why transcripts were detected for more genes in more accessions in the *SCHMITZ-data.* Few (27) genes were uniquely expressed in the *DUBIN-data,* but it is noteworthy that 13 of these are t-RNA genes and that 9 of these code for proline (Table S1).

### *The number of genes with detected transcripts in the leaf of individual* A. thaliana *accessions is highly variable*

Within the individual accessions of the *SCHMITZ-data* [32] we found transcripts (Normalized FPKM > 0) for a considerably lower number of genes than the total number of genes with detected transcripts across the entire population. The number of genes with RNA-seq detected transcripts in the individual accessions (Normalized FPKM > 0) varied from 22,574 to 26,967 with an average of 24,565 (Figure 2A). The proportion of genes with detected transcripts was thus higher in this dataset than that what has earlier been reported for shoots in two natural accessions [39], but similar to that in the study of 18 natural accessions [16] (73% in the *SCHMITZ-data,* 64% in [39] and about 70% in [16]). A similar pattern of variability in the number of genes with detected transcripts was also found in the DUBIN-data (Figure 2C).

**Figure 2.**
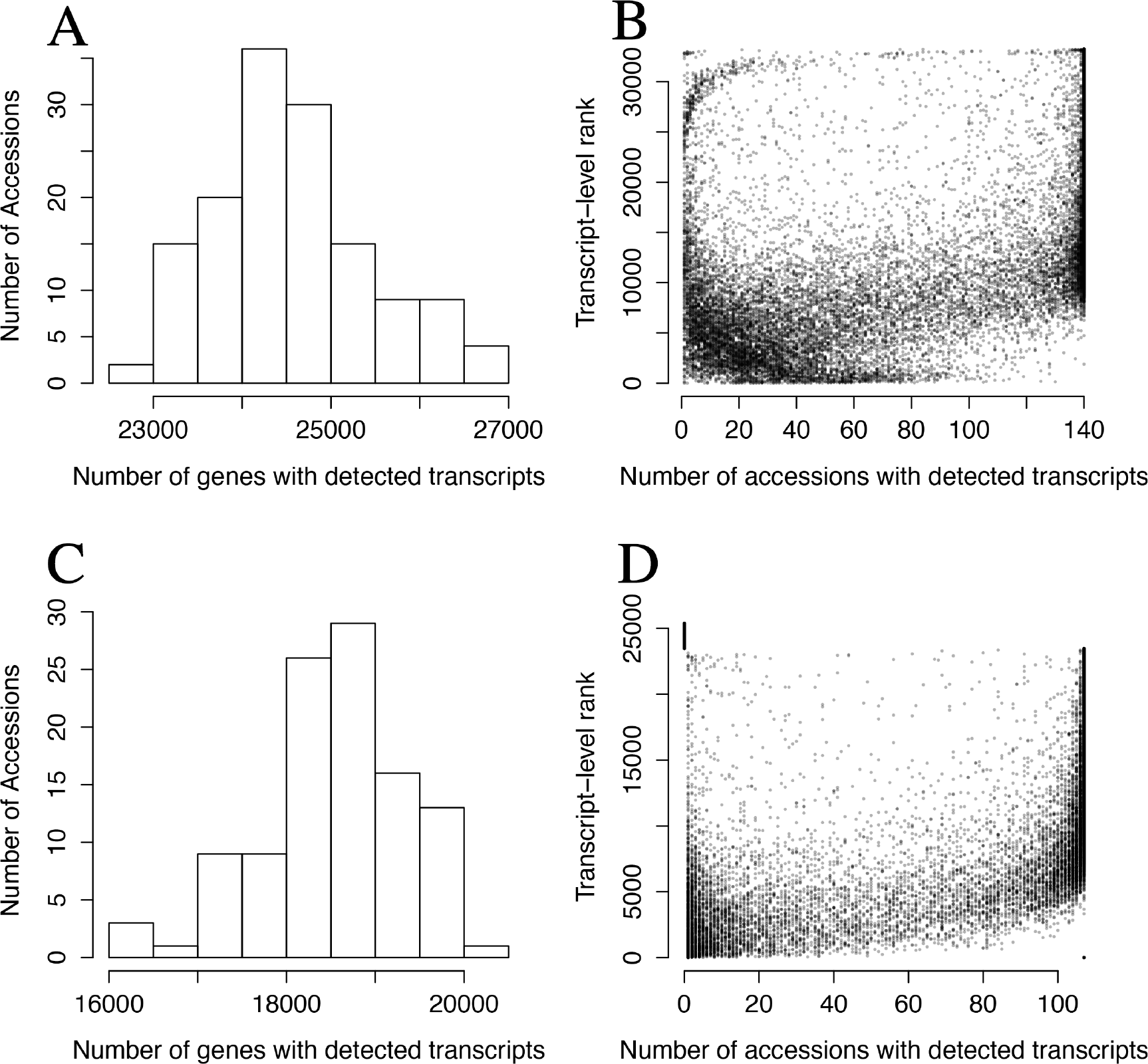
**(A)** Distribution of the number of genes with transcripts in the leaf of 140 natural A. thaliana accessions [32] scored by RNA-seq (Normalized FPKM > 0). **(B)** Relationship between the ranks of the average transcript levels for all genes with transcripts detected in at least one accession (y-axis) and the number of accessions in [32] where transcripts for the gene is found (x-axis). Each dot in the plot represent one of the 33,265 genes with FPKM >0 in at least one accession of [32]. The transcript-level rank is based on average transcript levels in the accessions where transcripts for a particular gene are detected. Due to this, the ranks are less precise for transcripts present in fewer accessions. **(C)** Distribution of the number of genes with detected transcripts in the leaf of 107 Swedish natural A. thaliana accessions [33] scored by RNA-sequencing (RPKM > 0). **(D)** Relationship between the ranks of the average transcript levels for all genes with detected transcripts in at least one accession and the number of accessions in [33] where the gene is expressed. Each dot in the plot represent one of the 25,382 genes with RPKM > 0 in at least one accession of [33].

### *RNA-seq bias likely explains part of the variability in detected transcripts of the individual* A. thaliana *accessions for lowly expressed genes*

RNA-seq detects transcripts from nearly all genes in the genome in at least one of the accessions in the *SCHMITZ-data*. In addition to a core set of genes with transcripts in all accessions, there is a large set of genes (on average 6,256 per accession) for which transcripts are detected only in some of the accessions. A possible explanation for this might be that RNA-seq is unable to reliably detect low levels of transcripts in the samples. Such RNA-seq introduced bias would result in a random detection of transcripts for individual accessions (i.e. produce a score = 0) for lowly expressed genes, and lead to a similar variability in the presence or absence of transcripts among the accessions as when there is a true accession-specific expression. To evaluate the potential contribution of RNA-seq bias to the results, we first evaluated the relationship between the rank of the transcript-levels across the genes and the number of accessions in which transcripts were detected (Figure 2B). There was a clear overall trend in the data that transcripts were detected in fewer accessions for genes with lower overall transcript-levels. A similar trend was observed also in the *DUBIN-data* (Figure 2D). Based on this, we conclude that RNA-seq bias contributes to the observed variation in presence or absence of transcripts among the accessions. However, transcripts were missing in a large fraction of the studied accessions also for many of the genes with high rank/overall transcript level. As RNA-seq bias is an unlikely explanation in these cases, it suggests that at least part of the variability in the presence or absence of transcripts might be due to accession-specific expression, or structural variations leading to different sets of genes in divergent accessions [16]. Thus, our overall results agree across the two datasets in that i) the number of genes expressed in the leaf of an individual plant is large, and ii) that the actual set of genes that are present or expressed varies between the individual accessions.

### *Genetics contributes to the variability in detected transcripts of the individual* A. thaliana *accessions*

The observed variability in the RNA-seq scored presence or absence of transcripts for individual genes between accessions could also be caused by experiment specific environmental factors affecting e.g. plant growth or sample treatments. As transcript variability due to such effects on plants and samples are expected to be experiment-specific, they should not replicate across experiments. We therefore used the data from a second publicly available RNA-seq dataset from 107 natural Swedish accessions, with fully sequenced genomes and transcriptomes *(DUBIN-data;* [33]), to explore whether a similar pattern of accession-specific transcripts was present also in this data among the genes with high transcript levels. In the DUBIN-data, the number of genes with detected transcripts were lower both when measured within any accession (23,478 with RPKM - Reads Per Kilobase per Million mapped reads - > 0), or within the individual accessions (16,136 to 20,109 with an average of 18,663; Figure 2C). The core set of genes expressed in all accessions contained 12,927 genes. The lower number of genes with transcripts is likely a result of the lower sequencing depth (˜1/4 of the *SCHMITZ-data)* and the more stringent filtering of the reads. However, the overall trend in the results is the same as in the *SCHMITZ-data:* genes with lower transcript-levels had transcripts detected in fewer accessions, but transcripts for many genes with high overall transcript levels were also found only in a few accessions (Figure 2D). This overlap between the results from the two datasets suggests that the variation in which transcripts are highly expressed in the leaf of several, but not all, accessions is unlikely to be due to either technical bias or experimental specific errors as described above.

To explore whether there was a genetic basis for the presence or absence of transcripts for individual genes, we studied a set of 4,317 genes in the *SCHMITZ-data* in more detail. These genes were selected to i) have detected transcripts in more than 14 (10%), but fewer than 126 (90%), of the accessions and ii) have transcript levels that were higher than the 2^nd^ lowest expressed gene with transcripts in all accessions (Materials and Methods; Figure 2B). This subset was chosen to remove most of the genes influenced by the RNA-seq bias for lowly expressed genes (as discussed above). To estimate the genetic basis for the transcriptome variation, we first created a covariance matrix for the proportion of transcript sharing among all pairs of accessions based on the average number of shared genes with detected transcripts (FPKM > 0). Then, we created a covariance matrix for the relationship between the accessions based on the genome-wide genotype as a genetic kinship matrix weighted by allele frequencies based on all SNPs with MAF > 0.05. As illustrated in Figure 3, there was a highly significant correlation between the genetic and transcriptome covariances for this set of genes (r = 0.035, p = 1.0 × 10^-16^; Figure 3). Although the correlation is low, it indicates that plants that are genetically identical (i.e. the same accession), on average share 4,317 × 0.035 = 151 transcripts more than they would with a genetically unrelated accession. This shows that there is a significant genetic contribution to the sharing of transcripts between accessions, suggesting that the accession-specificity of the transcripts has a genetic basis.

**Figure 3.**
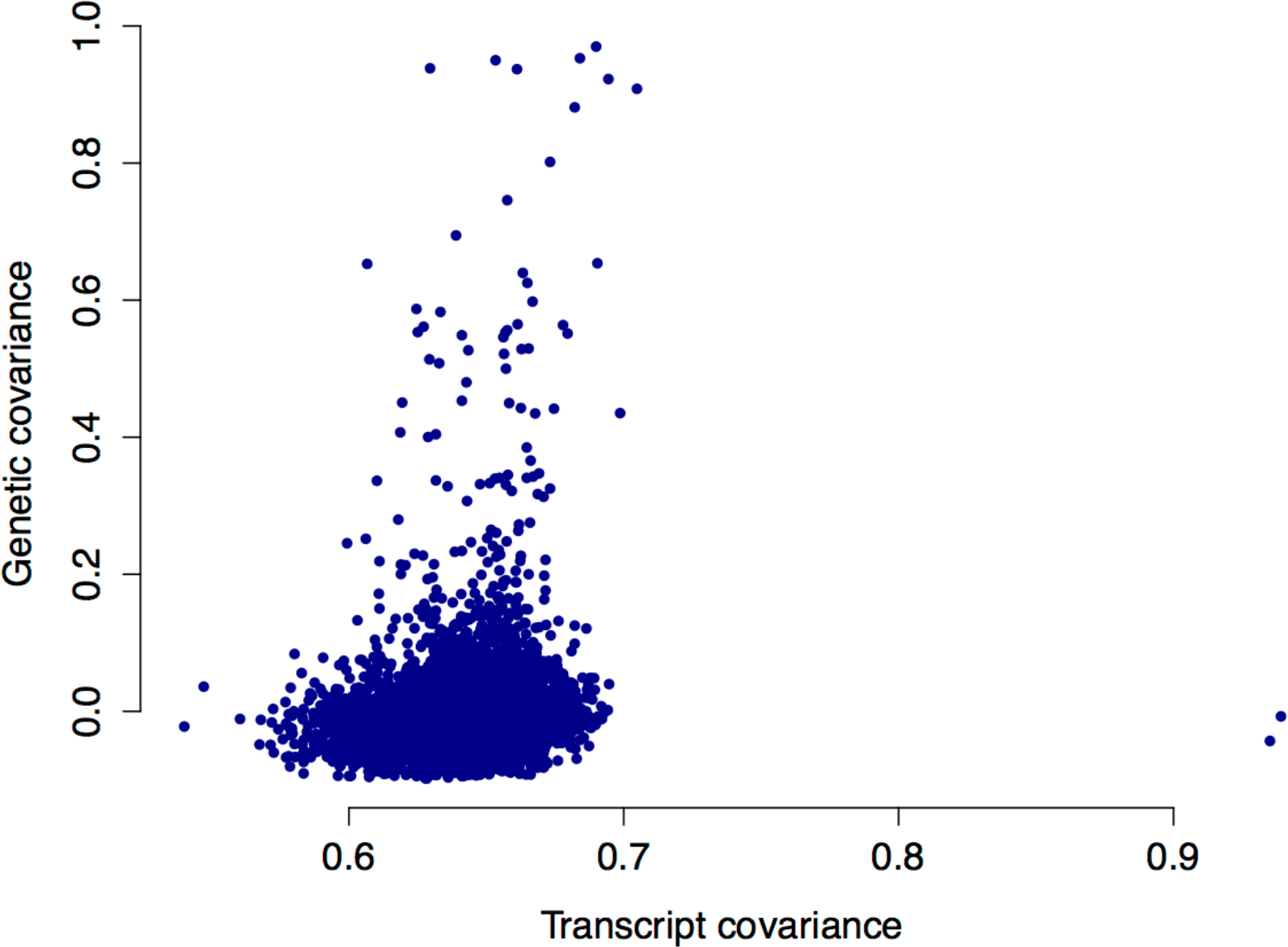
Correlation between the genetic and transcriptome covariances among 4,317 genes with transcripts detected in between 14 and 126 of the accessions in the SCHMITZ-data and that are expressed above a level where transcripts RNA-seq have been able to detect transcripts for a gene in all accessions. Each dot in the figure represents a pairwise relationship between two accessions, with the transcript covariance on the y-axis and the genetic covariance on the y-axis.

### cis-eQTL contribute to the accession-specific patterns in the transcriptome

Loss-of-function alleles are known to be important contributors to natural trait-variation in *A. thaliana* [5, 7, 8, 11, 31, 34-38, 40]. Earlier works on the genetic regulation of expression-variation in an *A. thaliana* have also shown that cis-eQTL have a larger effect on expression than trans-eQTL [23, 29]. It is also known that many such ELP genes carry deletions in the regulatory region [20], but in some cases the transcriptional variation is also due to large structural variations leading to, for example, loss of entire genes [16]. To maximize eQTL-mapping power, we therefore screened the 4,317 genes that are most likely to have a genetically controlled accession-specific expression in the *SCHMITZ-data* (i.e. that are the least likely to be affected by RNA-seq bias as discussed above) for cis-eQTL alleles affecting the presence or absence of transcripts in the accessions. A customized eQTL-mapping approach designed for this scenario was developed and used for this analysis (see Materials and Methods for details).

For each of the 4,317 genes, we first binarized (0/1) the phenotype to indicate presence (1; Normalized FPKM > 0) or absence (0; Normalized FPKM = 0) of transcripts for the assayed gene in each accession. Then, we performed an association analysis to all SNP markers in a 1 Mb region around the evaluated gene. For each SNP marker in this region, a logistic regression was fitted to the genotype of the marker with the binarized phenotype as response. Significant associations are interpreted as detection of a cis-eQTL contributing to the presence or absence of transcripts for the studied gene. In total, 349 such cis-eQTLs were detected (FDR=12.3%). For 172 of the 349 genes, whose transcript-levels were affected by these cis-eQTL, transcripts were also detected in some, but not all, of the accessions of the *DUBIN-data* [33]. By performing the same cis-eQTL analysis for the presence or absence of transcripts for these 172 genes, 81 of the cis-eQTL (FDR = 0.1) could be replicated (Table S2). Given that the collections of accessions in the *SCHMITZ-* and *DUBIN-data* were obtained from non-overlapping geographical locations, it is striking that so many of the cis-eQTL with the ability of almost shutting off the expression are present in, and could be replicated across, such diverse datasets.

### Mapping of eQTL for genes with transcripts in most accessions

The majority of the genes in the *SCHMITZ-data* (20,610) [32] were expressed in more than 90% of the accessions. The transcript-levels (Normalized FPKM) for these genes were quantile-transformed and used as phenotypes in linear mixed model based, kinship corrected genome-wide eQTL scans. In total, these analyses revealed 2,320 eQTL (FDR = 0.09), with 1,844 (79.5%) of the associations in cis (within a ±1 Mb window around the mapped gene) and 476 (20.5%) eQTL in trans affecting the expression of 2,240 genes (Table S4). All eQTLs were significant after correction for genome-wide analyses across multiple expression traits. Out of the 2,320 genes with eQTL in the *SCHMITZ-data,* 2,006 had transcripts in all accessions of the *DUBIN-data* [33], and 649 of the eQTL affecting 636 genes could be replicated at a 0.01 Bonferroni threshold correcting for the number of tested markers in the replication region (FDR = 0.031; Table S5). Among the 2,240 genes with eQTL in the *SCHMITZ-data,* 175 have earlier been shown to have an altered phenotype from loss-of-function alleles [41] and 38 of these were among those replicated in the DUBIN-data. As many of these genes have strong effects on potentially adaptive traits, including development, hormone pathways and stress responses, they are plausible functional candidate adaptive genes in *A. thaliana.* These genes are listed in Table S6.

### Significant contribution by common structural variants to many of the eQTL

Structural variations, including regulatory unit deletions [20] and larger genome rearrangements [16] have been found to contribute to the transcriptional variation in natural *A. thaliana* population both for individual genes [2, 16, 42, 43] and on a whole transcriptome level [16, 20]. We evaluated the contribution of common structural variations to the presence or absence of transcripts in individual accessions by quantifying the overlap of mapped reads in the genome and transcriptome sequencing to the 646 detected eQTL genes that completely lacked transcripts across the gene body of at least one accession. Using the genome re-sequencing data, we first identified a set of putative structural variants by identifying the individual genes and accessions where no reads were mapped to either the transcription start site (TSS) or the entire gene body of these eQTL genes. Since the probability of observing a lack of mapped reads at the exact same region in multiple accessions by chance is very low, we compiled a high-confidence set of common structural variants by retaining only those where reads were lacking in the TSS or gene body of the same gene in more than 5 accessions. In total, 155 of the 349 cis-eQTL genes showed this pattern. On average, each accession carried 58 genes with such structural variations and each gene was disrupted in 52 accessions (Figure S2). To quantify the contribution of these structural variants to the expression variation observed in the RNA-seq analysis, we tested for association between the presence/absence of the structural variation and the presence/absence of RNA-seq reads in the accessions using a Fisher exact test. This association was significant on a nominal level (P < 0.05) for 122 of these genes, and 94 of them passed a significance-threshold that was Bonferroni corrected for testing 155 genes. This suggests that the accession-specific transcript pattern observed is often due to structural variations (Table S2) and our eQTL approach is efficient in detecting such segregating variants. In the standard eQTL analysis that was used to map genes that had transcripts in >90% of the accessions, 297 genes lacked transcripts in at least one accession. By conducting the same structural-variant analysis for these genes, we identified an additional 93 genes with common high-confidence structural variations. In total, 37 of these were significantly associated with the transcript-levels in the accessions on a nominal level (P < 0.05) and 17 passed a multiple-testing corrected significance threshold. Thus, in total 248 genes were found to carry at least one common structural variation and for 159 of these, the variants were associated with the transcript-levels in the accessions. Of the 248 genes with common structural variations, 85 contained at least two common structural variants and 60 of the genes with two variants were significantly associated with the presence/absence of transcripts in the accessions. Our results show that structural variations are common, and often multi-allelic, in natural *A. thaliana* accessions and make important contributions to the observed transcriptome variation.

### Identification of functional candidate genes with cis-eQTL contributing to the accession specific presence or absence of transcripts

12 of the 349 genes affected by the cis-eQTL mapped in our study had earlier been subjected to functional studies in which the gene had a distinct phenotypic effect (Table 1). The cis-eQTL-mapping result for one of these genes (*AT2G21045*; High Arsenic Content 1; *HAC1)* is illustrated in Figure 4. *HAC1* has a skewed distribution of RNA-seq scored transcript levels (Figure 4A/4C) and a highly significant cis-eQTL signal in both the *SCHMITZ* and *DUBIN-data* (Figure 2B/2D; p = 1.75 × 10^-10^/ p = 9.32 × 10”^11^, respectively). It has been shown that loss of *HAC1* expression increases the amount of Arsenic in the leaf and that several functional polymorphisms might be present in natural *Arabidopsis thaliana* population [3][4]. Here, we detected a cis-eQTL for the expression of *HAC1* with the top SNP located within an exon of *HAC1* (Figure 4B; top SNP Chromosome 2 at 9,028,685bp) in the *SCHMITZ-data.* This signal was replicated with a high significance also in the *DUBIN-data* (Figure 4D). This gene is thus a highly interesting functional candidate adaptive gene as i) it has earlier confirmed effects on Arsenic-levels in the plant, ii) there is a strong cis-eQTL regulating presence or absence of transcript for the gene segregating in natural *A. thaliana* accessions, and iii) effect of the polymorphisms could be replicated at high significance in both analysed populations. Further experimental studies are, however, needed to functionally replicate the effect of the remaining 11 polymorphisms on expression and the indicated phenotype. This to clarify whether such a link, together with the segregation in the worldwide *A. thaliana* population, means that it has contributed to adaptation.

**Figure 4.**
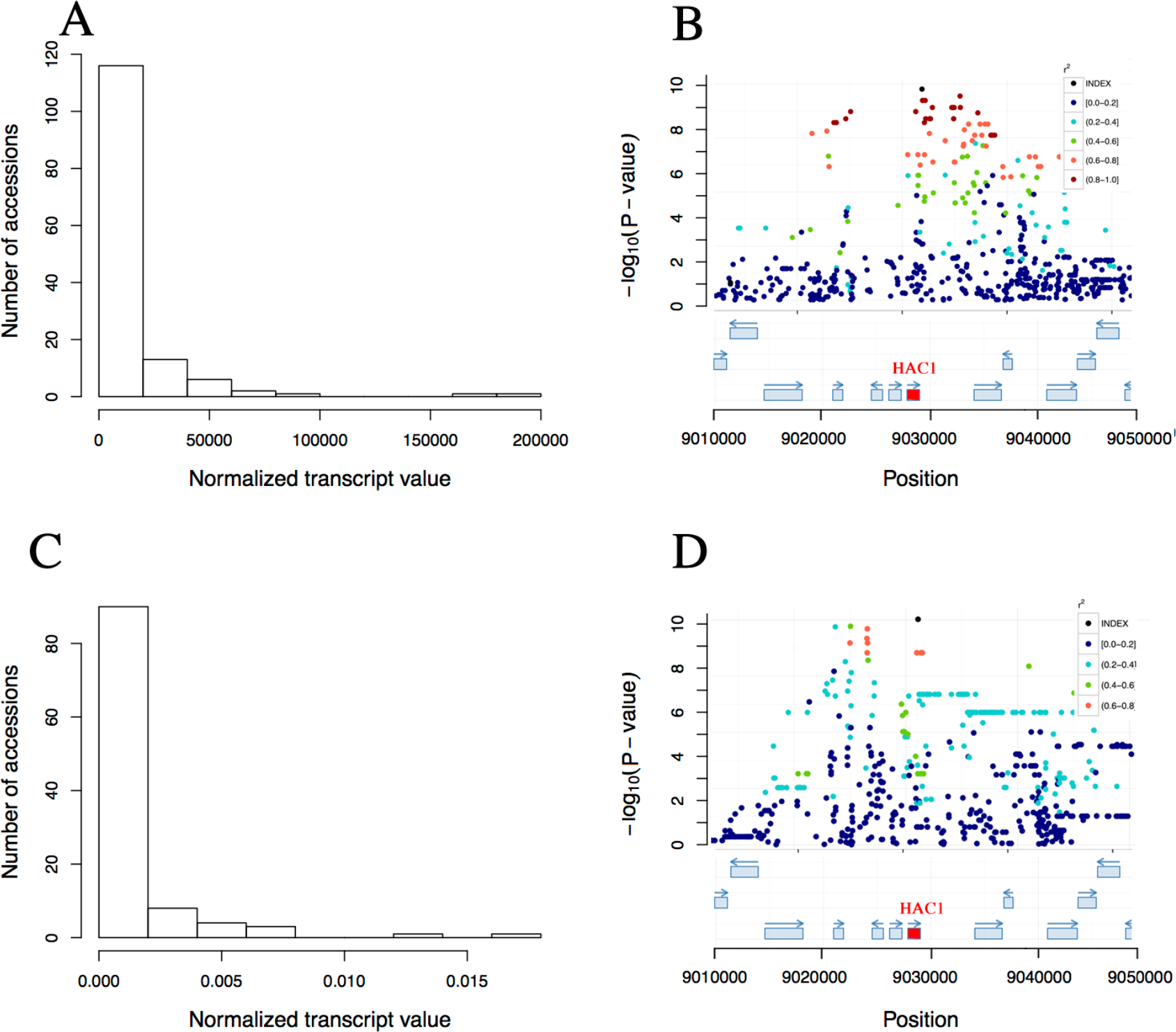
The eQTL analysis detects a highly significant, replicable association for the expression of the gene HAC1 (AT2G21045). The peak SNP is located in an exon of HAC1. (A/C) Distributions of transcript-levels for the 140/107 accessions (FPKM/RPKM-values from RNA-sequencing) in the SCHMITZ-data (A) and DUBIN-data (C) [32, 33], respectively. (B/D) Illustrations of the association-profiles [55] for expression of the gene AT2G21045 (HAC1) in the SCHMITZ-data (B) and the DUBIN-data (D) ([32]/[33]), respectively. There is a highly significant cis-eQTL to a SNP located in an exon of HAC1.

**Table 1.** Genes with cis-eQTL detected in the population of 140 natural A. thaliana accessions (SCHMITZ-data) contributing to the accession specific presence or absence of transcripts and earlier reported biological function.

| *Locus* | *Gene* | *<sup>a</sup>SNP* | *<sup>b</sup>MAF* | *<sup>c</sup>Log odds ratio±SE* | *<sup>d</sup>P-value* | *<sup>e</sup>Replicated* | *Reference* |
| --- | --- | --- | --- | --- | --- | --- | --- |
| *AT2G21045* | *HAC1* | chr2_9028685 | 0.14 | 4.84±0.76 | 1.75 x 10<sup>-10</sup> | Yes-GBF | [3] |
| *AT5G59340* | *WOX2* | chr5_23935224 | 0.43 | 2.82±0.54 | 1.83 x 10<sup>-7</sup> | Yes | [44] |
| *AT1G77235* | *MIR402* | chr1_29007464 | 0.46 | 7.07±1.02 | 5.18 x 10<sup>-12</sup> | No.e | [45] |
| *AT2G38220* | *APD3* | chr2_16017043 | 0.29 | 0.37±0.07 | 3.64 x 10<sup>-7</sup> | No.e | [46] |
| *AT1G13430* | *ATST4C* | chr1_4601762 | 0.51 | 0.36±0.06 | 3.87 x 10<sup>-10</sup> | No | [47] |
| *AT1G66600* | *WRKY63* | chr1_24217798 | 0.06 | 0.14±0.02 | 3.22 x 10<sup>-10</sup> | No | [48] |
| *AT2G19500* | *ATCKX2* | chr2_8168512 | 0.07 | 0.18±0.03 | 6.13 x 10<sup>-9</sup> | No | [49] |
| *AT3G27620* | *AOX1C* | chr3_10230473 | 0.39 | 0.4±0.07 | 6.45 x 10<sup>-8</sup> | No | [50] |
| *AT3G57270* | *BG1* | chr3_22031771 | 0.06 | 0.15±0.02 | 1.09 x 10<sup>-9</sup> | No | [51] |
| *AT4G01420* | *CBL5* | chr4_583422 | 0.12 | 0.2±0.03 | 7.48 x 10<sup>-12</sup> | No | [52] |
| *AT4G02850* | *DAAR1* | chr4_597373 | 0.06 | 0.15±0.02 | 1.41 x 10<sup>-9</sup> | No | [53] |
| *AT4G29340* | *PRF4* | chr4_14639984 | 0.21 | 0.36±0.07 | 2.32 x 10<sup>-7</sup> | No | [54] |

<sup>a</sup>SNP: Top SNP in association analysis; <sup>b</sup>MAF: Minor allele frequency of the top associated SNP in the *SCHMITZ-data*; <sup>c</sup>Log odds ratio of the top associated SNP ± Standard Error; <sup>d</sup>P-value: Nominal p-value for the top associated SNP; <sup>e</sup>Replicated: Could the original association in the *SCHMITZ-data* be replicated in the *DUBIN-data*, Yes-GBF: Replicated at Genome Wide Bonferroni threshold, Yes, replicated at Bonferroni threshold correcting for number of markers in ± 1 Mb window of the peak SNP in *Dubin-data* No.e: not expressed in *DUBIN-data* so could not be tested for replication. No: Non-significant in replication analysis.

## Discussion

*Arabidopsis thaliana* is a small flowering plant that has colonized a wide range of habitats, and while adapting to these it has evolved a remarkable genotypic and phenotypic variation in the world-wide population [1, 2]. Genetic regulation of gene expression is known to be an important mechanism underlying adaptation [38, 56-58] and many studies have been conducted to investigate the gene expression variation in natural *A. thaliana* accessions [16-18, 20, 21, 23, 39]. These earlier studies have provided valuable insights to the general characteristics of expression variation in *A. thaliana.* Here, we extend these studies by exploring the whole transcriptome in a considerably larger set of natural accessions from the worldwide population.

We have studied the transcript variation in the leaf of natural *A. thaliana* accessions by analyzing data from two large sets of natural accessions from the global *A. thaliana* population. We find that the transcript variation correlates with the genome divergence between the accessions and map thousands of individual eQTL that contribute to the variation in leaf transcript-levels across the accessions. These extensive and genetically controlled differences in expression-levels controlled by alleles that segregate in the wild are a useful resource to explore how selection might have acted on different regulatory variants of genes important for the adaptation of *A. thaliana* to diverse living conditions. Several functional candidate adaptive genes were identified among the eQTL, for which further experimental validation would be highly motivated.

We found that the average proportion of genes with detected transcripts in the leaf within the individual accessions was slightly higher in this study, than in earlier studies based on smaller numbers of accessions (73% here vs 64% in [39]). In the seedling, Gan *et al.* [16] report a slightly higher estimate for protein-coding genes, but lower if all genes are considered. A large core-set of genes (61% of all with scored transcripts) was detected with transcripts in nearly all (>90%) of the accessions. The genes in this core-set overlapped to a great extent between the two independent collections of natural *A. thaliana* accessions studied here, suggesting that these genes include both basal, and leaf-specifically, expressed genes.

When analysing the deep-coverage RNA-seq from the largest collection of natural accessions, we detected transcripts in the leaf from nearly all genes in the genome in at least one of the accessions. Further, a large number of apparently accession-specific transcripts were also detected due to high sensitivity of the RNA-seq approach. The majority of the accession-specific transcripts were found at low levels in only a few accessions, making it difficult to separate true accession specific expression from bias in the RNA-seq analysis. For example, some genes are known to vary their transcripts levels in a circadian manner (2% according to [59] or 6% according to [60]), which might reduce the expression of these genes to a lower level in all accessions and lead to random detection in RNA sequencing. However, transcripts from a relatively large number of genes were abundant in some accessions, and completely absent in others. This suggests that these genes are more likely to be expressed, or lost, in an accession-specific manner, which is supported by the highly significant correlation between the genetic and transcriptome covariances amongst the accessions.

Here, we developed an approach to map cis-eQTL affecting the presence or absence of transcripts in the accessions motivated by earlier findings that illustrated the importance of strong loss-of-function alleles for adaptation in *A. thaliana* [5, 7, 8, 11, 31, 34, 35, 37, 38]. Given that phenotypes of many mutant alleles are already described in the literature, our aim was to identify new functional candidate adaptive genes by screening for cis-eQTL where alleles contributing to the presence or absence of transcripts for the studied genes segregated in populations of natural *A. thaliana* accessions. If these cis-eQTL affect a gene with a known phenotype, it can then be considered as a strong candidate adaptive gene. Among our results, we find a particularly interesting example of where our approach is able to detect adaptive genes that have been studied in detail. *HAC1* [3, 4] was detected by our approach in the larger collection [32] and replicated in the smaller [33], and its role in the reduction of Arsenik has earlier been described in the literature. We find this to be a useful proof-of-principle finding suggesting that at least some of the highlighted functional candidate adaptive genes might be found important for adaption in future studies.

Earlier studies have shown that utilization of several natural *A. thaliana* accessions, in addition to Col-0, is useful for uncovering biologically important genetic and phenotypic variations that are not present in this reference accession [5, 7, 8, 11, 16, 31, 34-36]. Our results further support this as 5% (111) of the cis-eQTL genes lacked transcripts in the leaf of *Col-0.* The fact that half of them (53) also lacked transcripts in *Col-0* seedling and pollen [61](Table S7) suggests that these genes are not expressed at all in this accession [16].

Structural variations have earlier been found to be common in the genome and make a significant contribution to the transcriptional variation in *A. thaliana* genes (see for example [2, 16, 20, 36, 42, 43]). We found that 248 of the 646 genes that lacked transcripts in one or more accessions, and for which eQTL were mapped, also lacked reads that covered the transcription start site, or the entire gene, in the available whole-genome sequencing data. For about one third of the genes (80), both types of structural variants were common. For a majority (159) of these genes, the structural variants were significantly associated with the presence or absence of the transcript. This result confirms that structural variations regulated transcript variation are relatively common in *A. thaliana* [16, 20], that they are often multi-allelic, and are likely to make significant contributions to the transcriptome variation in the worldwide population.

When there is a leap in technology, it is important to adapt the existing analytical methods to better utilize the additional information. In our particular case, we make use of the information about expression-traits where the transcript distributions for the assayed genes are heavily zero-inflated (i.e. contain many accessions with no detectable transcripts). We show that these distributions, in many cases, are influenced by cis-eQTL with strong effects on the transcript levels. These distributions are too skewed to be appropriately transformed or modelled using other distributions such as a negative binomial [62]. Earlier eQTL-mapping studies based on RNA-seq data have therefore either removed genes whose transcript levels were zero-inflated [63], utilized non parametric testing to avoid the assumption of normality [64] or utilized regression without first addressing the potential issues arising from the non-normality of the transcript-levels [65]. Our approach to, based on the observed distribution properties, propose and test for a particular type of genetic effects causing accession-specific expression, we could reveal many new cis-eQTL that in many cases are likely to be due to structural variants that either delete important regulatory regions for the genes, or the entire gene. Many of these associations replicated in an independent dataset and several genes were promising functional candidate adaptive genes. Although the proposed cis-eQTL-mapping approach is not a general framework for mapping all eQTL, it illustrates the value of developing and utilizing analysis methods for genome and transcriptome data to test well-defined biological hypotheses.

## Materials and methods

### *Whole genome re-sequencing and RNA-seq data for a population of 144 natural* A. thaliana *accessions*

In an earlier study, Schmitz *et al.* [32] RNA-sequenced a collection of 144 natural *Arabidopsis thaliana* accessions. We downloaded this data together with their corresponding whole-genome SNP genotypes made available as part of the 1001 Genomes project [2]. According to the author’s description and the original publication [32], the plants were grown in 22°C and leaves had been collected from rosettes prior to flowering. Further, RNA reads had been aligned to SNP-substituted reference genomes for each accession using Bioscope version 1.3 with default parameters. Cufflinks version 1.1 had been used to quantify Transcript levels using the following parameters: ‘-F’ 0; ‘-b’; ‘-N’; ‘-library-type’ fr-secondstrand; ‘-G’ TAIR10.gtf. Raw FPKM values were quantile normalized by the 75th percentile and multiplied by 1,000,000 to be transformed to a comfortable scale. We removed two accessions from the data (Alst_1 and Ws_2) due to missing genotype data and two accessions (Ann_1 and Got_7) due to their low transcript Call Rate (16,861 and 18,693 genes with transcripts as compared to the range of 22,574 to 26,967 for the other the accessions). The final dataset used for eQTL mapping *(SCHMITZ-data)* included 1,347,036 SNPs with MAF above 0.05, a call-rate above 0.95 and RNA-seq derived FPKM-values for 33,554 genes. For identifying regions without mapped reads in the genome re-sequencing, we used the information about this that was available on the SALK webpage (http://signal.salk.edu/atg1001/download.php).

### *Whole genome re-sequencing and RNA-seq data for a population of 107 Swedish natural* A. thaliana *accessions*

We downloaded a second public dataset of 107 Swedish *A. thaliana* lines with fully sequenced genomes and transcriptomes [33] from plants grown in 10°C and 16°C. Here, RNA had been prepared from whole rosettes collected at the 9-true-leaf stage. RNA reads had been aligned using PALMapper aligner using a variant-aware alignment strategy. Reads that were longer than 24 bp and uniquely mapped into the exonic regions had been used to quantify expression. Further details about reads filtering and transcripts quantification can be found in [33]. We used the same quality control procedures for this dataset as for the larger dataset described above. We compared the data from the experiments done at 10 C and 16 C and found that they were quantitatively similar and therefore only used the data generated at 10°C *(DUBIN-data)* for further analysis. In total, this data contained 1,785,214 SNPs with MAF above 0.05, a call-rate above 0.95, and RNA-seq derived RPKM-values for 33,322 genes.

### Cis-eQTL mapping to detect polymorphisms contributing to the accession specific presence of gene-transcripts

First, we selected the 4,317 genes in the *SCHMITZ-data* that i) had transcripts in more than 14 (10%), but fewer than 126 (90%), of the accessions and ii) had transcript-levels higher than the 2^nd^ lowest expressed gene with transcripts in all accessions. Then, we binarized the transcript-level phenotype by assigning values zero or one for each accession depending on whether transcripts for the gene were present (Normalized FPKM > 0) or not (Normalized FPKM < 0). Then, a logistic regression approach was used to perform an analysis across the SNP markers in a ±1Mb region around each of the tested genes using the *qtsore (family = binomial)* function in GenABEL [66].

For significance testing, we first applied a Bonferroni corrected significance threshold correcting for the number of SNPs tested in the ±1Mb region around the gene. Second, we accounted for the potential noise in the transcript measurements by applying a second filtering based on a permutation test performed as follows. Under the assumption that all the 0 values (lack of transcripts) resulted from non-biological noise, the binarized phenotypes were randomly shuffled with respect to the cis-SNP genotypes 1,000 times and an association scan performed in each of these datasets as described above. In each scan, the minimum p-value was saved to provide an empirical null-distribution for every trait. Trait-specific significance-thresholds were obtained by taking the 1% cut off from these distributions. To account for multiple testing across all tested traits (genes), the FDR was calculated as the expected number of significant traits with eQTL at the given significance-threshold under the null hypothesis, divided by the number of traits where significant eQTL were detected.

### Mapping of eQTLs for genes with transcripts in most accessions

For the 20,610 genes in the *SCHMITZ-data* where transcripts were detected in more than 90% of the accessions, a standard eQTL mapping approach was used by performing a GWA analysis fitting inverse-Gaussian transformed expression values to the genome-wide SNP genotypes in a linear mixed model (1) using the *polygenic* and *mmscore* functions in GenABEL package [66].

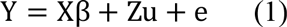

Y is here the transformed expression phenotype, which is normally distributed with mean 0. *X* is the design matrix with one column containing the allele-count of the tested SNP (0,1,2 for minor-allele homozygous, heterozygous and major-allele homozygous genotypes, respectively). *β* is a vector of the additive allele-substitution effect for the SNP. *Z* is the design matrix obtained from a Cholesky decomposition of the kinship matrix *G* estimated from the whole-genome, MAF-filtered SNP data with the *ibs* function (option weight = ‘freq’) in the GenABEL package [66]. The Z matrix satisfies ZZ’ *=* G, and thus, the random effect vector *u* is normally distributed, 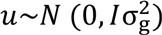is the normally distributed residual variance with 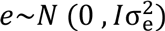The GWA results were visualized using the *cgmisc* R-package [55].

A permutation test was used to set the significance threshold in the analysis [67, 68]. As the number of traits was too large to derive a trait-specific threshold, a GWA-scan was performed across all traits to identify those with at least one SNP with p < 1 × 10^-6^. Among the genes that passed this threshold, a random set of 200 traits was selected. For each of these traits, GWA were performed in two hundred permuted datasets where GRAMMAR+ transformed residuals [7, 69] were used as phenotype, resulting in 40,000 permutations in total. Based on this total distribution, we derived a 1% significance threshold (-logi_0_(p-value) = 6.84).

### Replication of detected eQTLs

As 26/38% the SNPs in the *SCHMITZ-data* were missing from the *DUBIN-data* (before/after filtering for MAF), we replicated our findings by testing for associations to SNPs in the DUBIN-data that were physically close to the detected top SNP in the SCHMITZ-data. The average expected LD-block size in Arabidopsis is about 10 kb [70], and therefore we performed the replication analysis to SNPs located in a ±10 kb region around the eQTL peaks in the *SCHMITZ-data* and associate their genotypes to the transcript level of the corresponding genes in the *DUBIN-data*. We considered a cis-eQTL replicated if i) the significance for any of the SNPs tested in this region passed a significance threshold corrected for the number of tested SNPs in this region using Bonferroni correction, and ii) the overlapping SNPs show effects in the same direction. FDR was calculated for the results across the trait as the expected number of false positives (0.05 ⋆ the number of tested regions) divided by the number of regions with significant eQTL.

### Calculating covariances between transcriptome variation and genome variation

Based on the binarized expression values of the 4,317 selected genes (selection procedure described above), we created a relationship matrix following the same approach as when calculating the genomic IBS matrix (see details above). The correlation between the transcriptome and genome variation was calculated as the correlation between elements in these two relationship matrixes.

### Definition of cis and trans eQTL peaks

For each gene with a significant GWA, the SNP with the lowest P-value was selected as the peak location for the eQTL. When the leading SNPs had been defined for each trait, SNPs were considered to represent the same eQTL if they were located within 1 Mb of each other in the TAIR10 reference genome. eQTL peaks located within 1Mb up-or downstream of the genes whose expression was used as phenotype were classified as cis-eQTL, while the remaining eQTL peaks were classified as trans-eQTL.

### Supplementary Material

**Table S1.** Illustration of the geographical locations for the 140 natural Arabidopsis thaliana accessions utilized in the eQTL analyses.

| *Locus* | *Annotation* |
| --- | --- |
| AT1G11640 | tRNA-Trp |
| AT1G20210 | tRNA-Asp |
| AT1G22651 | Unknown |
| AT1G28740 | tRNA-Pro |
| AT1G28750 | tRNA-Pro |
| AT1G28790 | tRNA-Pro |
| AT1G28810 | tRNA-Pro |
| AT1G28830 | tRNA-Pro |
| AT1G28840 | tRNA-Pro |
| AT1G28910 | tRNA-Pro |
| AT1G28990 | tRNA-Pro |
| AT1G57710 | tRNA-Undet |
| AT1G80831 | Unknown |
| AT2G01667 | Unknown |
| AT2G16016 | Unknown |
| AT2G16881 | Unknown |
| AT2G34202 | MIR399D |
| AT2G40802 | Unknown |
| AT3G04732 | Unknown |
| AT3G06335 | tRNA-Pro |
| AT3G29195 | Unknown |
| AT3G29633 | Unknown |
| AT3G49051 | Unknown |
| AT3G50651 | Unknown |
| AT5G41774 | Unknown |
| AT5G65445 | tRNA-Ser |
| AT5G66211 | Unknown |

**Table S2.**
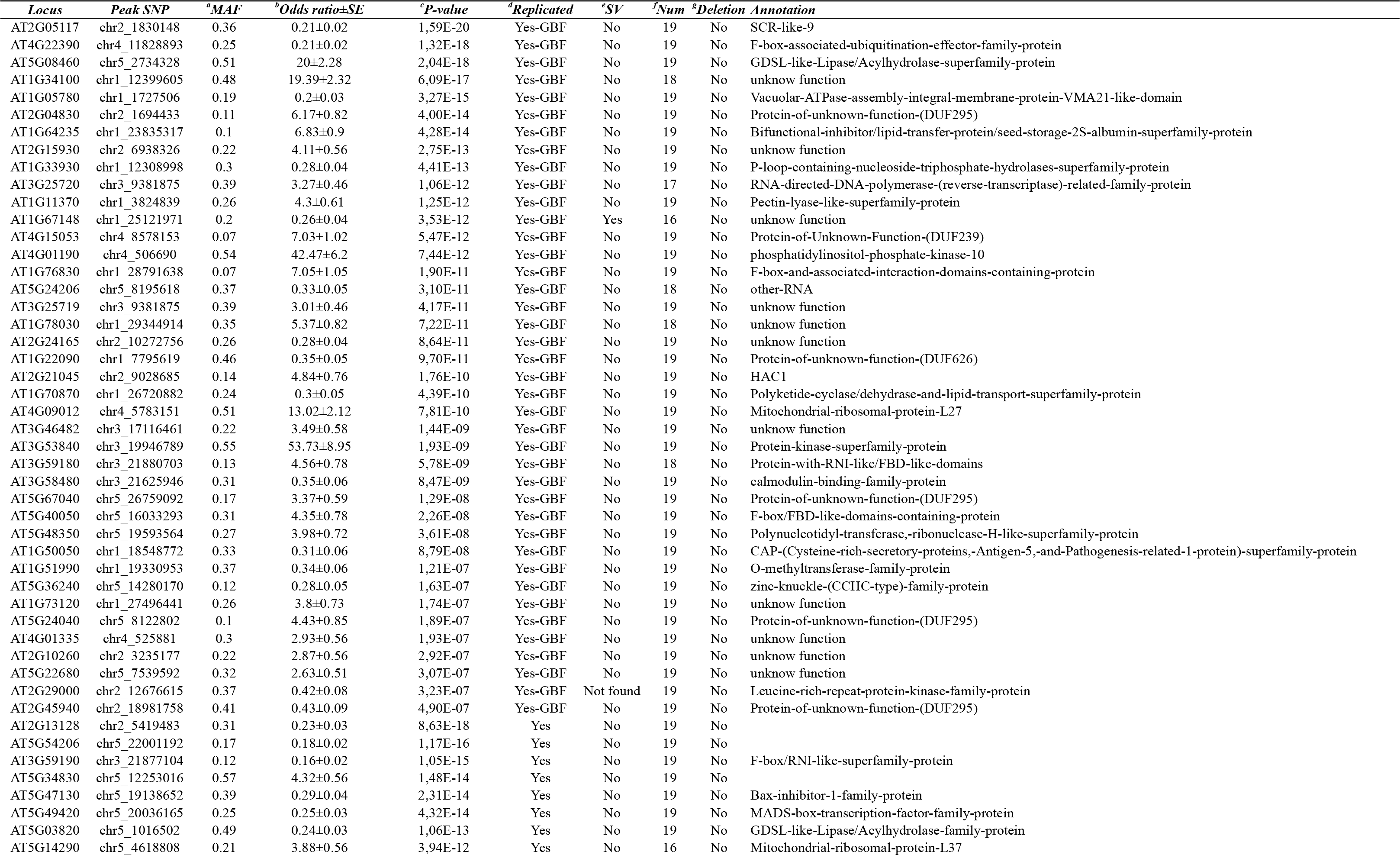
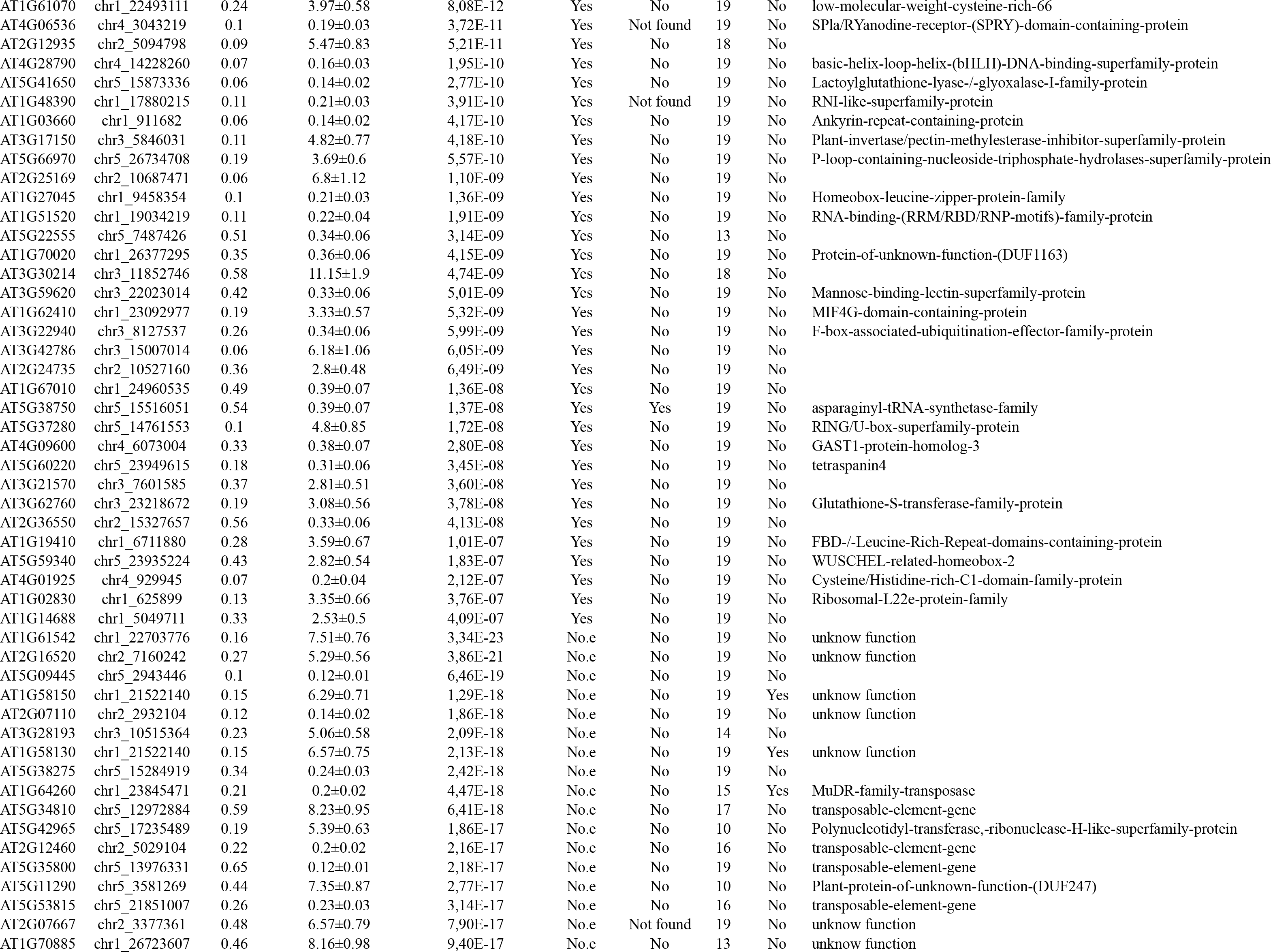
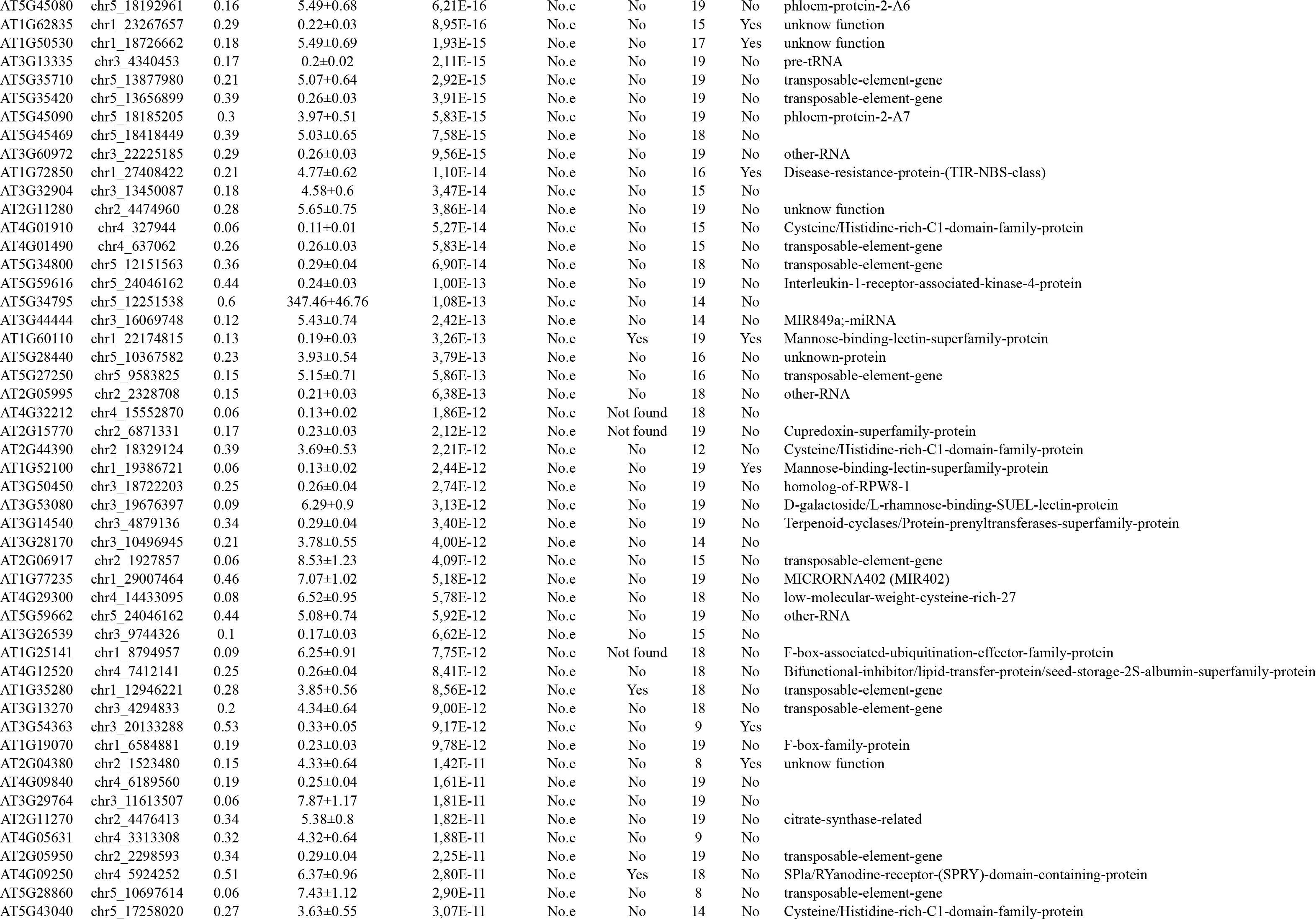
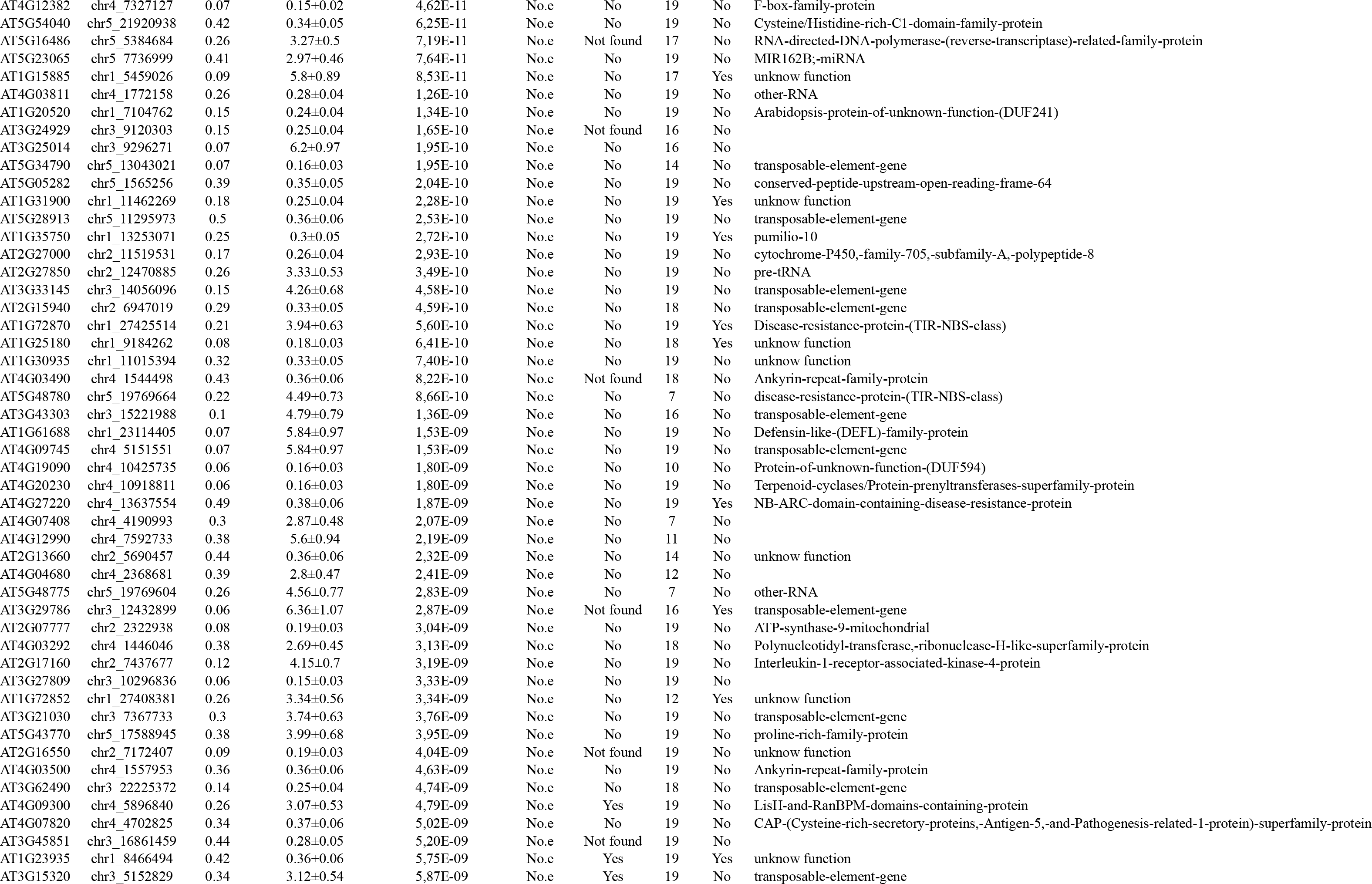
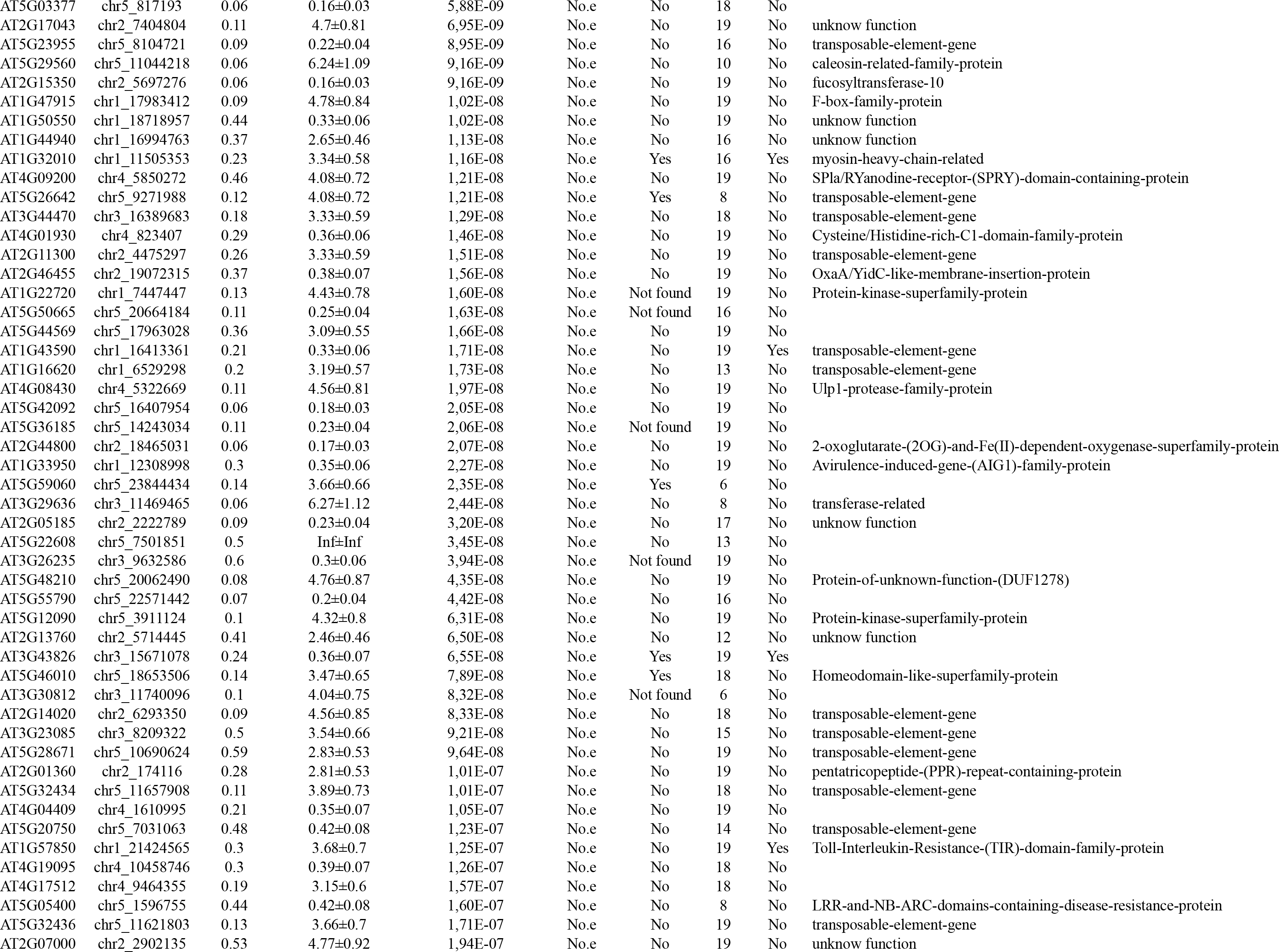
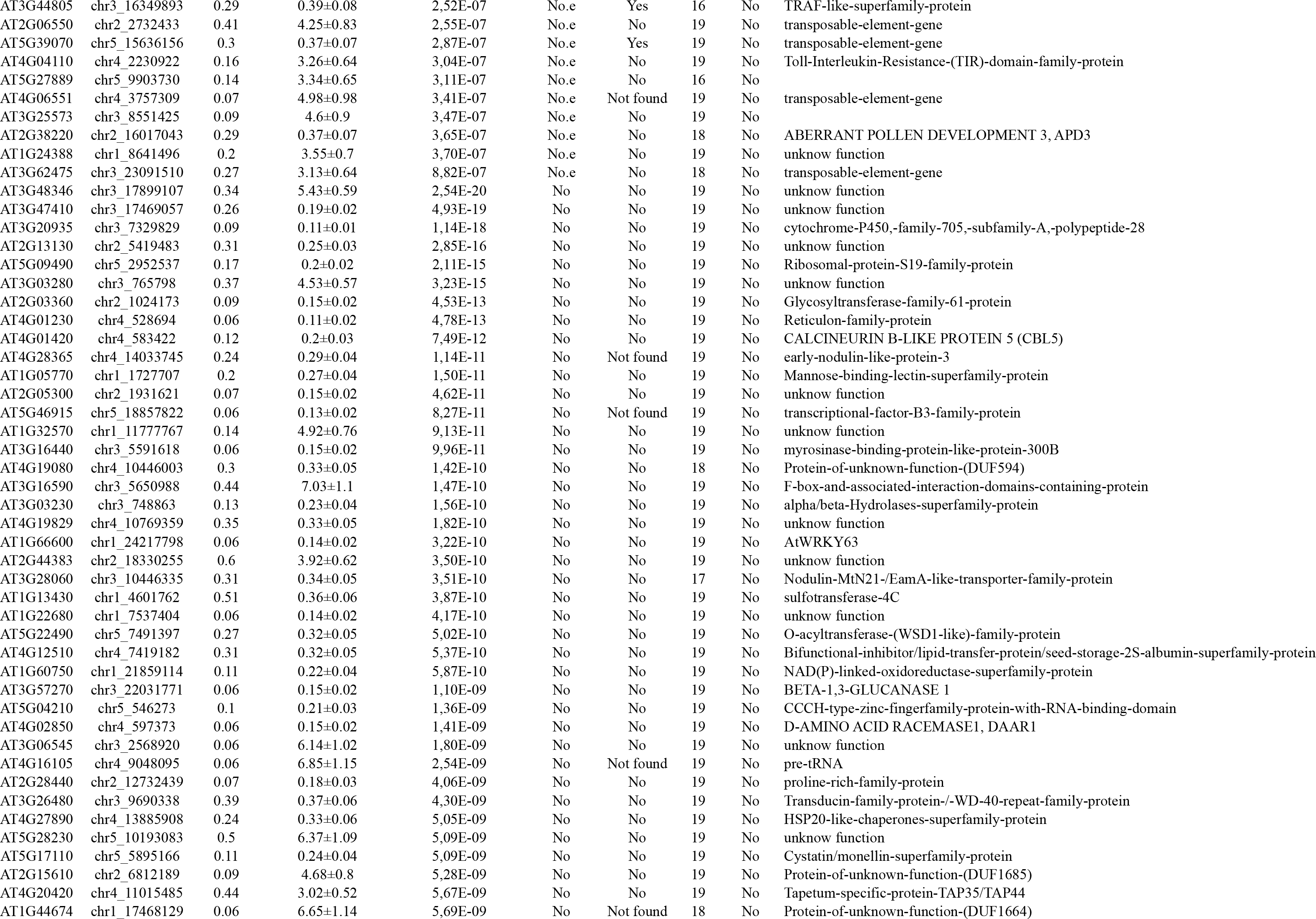
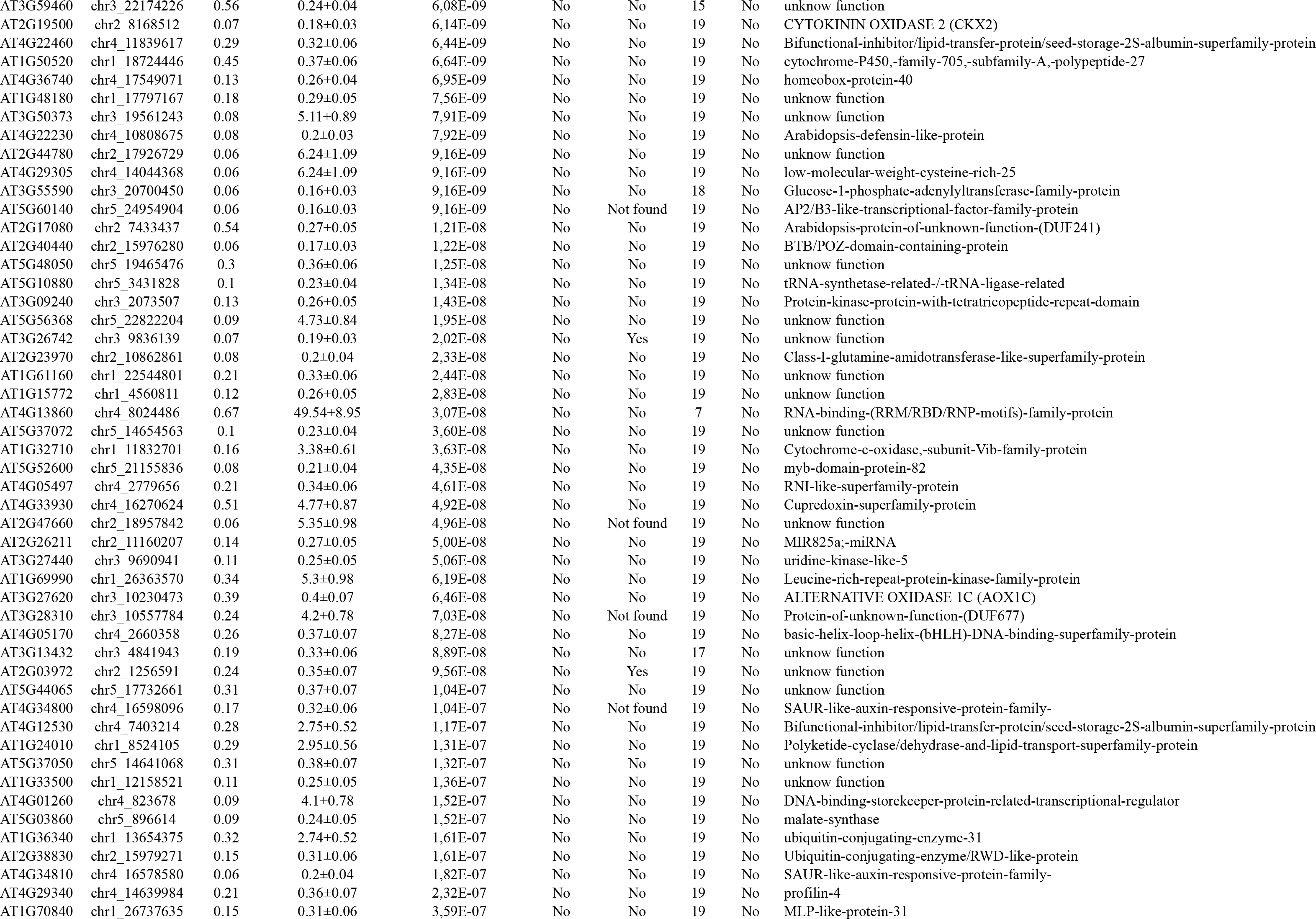
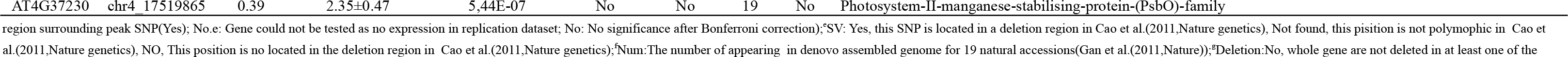
349 cis-eQTL detected in the population of 140 natural *A. thaliana* accessions.

**Table S3.** 81 cis-eQTL detected in the population of 140 natural *A. thaliana* accessions that were replicated in the population of 107 natural Swedish *A. thaliana* accessions.

| *Locus* | *a*SNP | *b*Genotype | *Schmitz-data* |  |  | *Dubin-data* |  |  |
| --- | --- | --- | --- | --- | --- | --- | --- | --- |
|  |  |  | *c*Odds ratio±SE | *d*MAF | *e*P-value | *c*Odds ratio±SE | *d*MAF | *e*P-value |
| AT1G02830 | chr1_624687 | C/C | 2.97±0.59 | 0.29 | 5.51E-07 | 1.89±0.61 | 0.2 | 1.82E-03 |
| AT1G03660 | chr1_926200 | A/A | 0.33±0.09 | 0.32 | 1.29E-04 | 0.45±0.13 | 0.5 | 4.35E-04 |
| AT1G05780 | chr1_1730502 | C/C | 0.32±0.07 | 0.06 | 8.55E-07 | 0.18±0.02 | 0.29 | 5.01E-14 |
| AT1G11370 | chr1_3828411 | T/T | 3.79±0.61 | 0.06 | 3.77E-10 | 4.94±0.78 | 0.1 | 2.48E-10 |
| AT1G14688 | chr1_5130918 | C/C | 0.57±0.25 | 0.09 | 2.09E-02 | 2.23±0.74 | 0.28 | 2.67E-03 |
| AT1G19410 | chr1_6703778 | A/A | 3.29±0.62 | 0.06 | 1.33E-07 | 1.99±0.65 | 0.2 | 2.23E-03 |
| AT1G22090 | chr1_7796024 | T/T | 3.02±0.68 | 0.39 | 8.90E-06 | 0.19±0.02 | 0.38 | 1.52E-15 |
| AT1G27045 | chr1_9947683 | A/A | 2.19±1.1 | 0.33 | 4.71E-02 | 2.06±0.67 | 0.07 | 2.02E-03 |
| AT1G33930 | chr1_12308998 | T/T | 0.28±0.04 | 0.06 | 4.41E-13 | 0.22±0.03 | 0.23 | 1.01E-13 |
| AT1G34100 | chr1_12399605 | A/A | 19.39±2.32 | 0.07 | 6.09E-17 | 0.27±0.05 | 0.06 | 1.30E-07 |
| AT1G48390 | chr1_17879580 | A/A | 0.32±0.07 | 0.42 | 8.55E-07 | 0.55±0.17 | 0.31 | 1.63E-03 |
| AT1G50050 | chr1_18564085 | C/C | 2.13±0.97 | 0.2 | 2.86E-02 | 3.44±1.29 | 0.25 | 7.68E-03 |
| AT1G51520 | chr1_19053511 | T/T | 0.49±0.15 | 0.12 | 8.64E-04 | 2.01±0.63 | 0.21 | 1.34E-03 |
| AT1G51990 | chr1_19304765 | G/G | 2.35±0.5 | 0.16 | 2.09E-06 | 0.36±0.08 | 0.21 | 4.87E-06 |
| AT1G61070 | chr1_22320587 | A/A | 0.56±0.27 | 0.09 | 4.08E-02 | 6.46±1.53 | 0.5 | 2.47E-05 |
| AT1G62410 | chr1_23196263 | T/T | 2.01±0.58 | 0.09 | 5.16E-04 | 2.75±1.04 | 0.2 | 8.20E-03 |
| AT1G64235 | chr1_23877480 | T/T | 3±0.6 | 0.16 | 4.59E-07 | 9.26±1 | 0.27 | 1.58E-20 |
| AT1G67010 | chr1_24961784 | A/A | 3.1±0.88 | 0.26 | 4.00E-04 | 0.38±0.13 | 0.07 | 4.54E-03 |
| AT1G67148 | chr1_25121971 | A/A | 0.26±0.04 | 0.53 | 3.53E-12 | 2.08±0.65 | 0.24 | 1.52E-03 |
| AT1G70020 | chr1_26377468 | C/C | 0.33±0.06 | 0.06 | 1.05E-08 | 2.48±0.52 | 0.08 | 1.85E-06 |
| AT1G70870 | chr1_26719733 | G/G | 0.3±0.05 | 0.12 | 1.55E-09 | 0.24±0.05 | 0.36 | 5.33E-07 |
| AT1G73120 | chr1_27497719 | G/G | 3.33±0.73 | 0.06 | 4.61E-06 | 4.2±0.75 | 0.11 | 2.33E-08 |
| AT1G76830 | chr1_28830886 | A/A | 1.71±0.7 | 0.26 | 1.38E-02 | 6.46±1.53 | 0.38 | 2.47E-05 |
| AT1G78030 | chr1_29342131 | A/A | 0.43±0.09 | 0.32 | 9.54E-07 | 4.65±0.64 | 0.2 | 3.34E-13 |
| AT2G04830 | chr2_1693588 | A/A | 5.52±0.76 | 0.19 | 4.84E-13 | 2.5±0.73 | 0.31 | 5.95E-04 |
| AT2G05117 | chr2_1831031 | T/T | 0.27±0.04 | 0.19 | 4.86E-13 | 0.11±0.02 | 0.1 | 1.12E-09 |
| AT2G10260 | chr2_3235727 | G/G | 2.63±0.54 | 0.19 | 1.08E-06 | 0.35±0.13 | 0.36 | 7.31E-03 |
| AT2G12935 | chr2_5190356 | A/A | 2.34±0.57 | 0.3 | 3.86E-05 | 2.4±1.07 | 0.42 | 2.58E-02 |
| AT2G13128 | chr2_5108880 | T/T | 0.3±0.05 | 0.64 | 1.52E-08 | 0.6±0.29 | 0.27 | 3.64E-02 |
| AT2G15930 | chr2_6939722 | A/A | 3.32±0.54 | 0.06 | 9.21E-10 | 4.18±0.82 | 0.24 | 3.36E-07 |
| AT2G21045 | chr2_8997433 | A/A | 4.56±0.81 | 0.06 | 1.97E-08 | 4.3±0.76 | 0.09 | 1.50E-08 |
| AT2G24165 | chr2_10272982 | A/A | 0.27±0.04 | 0.05 | 2.17E-10 | 0.16±0.02 | 0.07 | 3.45E-12 |
| AT2G24735 | chr2_10523189 | C/C | 3.73±0.65 | 0.06 | 8.47E-09 | 3.18±0.61 | 0.36 | 2.12E-07 |
| AT2G25169 | chr2_10851248 | T/T | 0.92±6.21 | 0.29 | 8.82E-01 | 5.64±2.05 | 0.28 | 5.88E-03 |
| AT2G29000 | chr2_12631021 | A/A | 2.11±0.6 | 0.42 | 4.49E-04 | 2.51±0.77 | 0.25 | 1.15E-03 |
| AT2G36550 | chr2_15327657 | A/A | 0.33±0.06 | 0.53 | 4.13E-08 | 0.2±0.05 | 0.27 | 9.16E-05 |
| AT2G45940 | chr2_18902487 | C/C | 0.46±0.11 | 0.12 | 3.75E-05 | 0.29±0.08 | 0.47 | 5.08E-04 |
| AT3G17150 | chr3_5842339 | G/G | 2.38±0.63 | 0.46 | 1.60E-04 | 2.25±0.89 | 0.24 | 1.12E-02 |
| AT3G21570 | chr3_7727067 | A/A | 3.44±0.97 | 0.54 | 3.78E-04 | 1.92±0.58 | 0.5 | 8.59E-04 |
| AT3G22940 | chr3_8127537 | G/G | 0.34±0.06 | 0.21 | 5.99E-09 | 0.57±0.2 | 0.21 | 4.31E-03 |
| AT3G25719 | chr3_9381875 | G/G | 3.01±0.46 | 0.22 | 4.17E-11 | 0.11±0.01 | 0.35 | 4.19E-14 |
| AT3G25720 | chr3_9381875 | G/G | 3.27±0.46 | 0.22 | 1.06E-12 | 0.12±0.02 | 0.35 | 1.91E-15 |
| AT3G30214 | chr3_11891239 | T/T | 0.43±0.09 | 0.43 | 6.08E-07 | 3.83±1.08 | 0.15 | 3.78E-04 |
| AT3G42786 | chr3_14917003 | G/G | 1.52±0.67 | 0.38 | 2.41E-02 | 4.29±1.41 | 0.26 | 2.38E-03 |
| AT3G46482 | chr3_17105125 | C/C | 4.67±0.93 | 0.06 | 5.45E-07 | 3.67±0.61 | 0.1 | 1.92E-09 |
| AT3G53840 | chr3_19949277 | C/C | 0.52±0.14 | 0.39 | 1.79E-04 | 0.38±0.08 | 0.13 | 1.51E-06 |
| AT3G58480 | chr3_21628568 | T/T | 0.39±0.08 | 0.31 | 2.28E-06 | 0.23±0.03 | 0.21 | 2.70E-11 |
| AT3G59180 | chr3_21760130 | A/A | 2.22±0.57 | 0.07 | 1.05E-04 | 1.64±0.7 | 0.07 | 1.90E-02 |
| AT3G59190 | chr3_22175616 | T/T | 0.23±0.04 | 0.22 | 1.45E-08 | 0.54±0.27 | 0.28 | 4.75E-02 |
| AT3G59620 | chr3_22023014 | C/C | 0.33±0.06 | 0.06 | 5.01E-09 | 0.2±0.08 | 0.06 | 1.76E-02 |
| AT3G62760 | chr3_23189064 | T/T | 0.43±0.14 | 0.11 | 1.80E-03 | 1.96±0.9 | 0.1 | 2.86E-02 |
| AT4G01190 | chr4_506690 | C/C | 42.47±6.2 | 0.54 | 7.44E-12 | 3.96±0.55 | 0.36 | 4.88E-13 |
| AT4G01335 | chr4_463836 | T/T | 2.42±0.97 | 0.56 | 1.29E-02 | 8.81±1.75 | 0.45 | 5.11E-07 |
| AT4G01925 | chr4_822143 | A/A | 4.16±0.89 | 0.08 | 3.32E-06 | 2.51±0.63 | 0.43 | 6.66E-05 |
| AT4G06536 | chr4_3334822 | C/C | 0.34±0.07 | 0.09 | 3.68E-06 | 0.57±0.26 | 0.34 | 2.85E-02 |
| AT4G09012 | chr4_5784087 | G/G | 0.47±0.11 | 0.29 | 8.72E-06 | 3.63±0.86 | 0.44 | 2.47E-05 |
| AT4G09600 | chr4_6074156 | T/T | 6.75±1.31 | 0.2 | 2.67E-07 | 1.73±0.64 | 0.49 | 6.64E-03 |
| AT4G15053 | chr4_8598649 | T/T | 6.51±1.1 | 0.57 | 3.33E-09 | 5.89±1.02 | 0.31 | 8.31E-09 |
| AT4G22390 | chr4_11846652 | G/G | 0.24±0.03 | 0.06 | 3.33E-16 | 0.32±0.07 | 0.29 | 1.38E-06 |
| AT4G28790 | chr4_14357466 | A/A | 0.32±0.09 | 0.4 | 4.82E-04 | 3.56±1.79 | 0.25 | 4.62E-02 |
| AT5G03820 | chr5_1016604 | G/G | 0.24±0.03 | 0.2 | 1.06E-13 | 0.31±0.1 | 0.08 | 2.64E-03 |
| AT5G08460 | chr5_2734328 | T/T | 20±2.28 | 0.39 | 2.04E-18 | 6.24±0.93 | 0.29 | 1.62E-11 |
| AT5G14290 | chr5_5004746 | T/T | 1.65±0.58 | 0.34 | 4.34E-03 | 1.7±0.84 | 0.2 | 4.21E-02 |
| AT5G22555 | chr5_7487426 | A/A | 0.34±0.06 | 0.11 | 3.14E-09 | 0.38±0.09 | 0.28 | 1.33E-05 |
| AT5G22680 | chr5_7625326 | G/G | 0.52±0.15 | 0.24 | 5.32E-04 | 2.56±1.28 | 0.28 | 4.50E-02 |
| AT5G24040 | chr5_8150163 | C/C | 3.67±0.79 | 0.1 | 3.75E-06 | 9.03±1.34 | 0.08 | 1.74E-11 |
| AT5G24206 | chr5_8210609 | A/A | 4.59±0.9 | 0.59 | 3.04E-07 | 0.14±0.02 | 0.39 | 2.96E-09 |
| AT5G34830 | chr5_12151563 | G/G | 0.2±0.03 | 0.06 | 7.97E-13 | 2.26±0.79 | 0.16 | 4.03E-03 |
| AT5G36240 | chr5_14277903 | G/G | 3.33±0.89 | 0.56 | 1.83E-04 | 4.52±0.75 | 0.17 | 2.17E-09 |
| AT5G37280 | chr5_14761553 | T/T | 4.8±0.85 | 0.06 | 1.72E-08 | 3.66±0.96 | 0.38 | 1.32E-04 |
| AT5G38750 | chr5_15513299 | G/G | 0.39±0.08 | 0.11 | 2.57E-07 | 0.38±0.11 | 0.35 | 6.96E-04 |
| AT5G40050 | chr5_16033285 | C/C | 2.15±0.57 | 0.06 | 1.73E-04 | 2.12±0.69 | 0.07 | 1.99E-03 |
| AT5G41650 | chr5_16309822 | T/T | 0.3±0.08 | 0.06 | 1.31E-04 | 0.35±0.16 | 0.49 | 2.82E-02 |
| AT5G47130 | chr5_19140035 | T/T | 0.29±0.04 | 0.29 | 5.54E-14 | 0.51±0.2 | 0.29 | 9.35E-03 |
| AT5G48350 | chr5_19591922 | G/G | 3.79±0.72 | 0.08 | 1.23E-07 | 0.26±0.04 | 0.25 | 1.38E-12 |
| AT5G49420 | chr5_20036039 | G/G | 21.25±4.8 | 0.24 | 9.51E-06 | 0.4±0.13 | 0.35 | 2.81E-03 |
| AT5G54206 | chr5_21999241 | A/A | 0.23±0.03 | 0.32 | 2.06E-13 | 0.36±0.08 | 0.38 | 1.09E-05 |
| AT5G59340 | chr5_24032779 | A/A | 0.53±0.16 | 0.13 | 1.19E-03 | 2.03±0.79 | 0.21 | 9.83E-03 |
| AT5G60220 | chr5_24003551 | G/G | 0.53±0.15 | 0.09 | 6.16E-04 | 2±0.62 | 0.23 | 1.27E-03 |
| AT5G66970 | chr5_26741775 | C/C | 2.02±0.48 | 0.16 | 2.56E-05 | 2.65±0.83 | 0.21 | 1.43E-03 |
| AT5G67040 | chr5_26756966 | C/C | 0.53±0.24 | 0.11 | 2.80E-02 | 5.53±0.98 | 0.22 | 1.41E-08 |

<sup>a</sup>SNP: Overlapping SNP between the two datasets; <sup>b</sup>Genotype: Genotype at the coding allele from SCHMITZ data; <sup>c</sup>odds ratio ±SE: log odds ration ± Standard Error from the logistic regression; <sup>d</sup>MAF: Minor Allele Frequency; <sup>e</sup>P-value: nominal P-value from single-SNP association

**Table S4.**
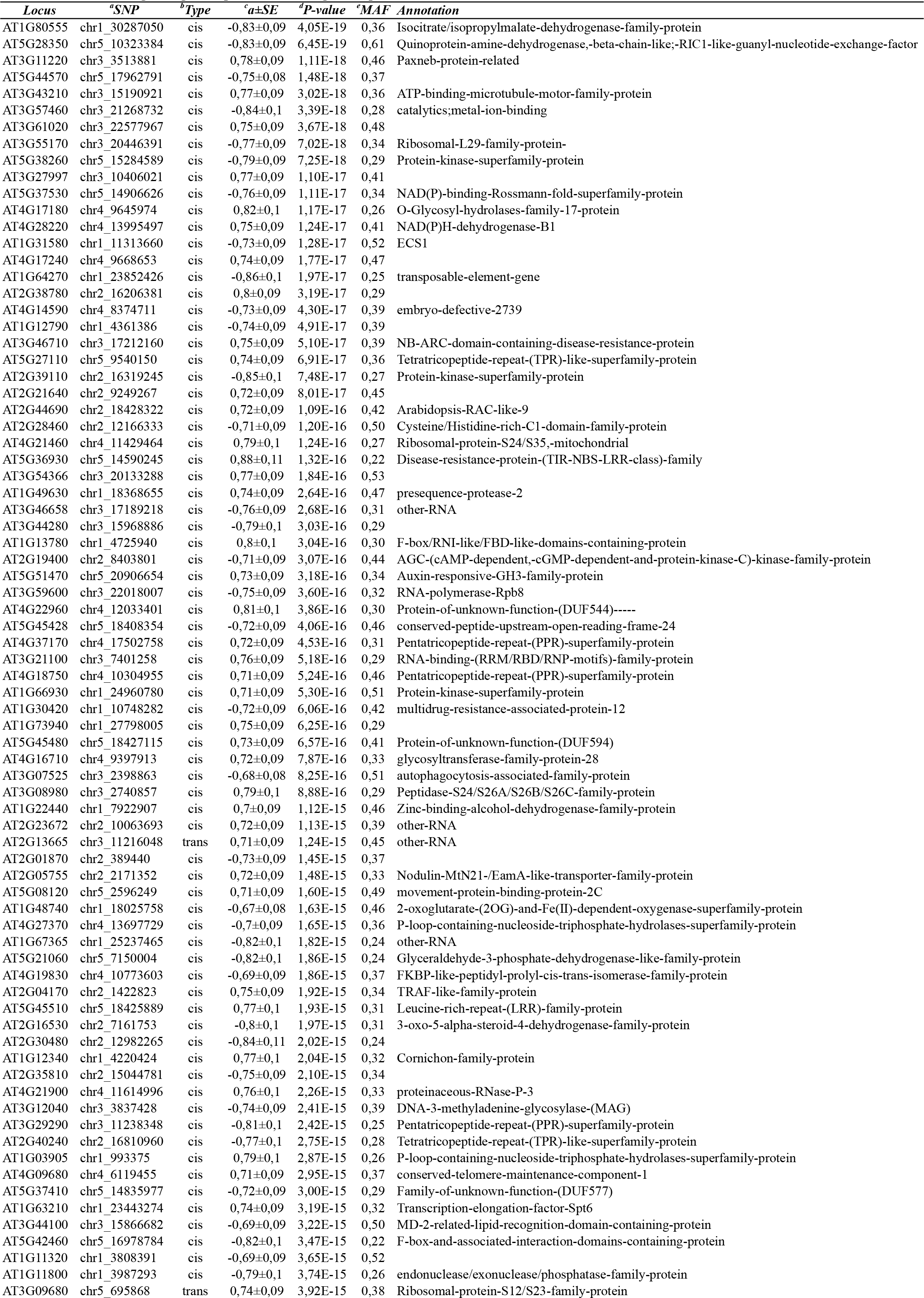
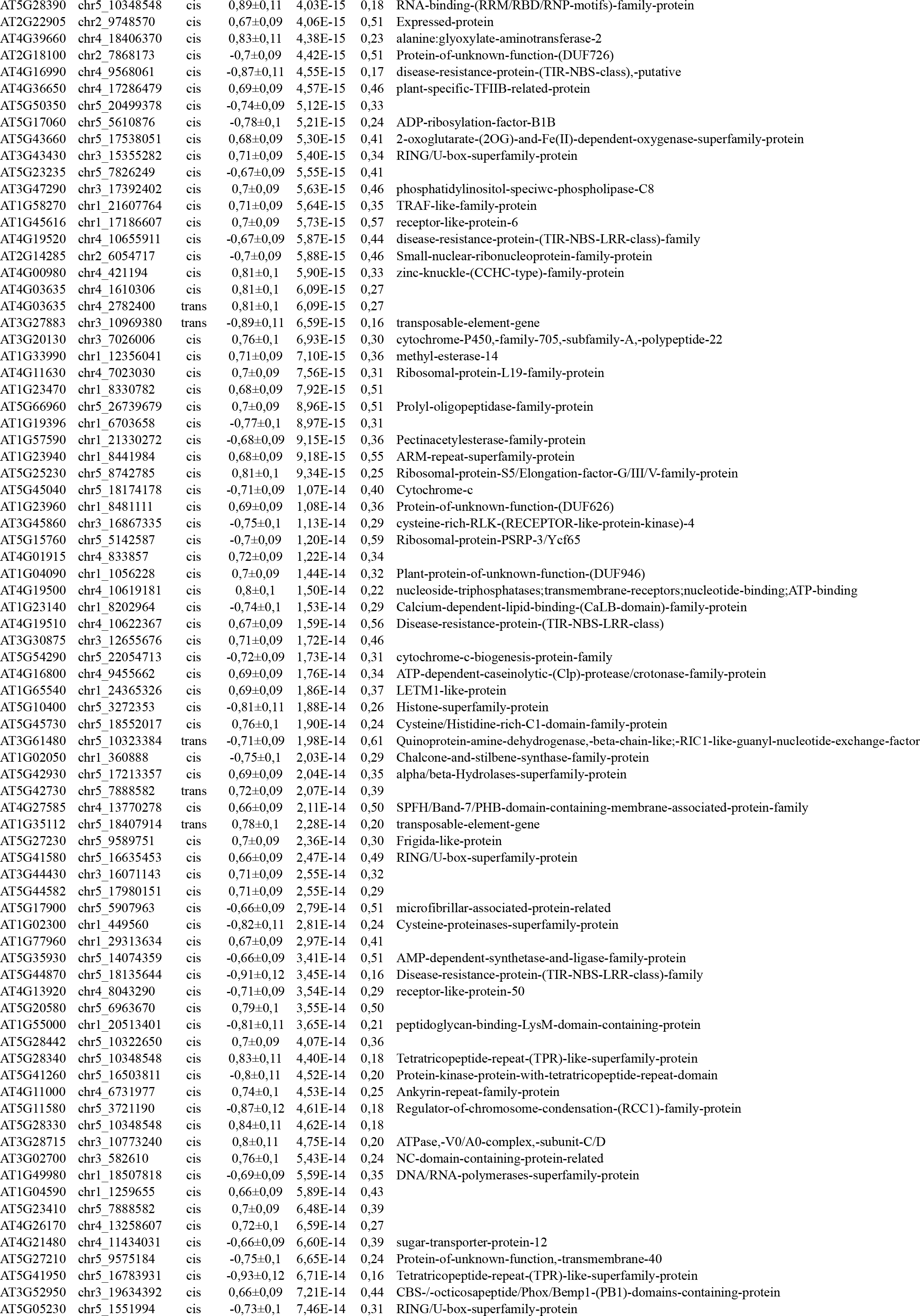
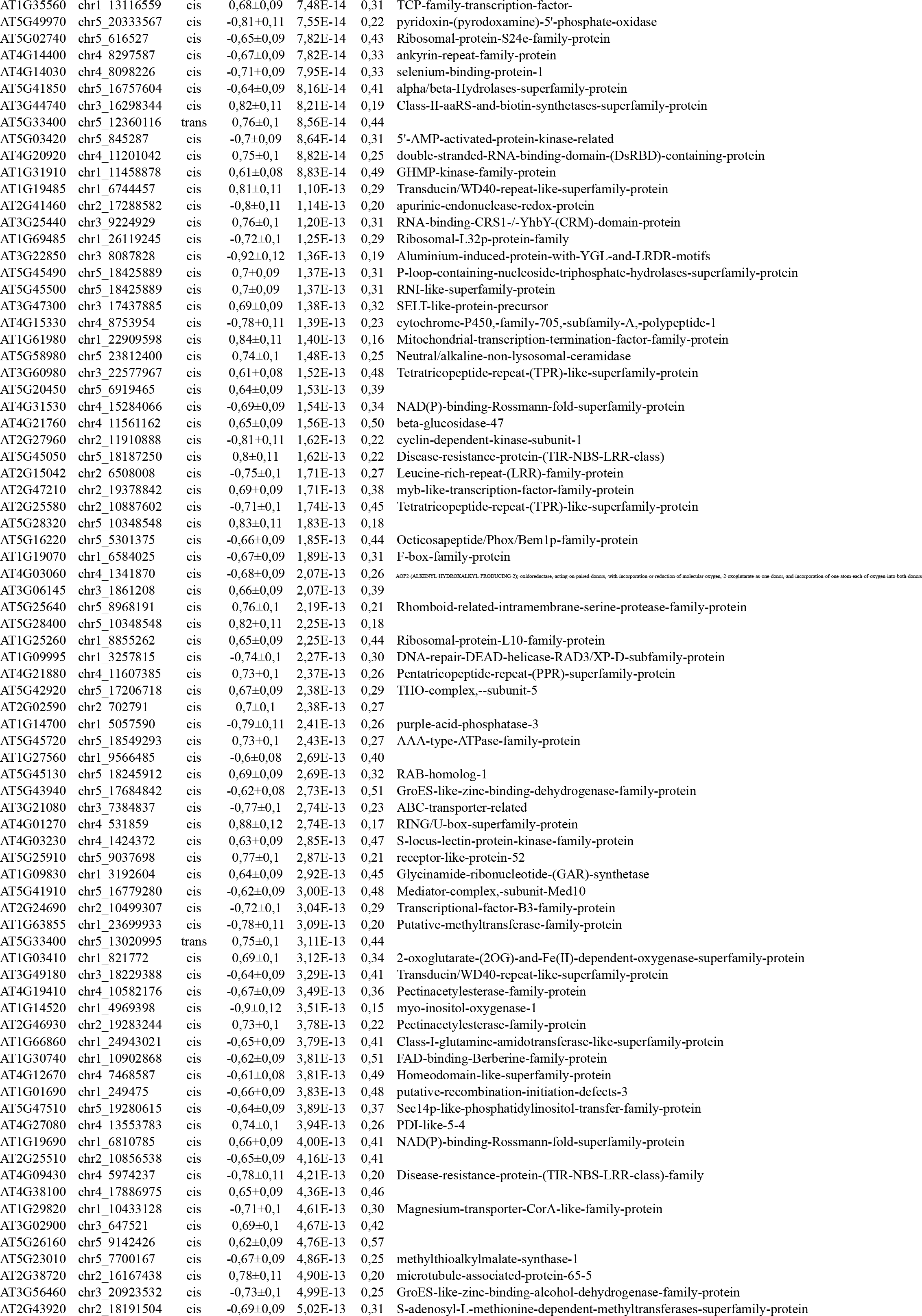
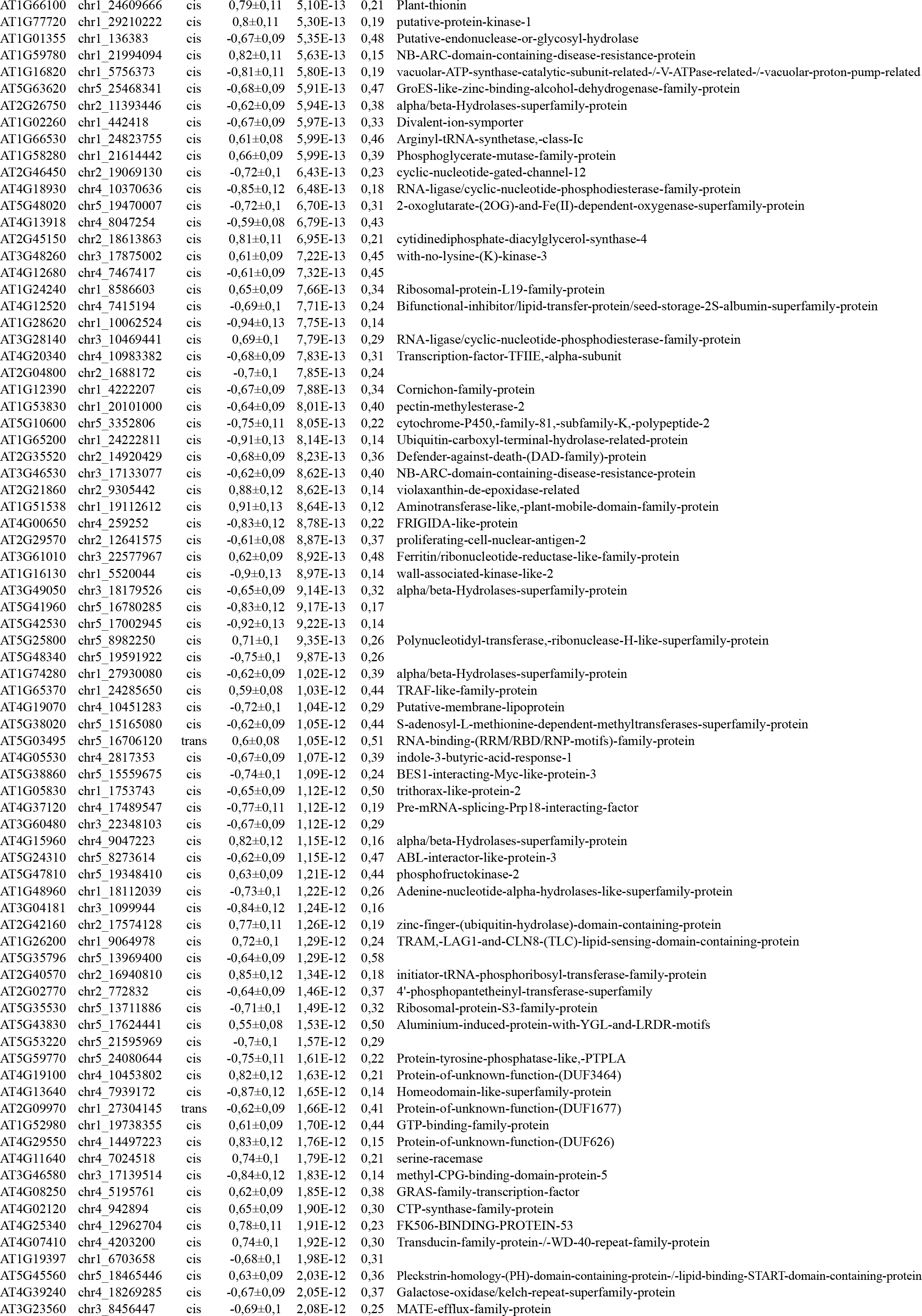
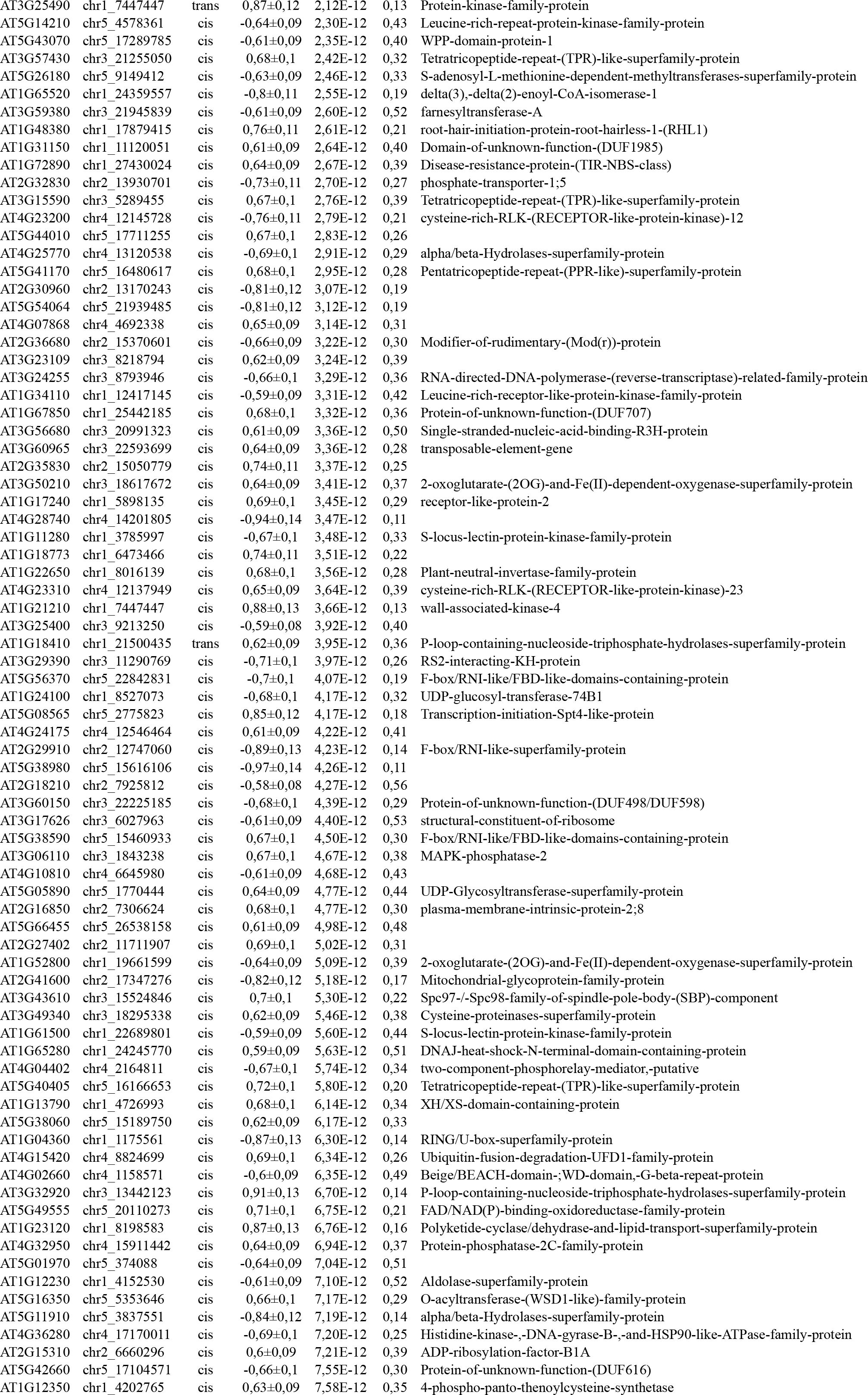
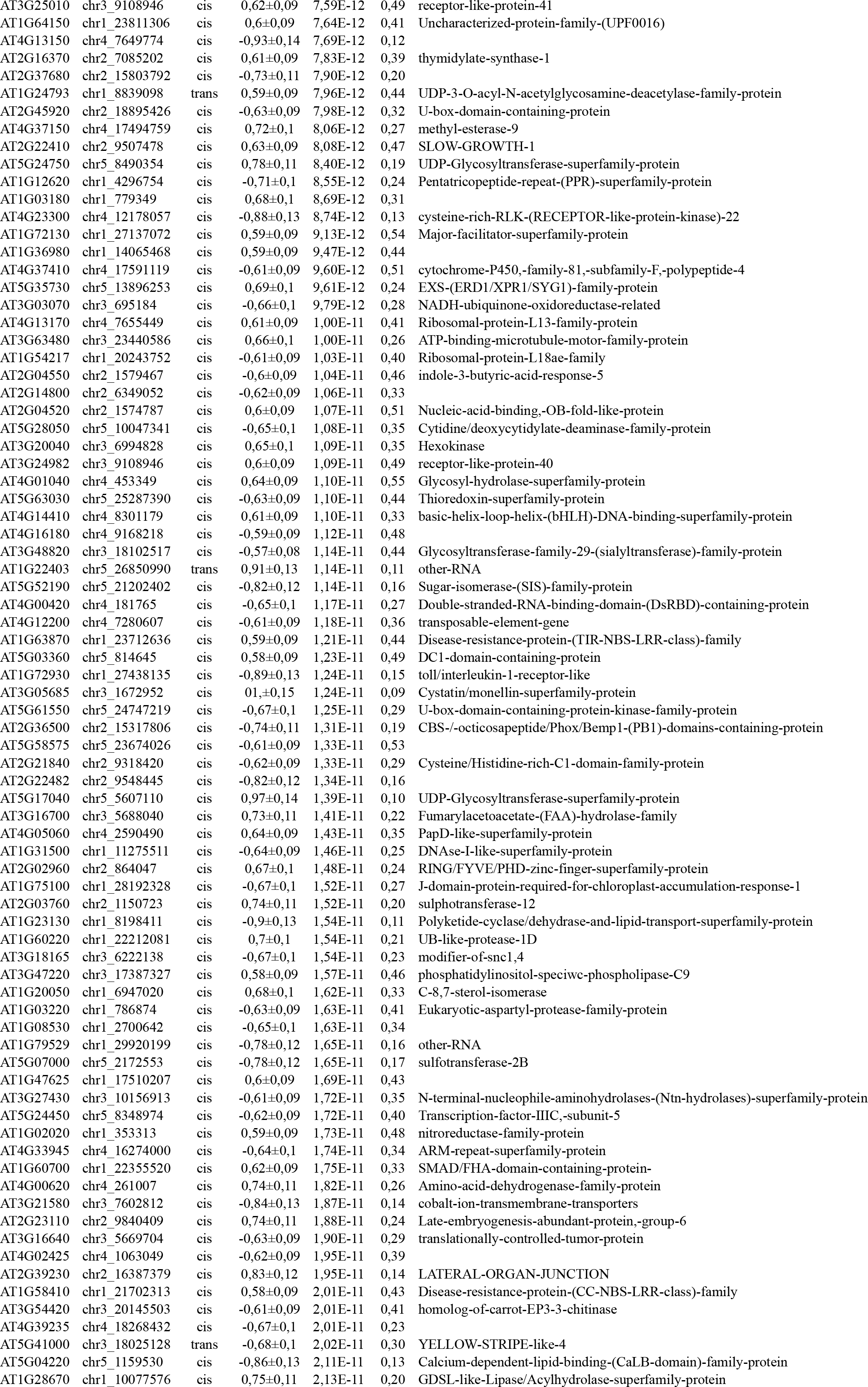
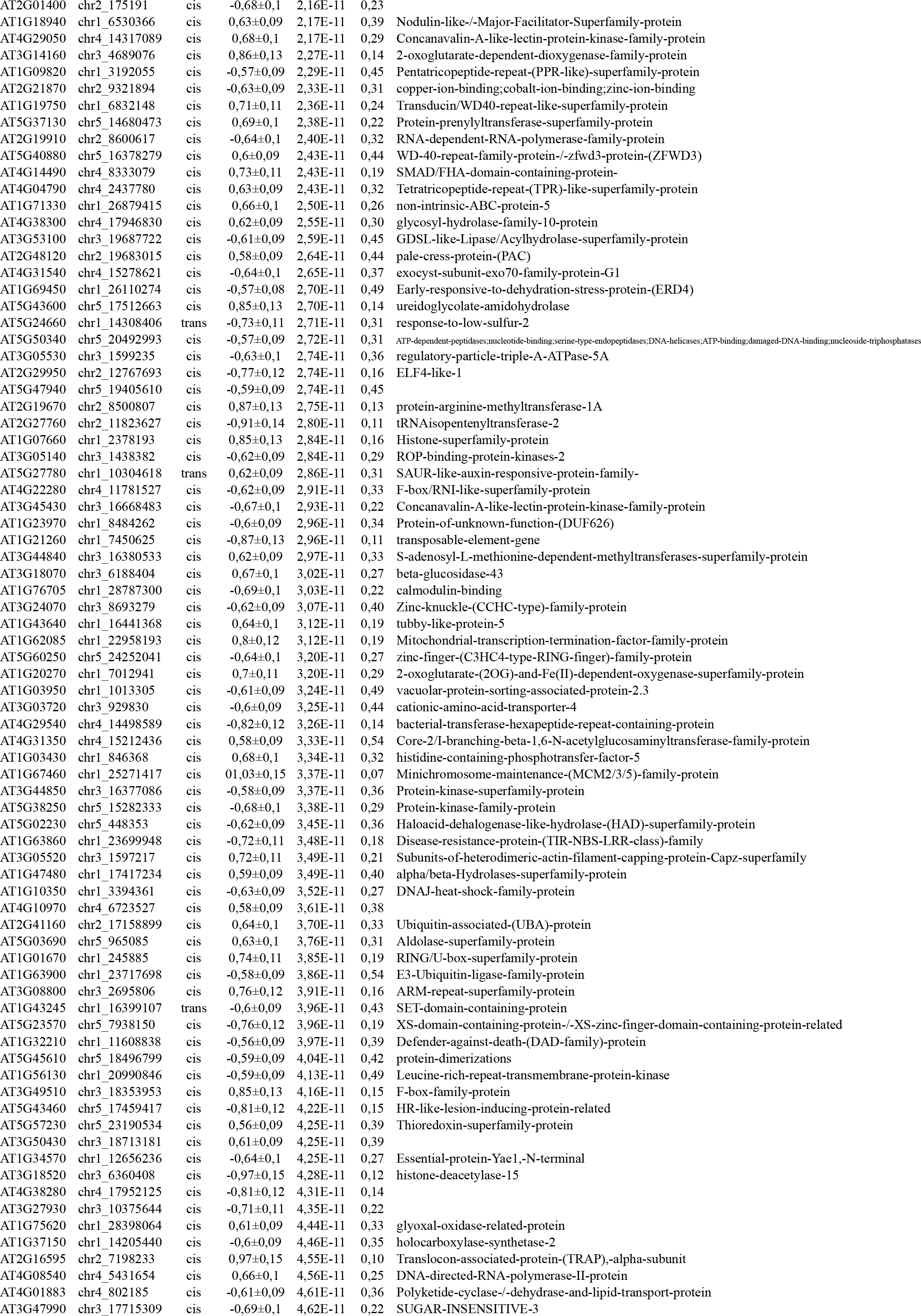
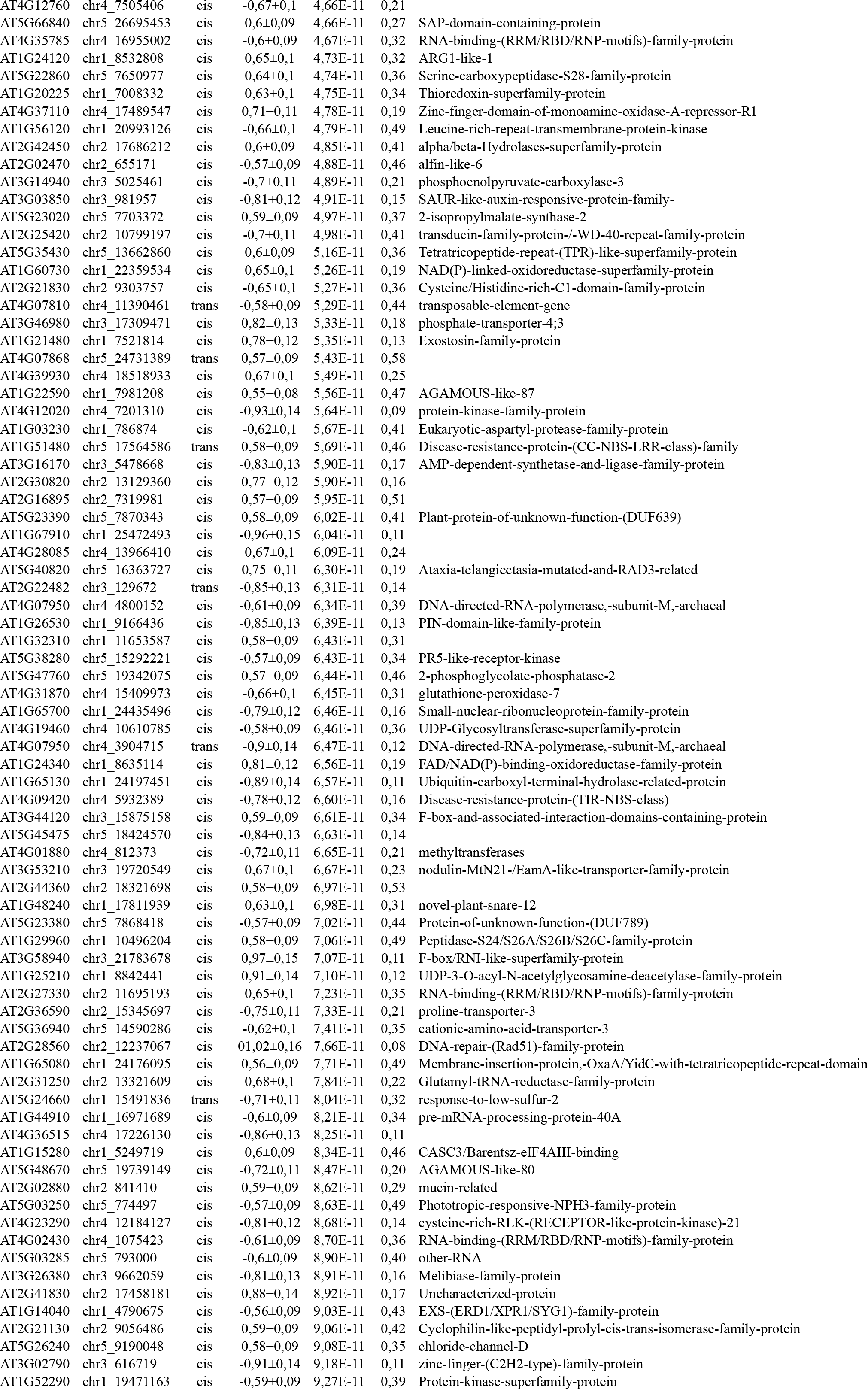
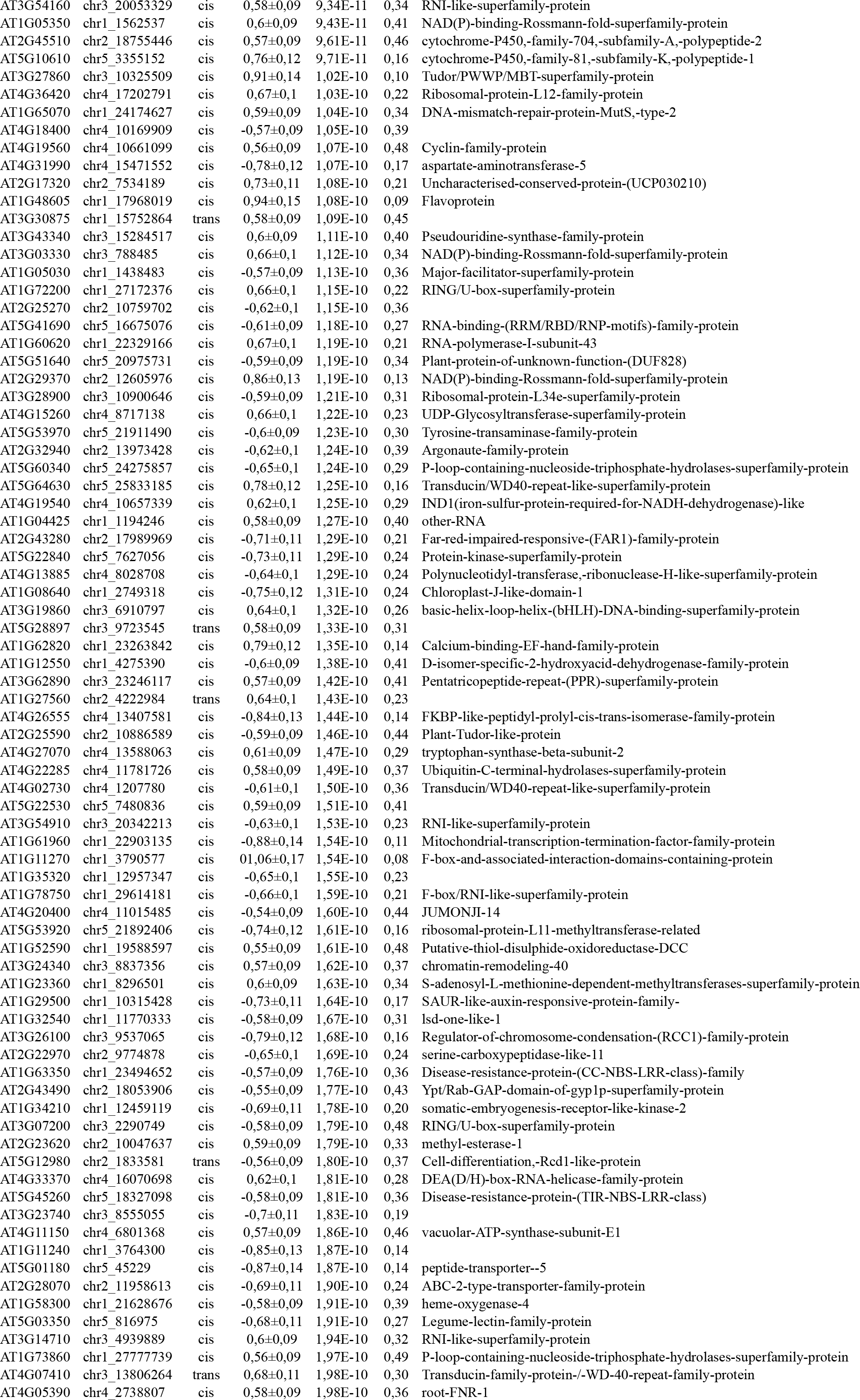
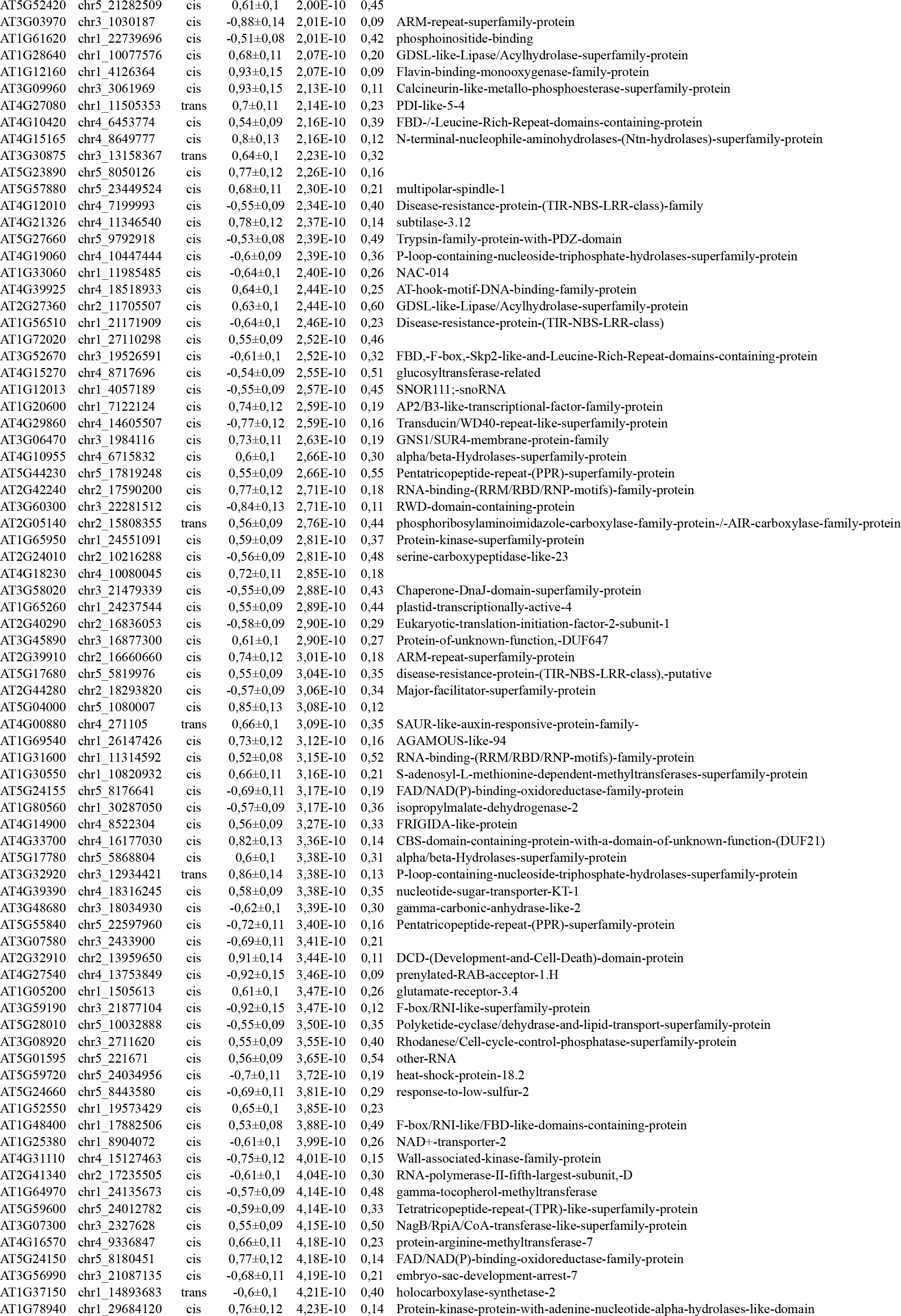
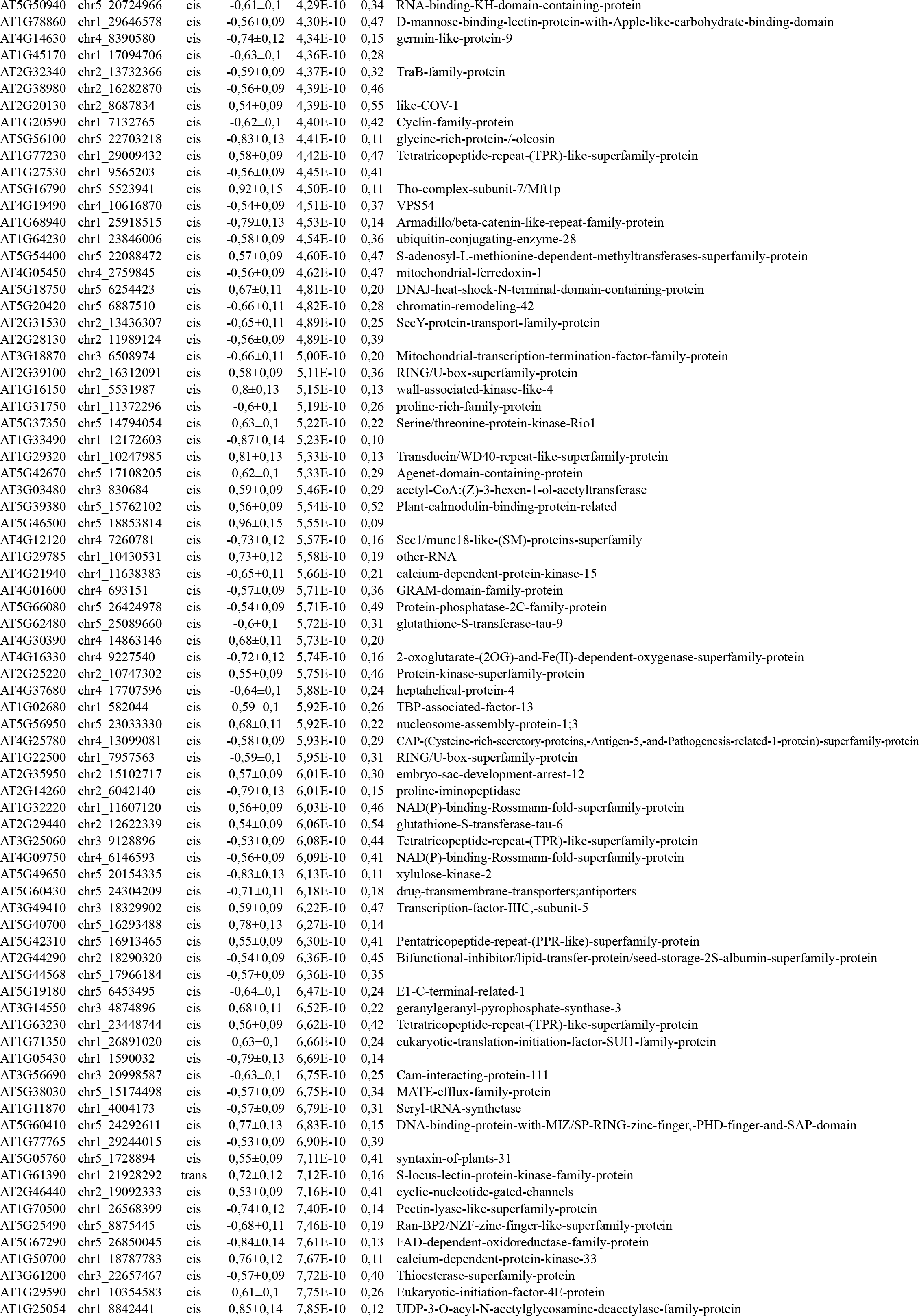
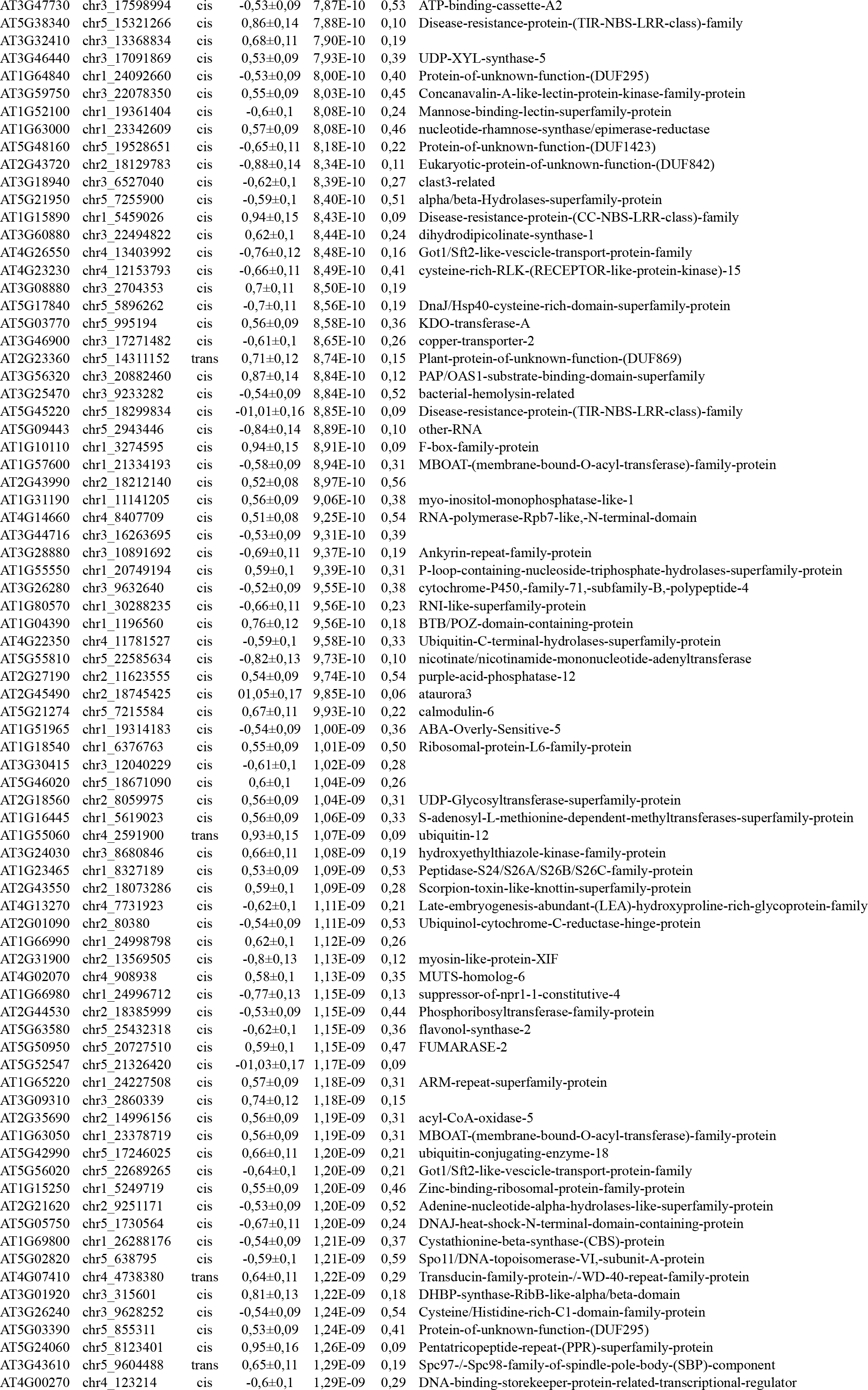
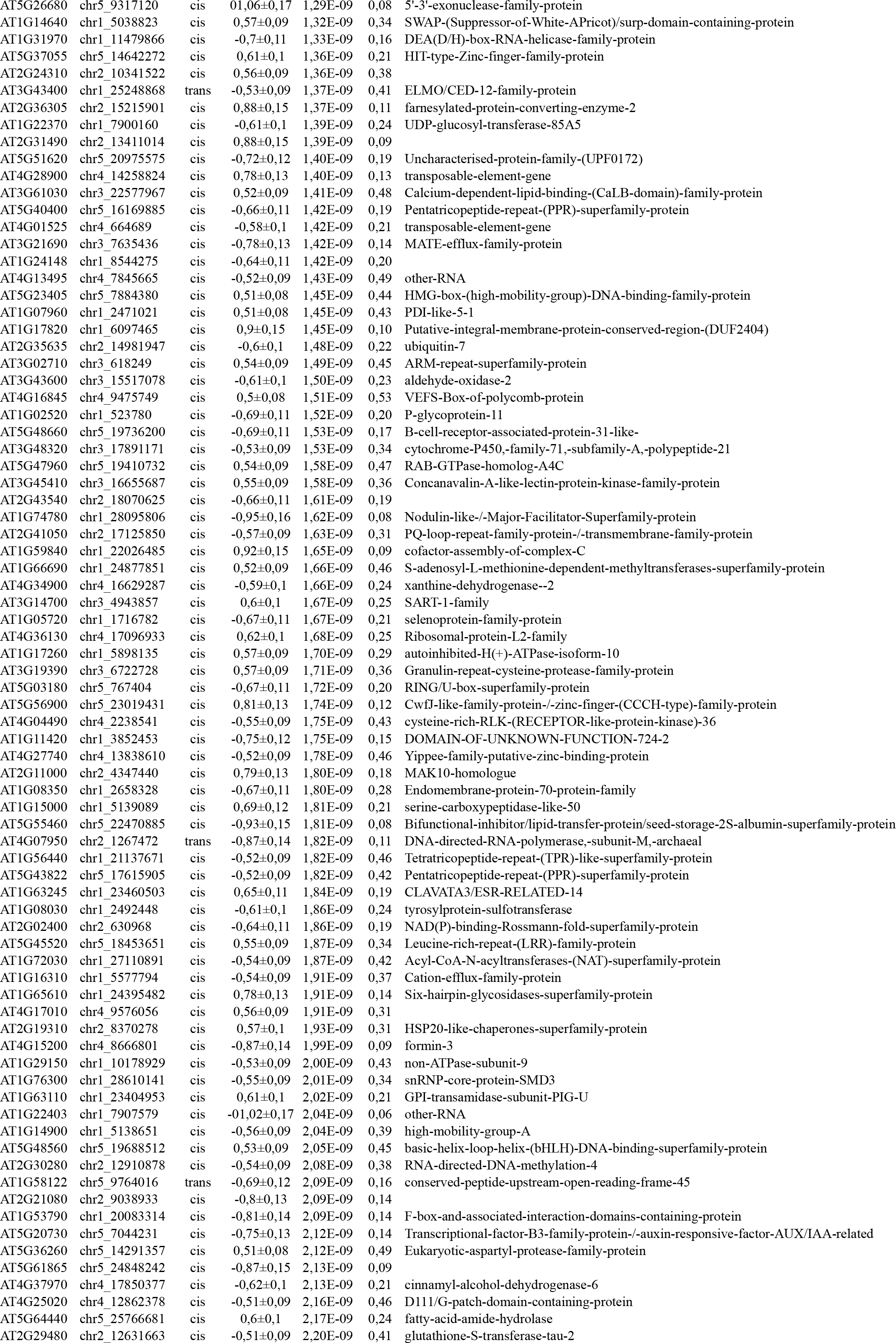
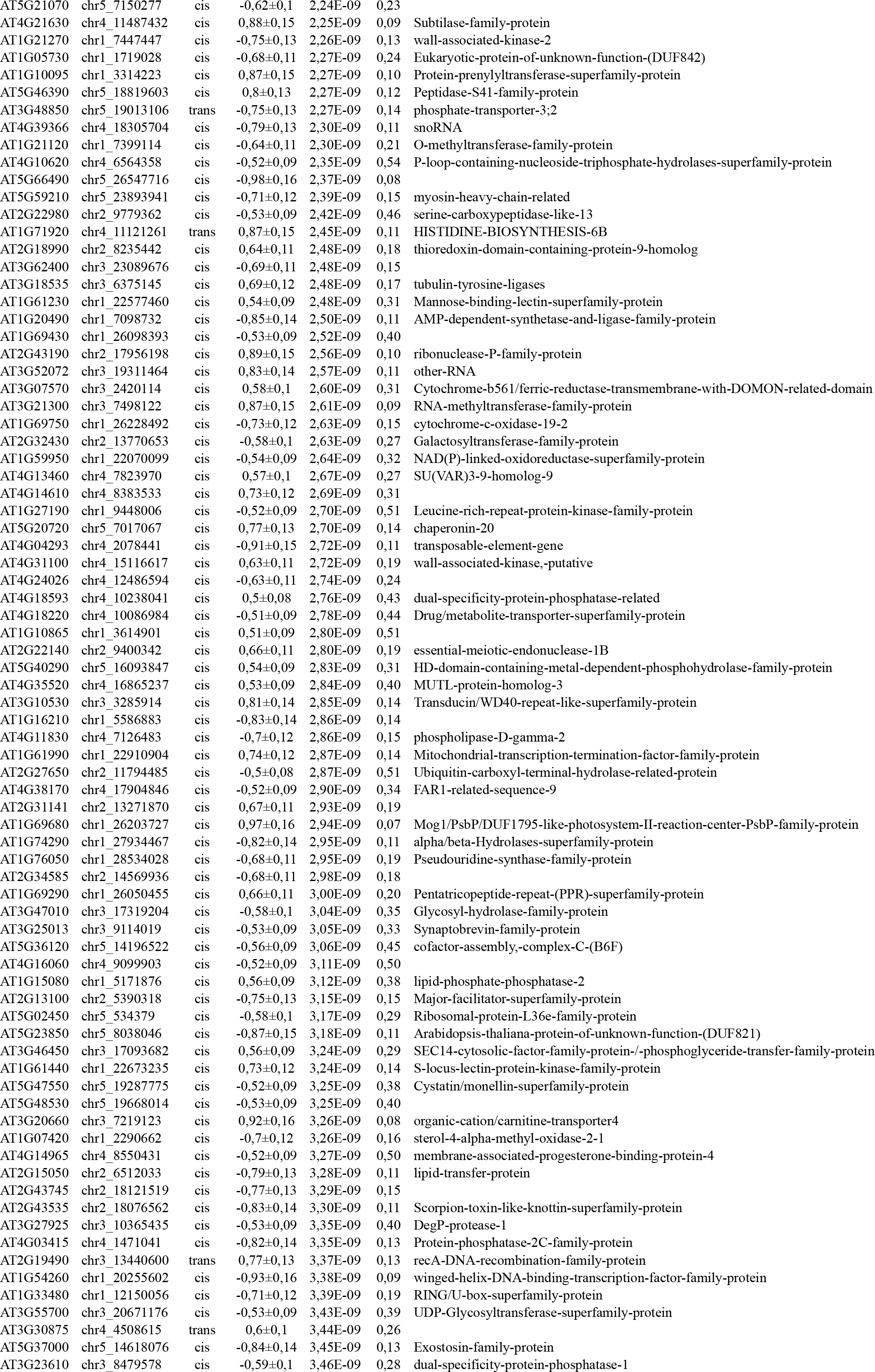
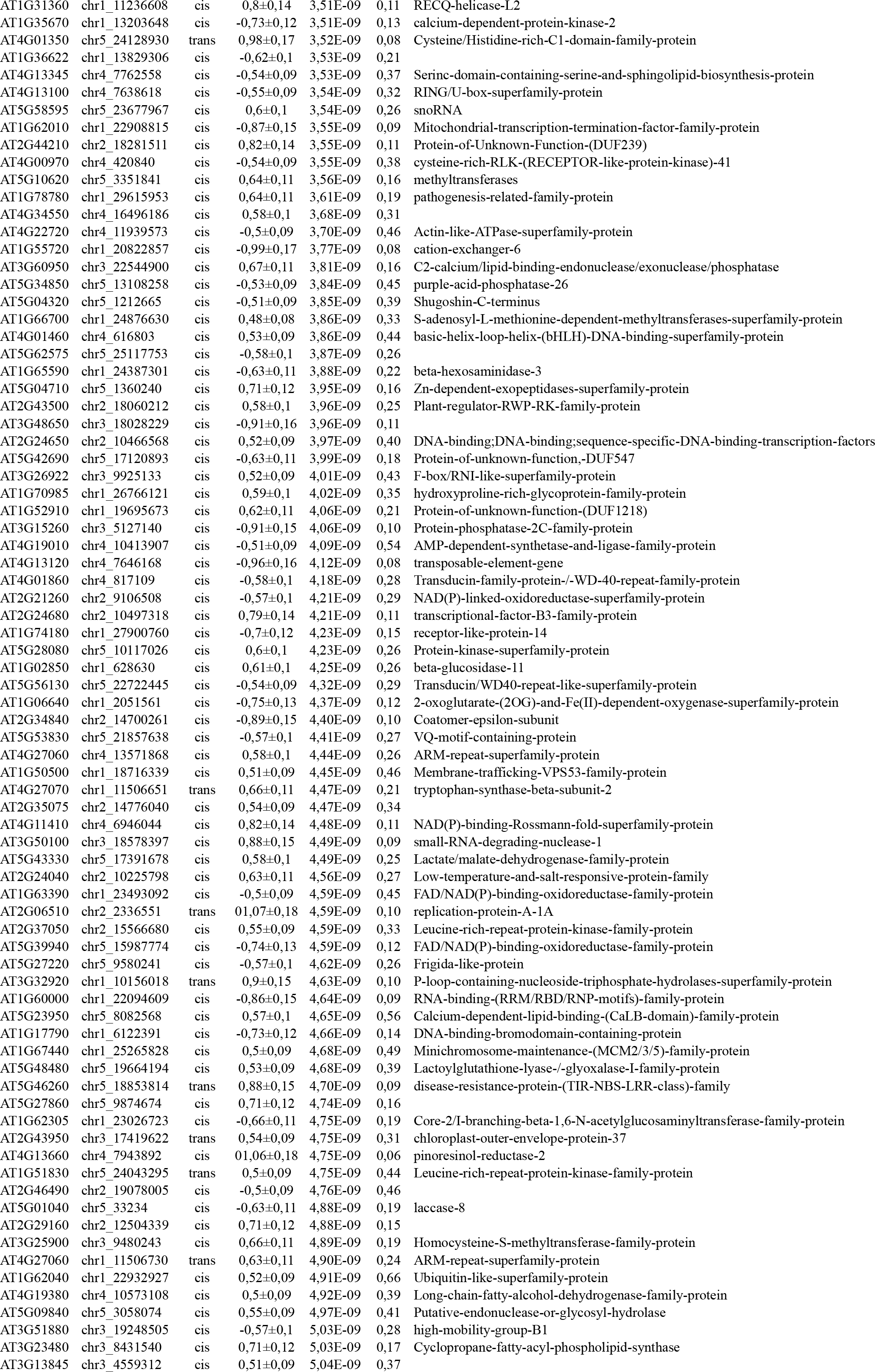
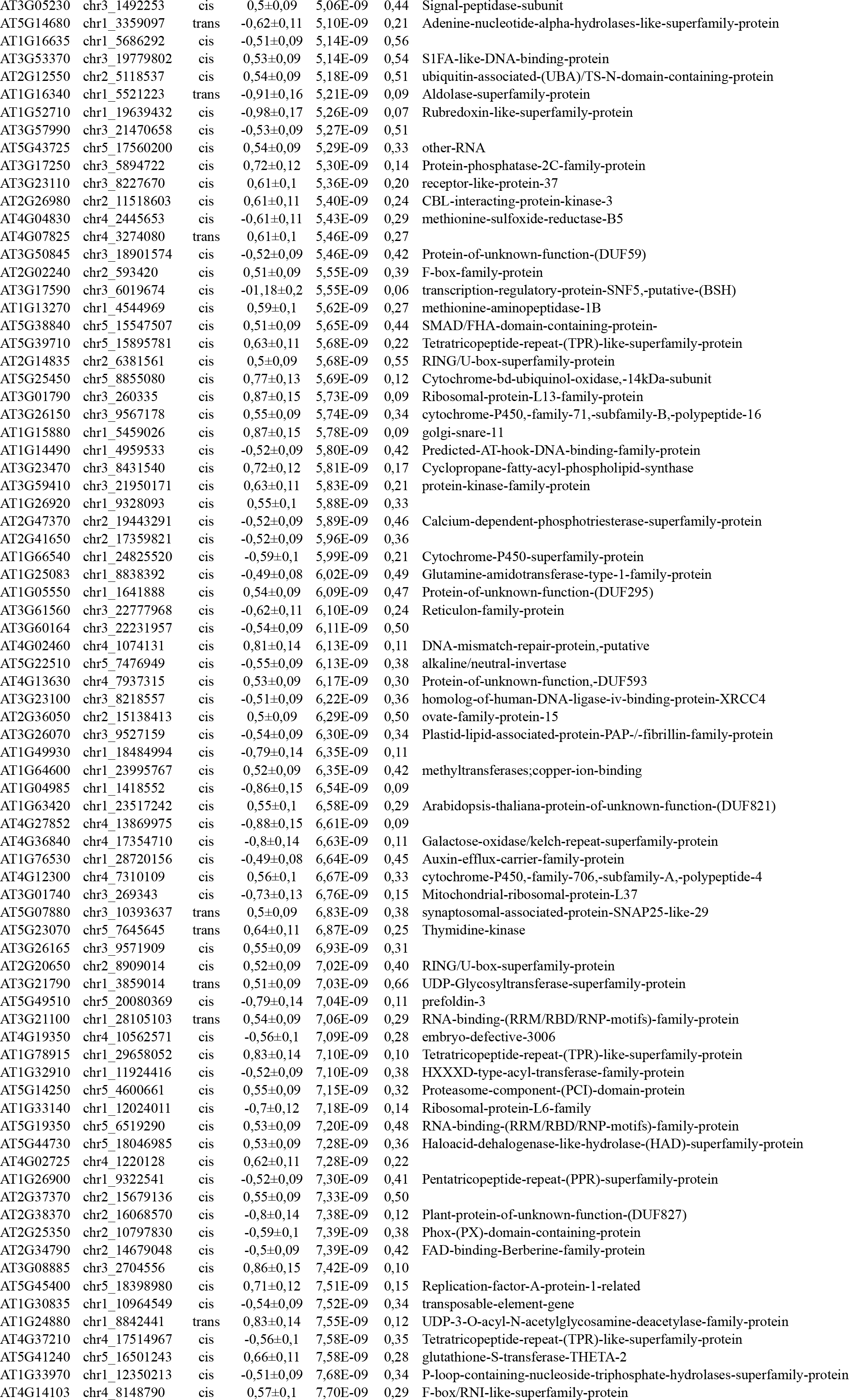
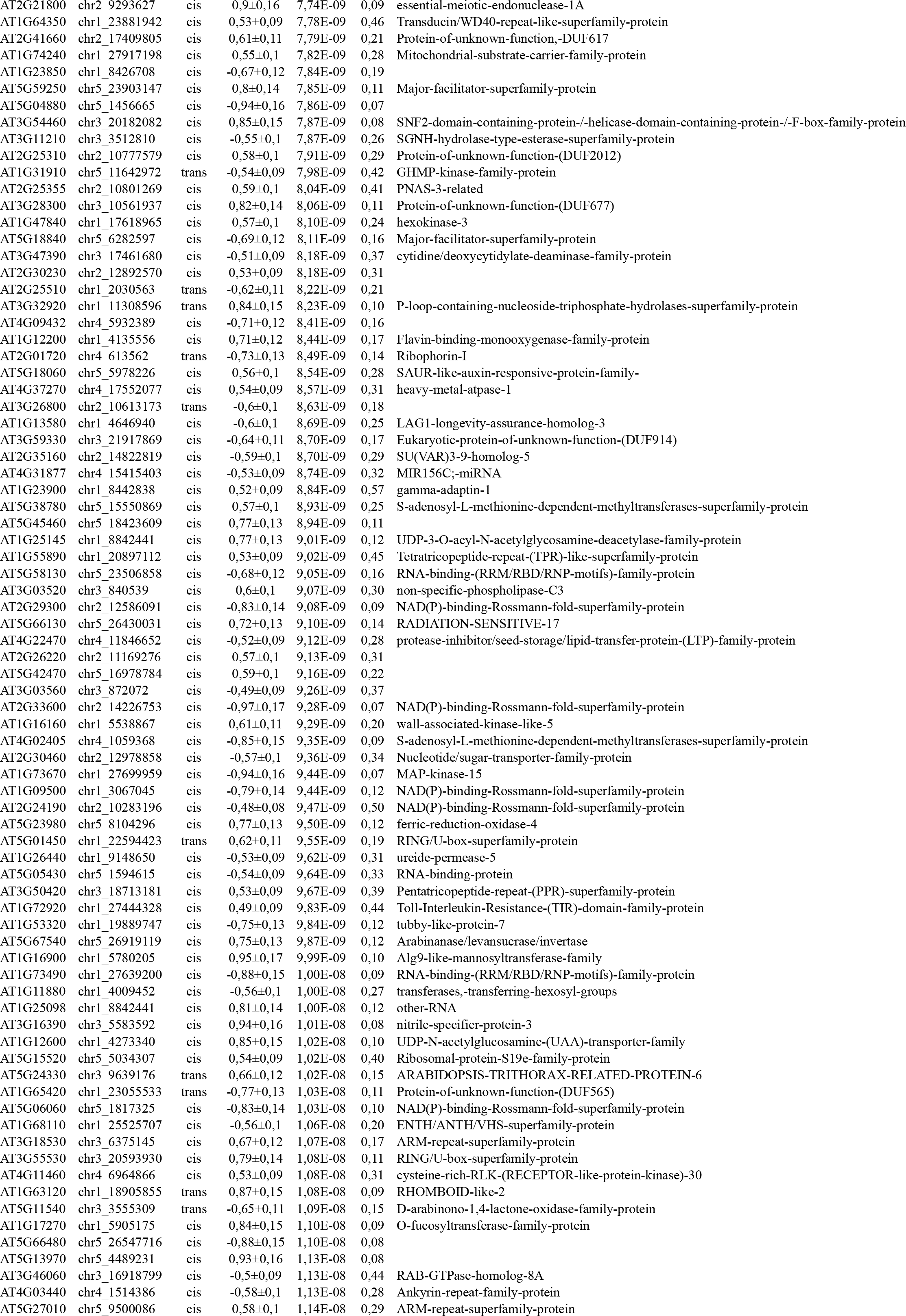
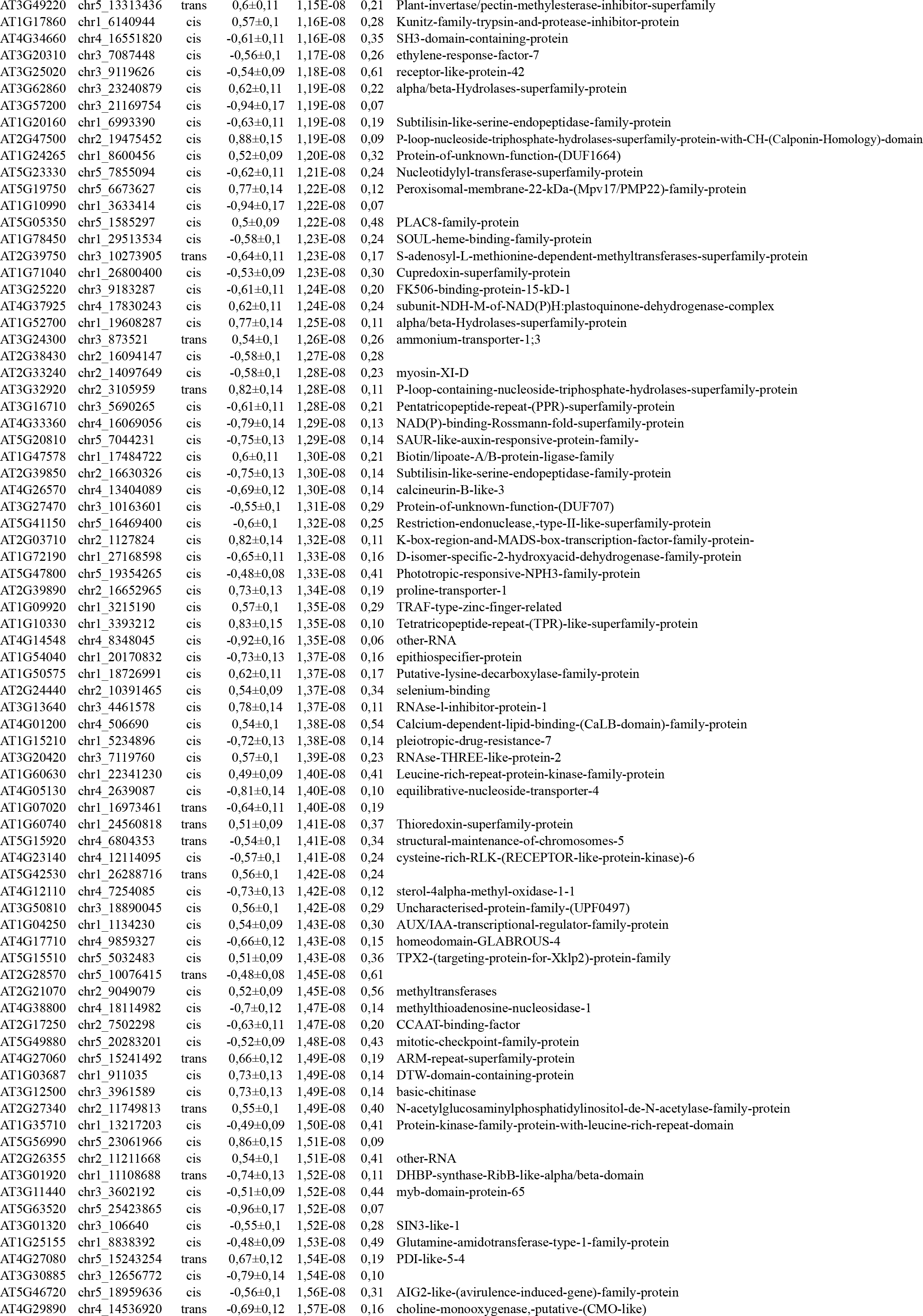
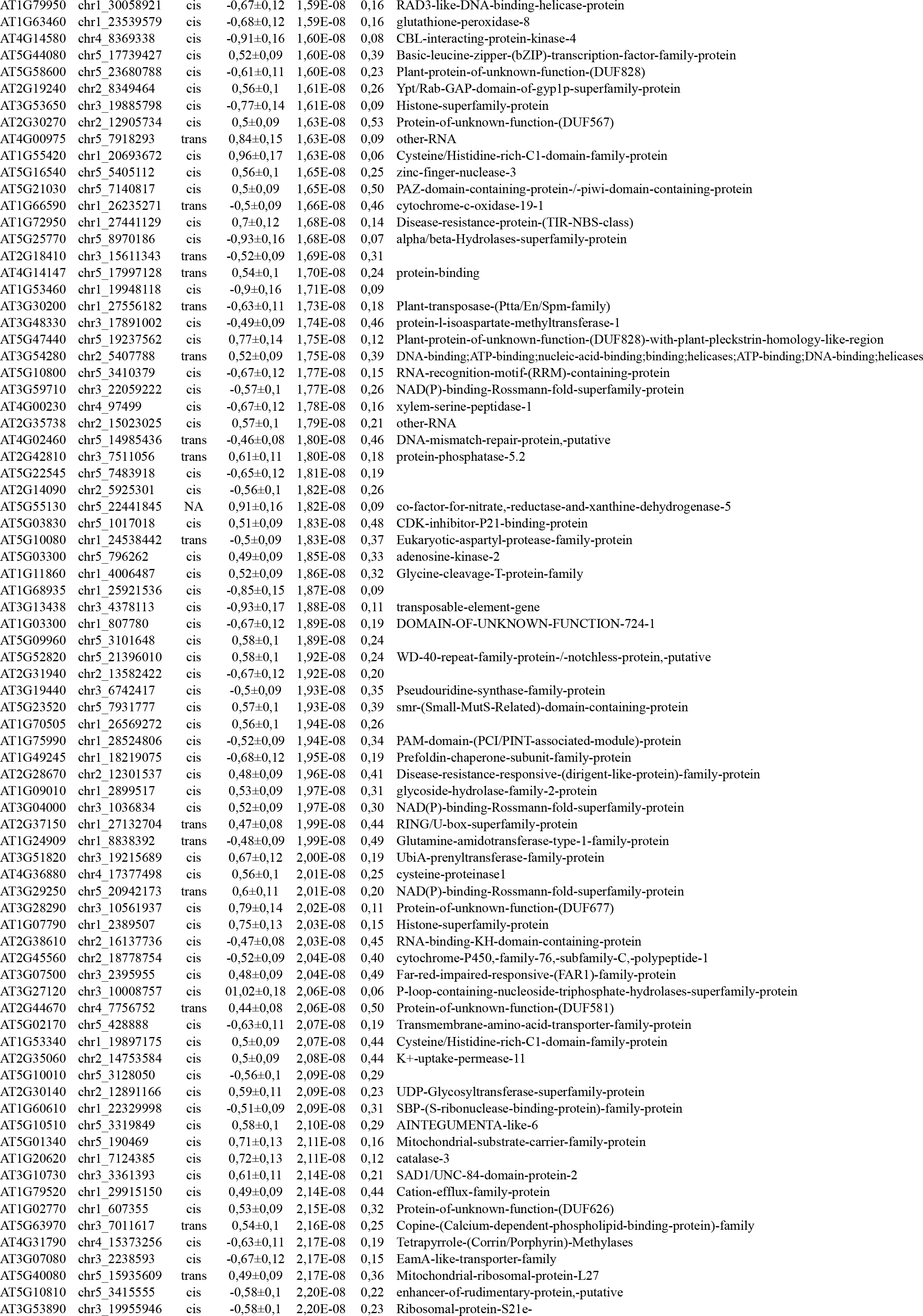
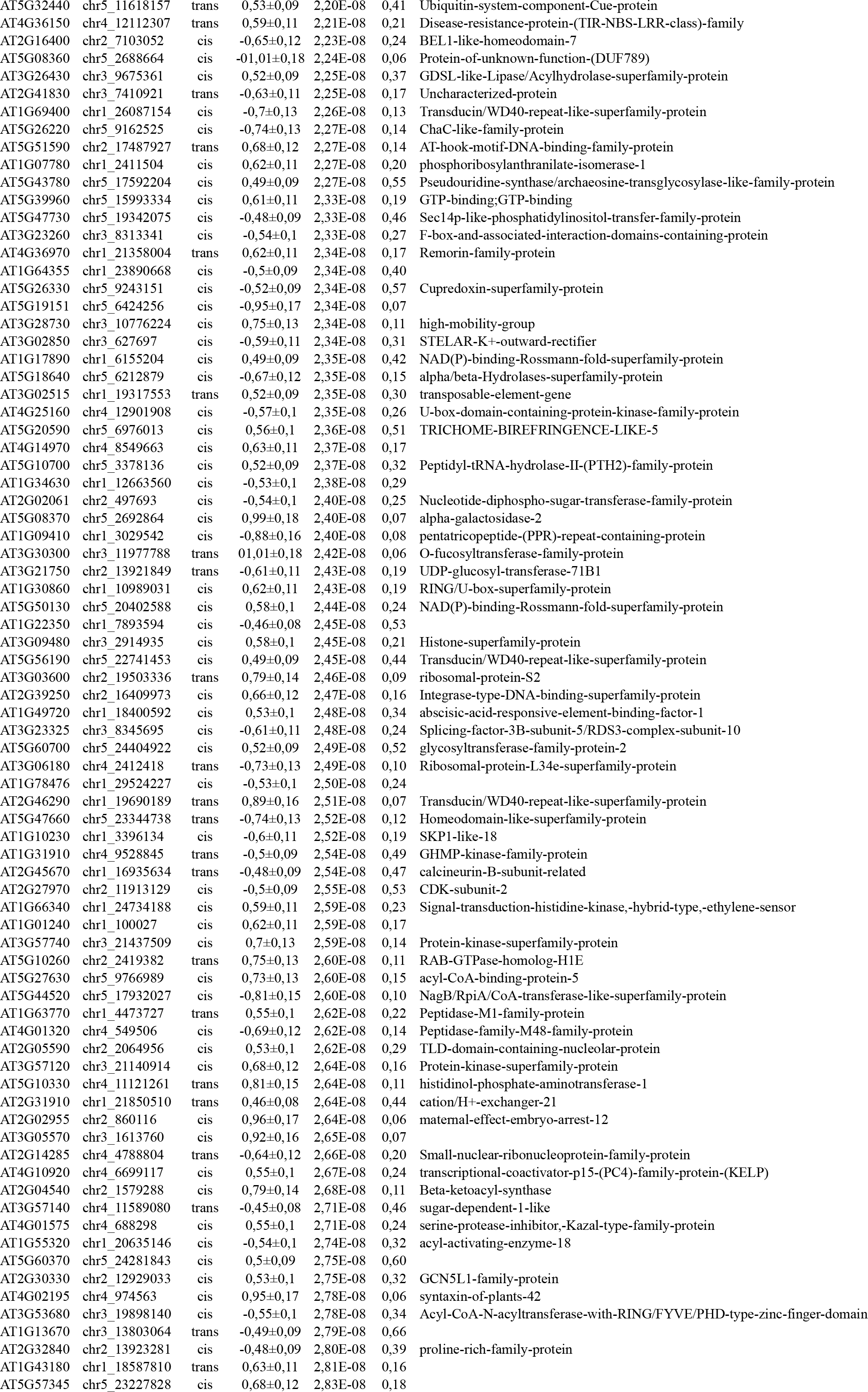
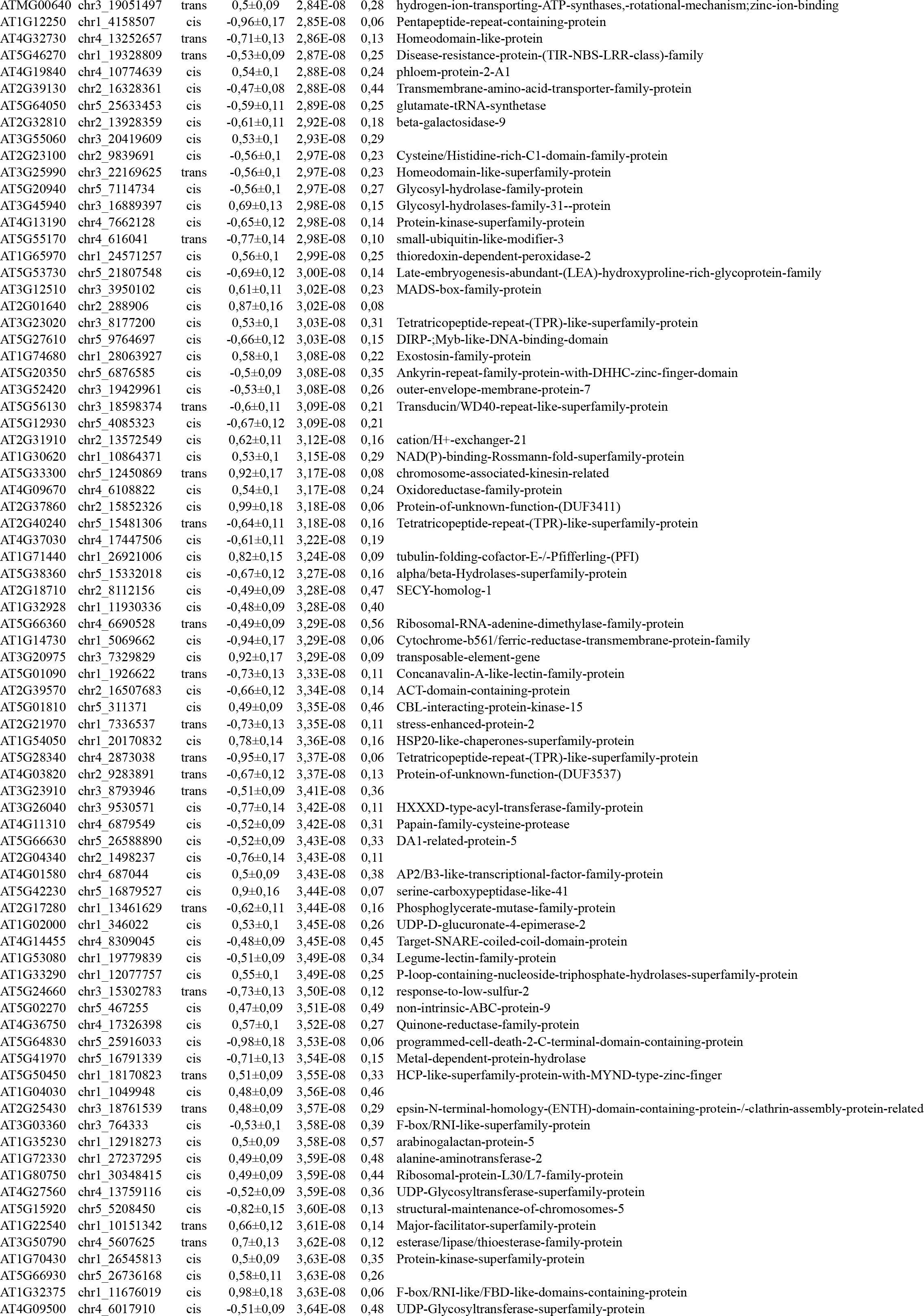
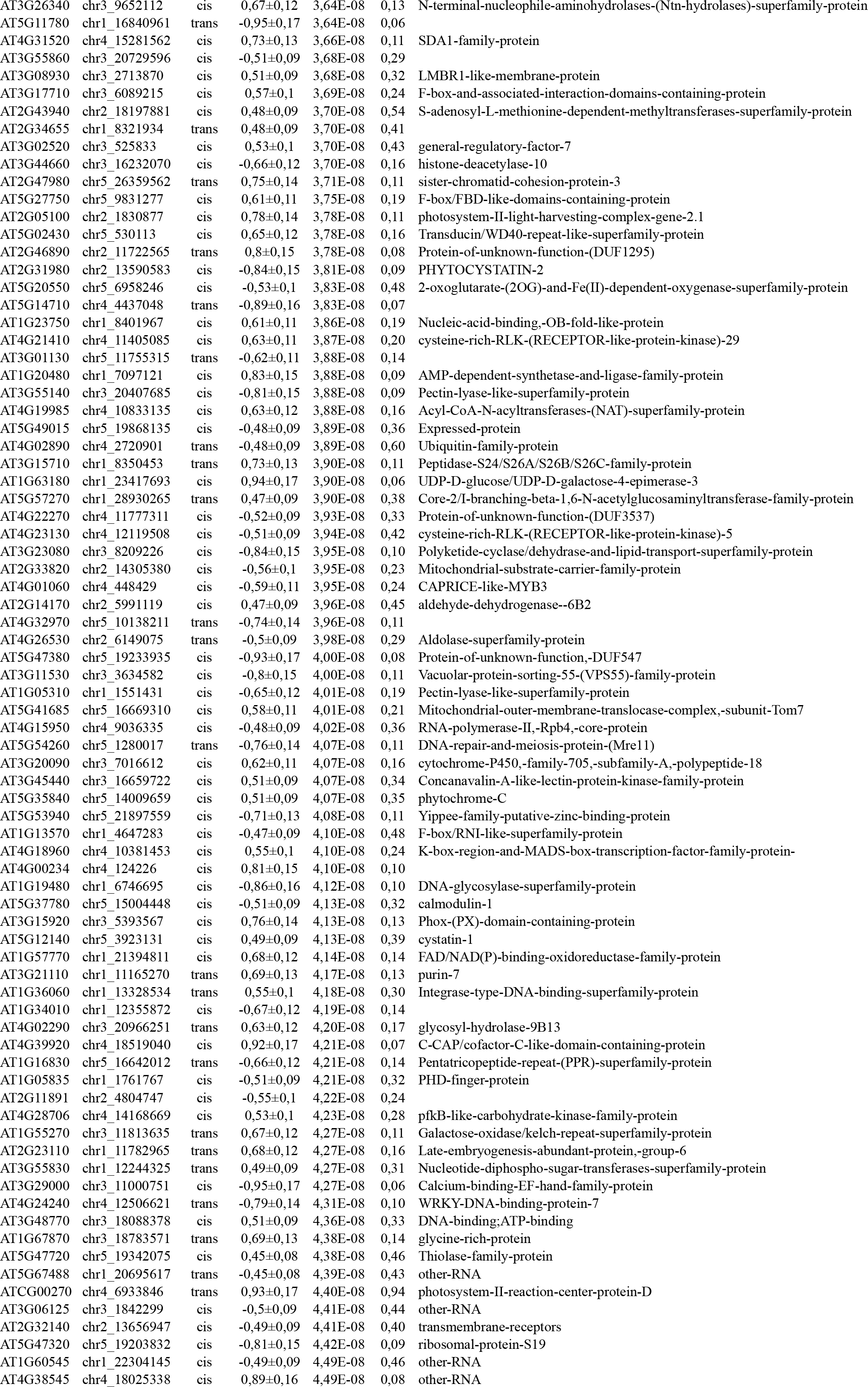
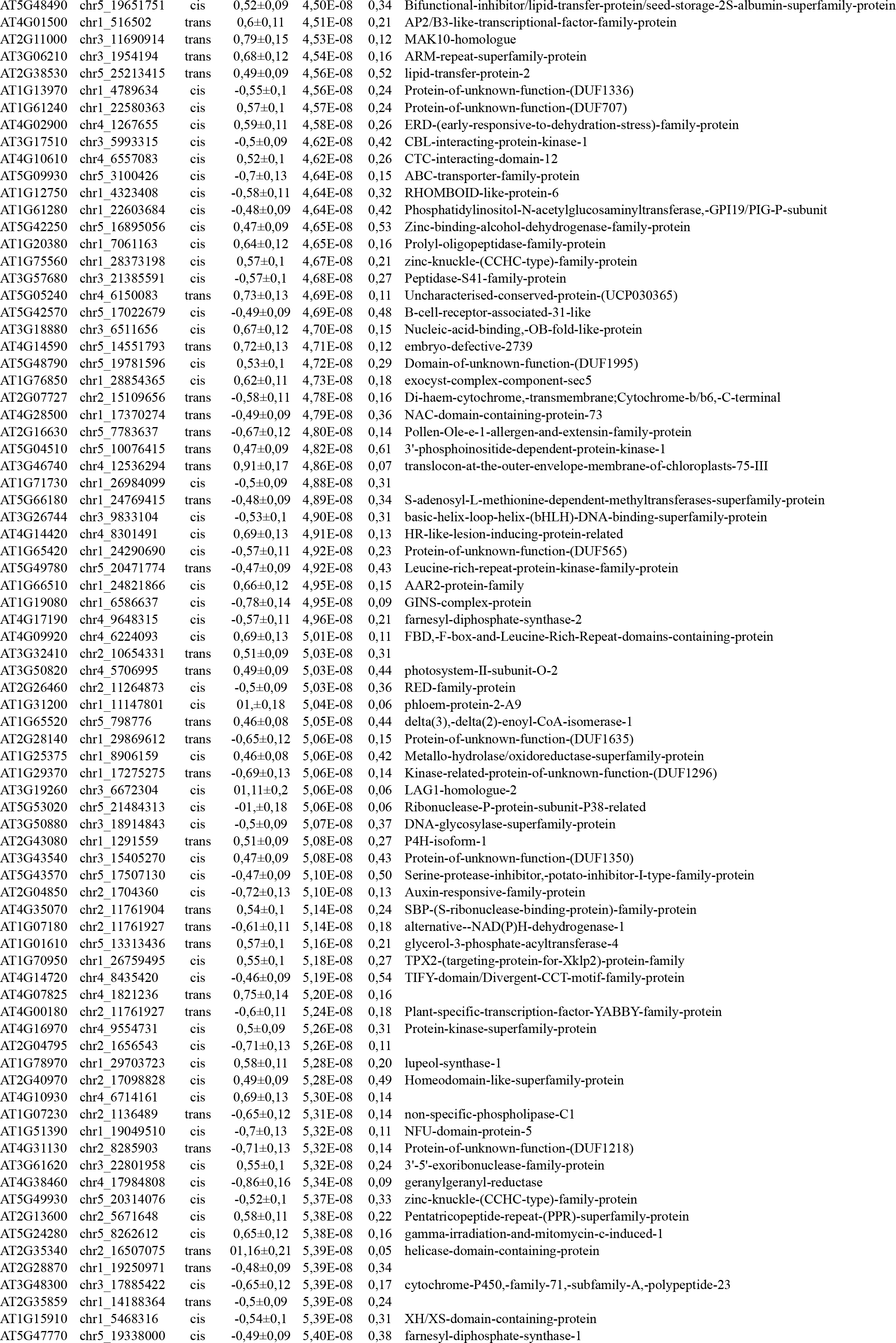
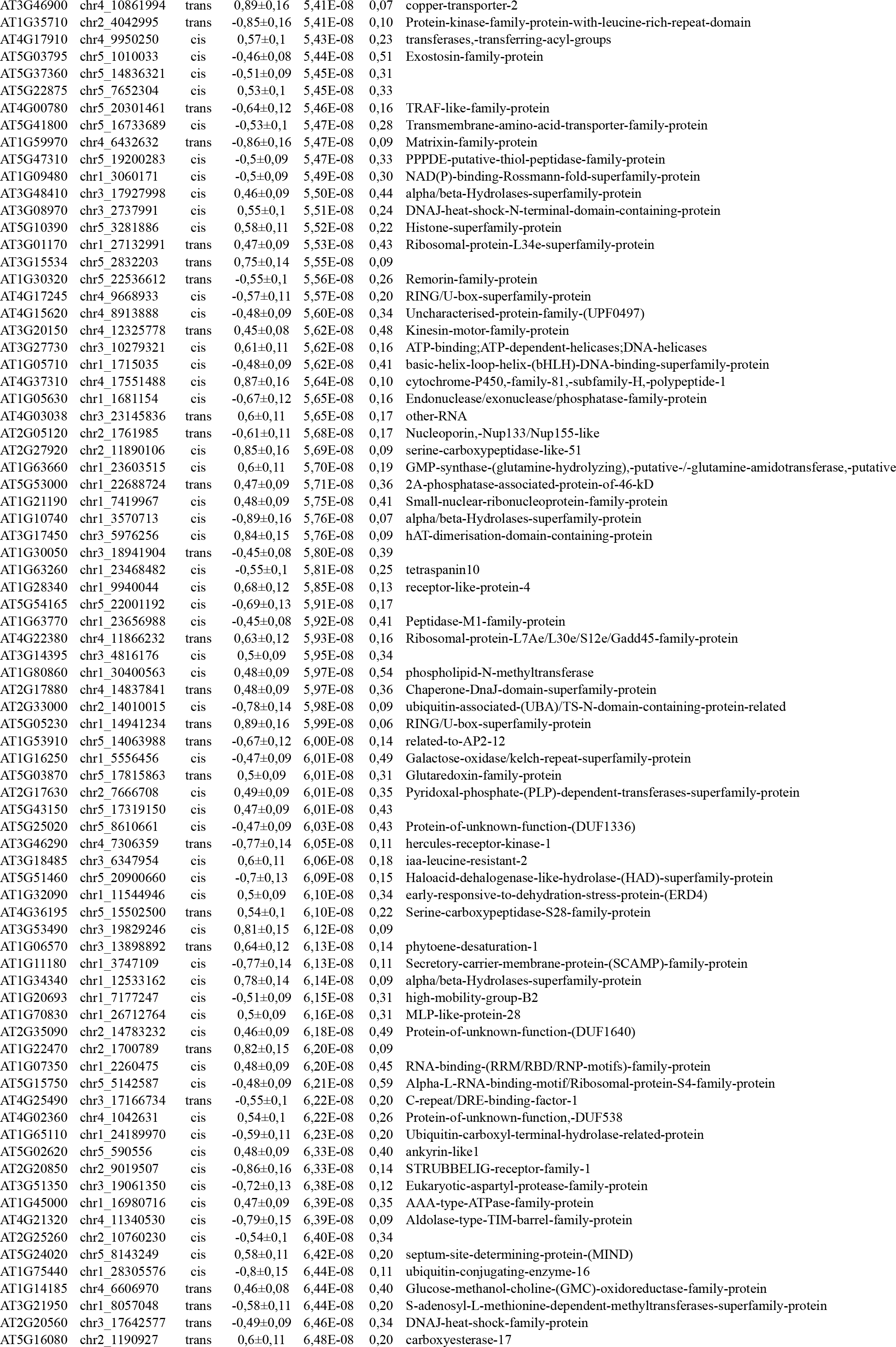
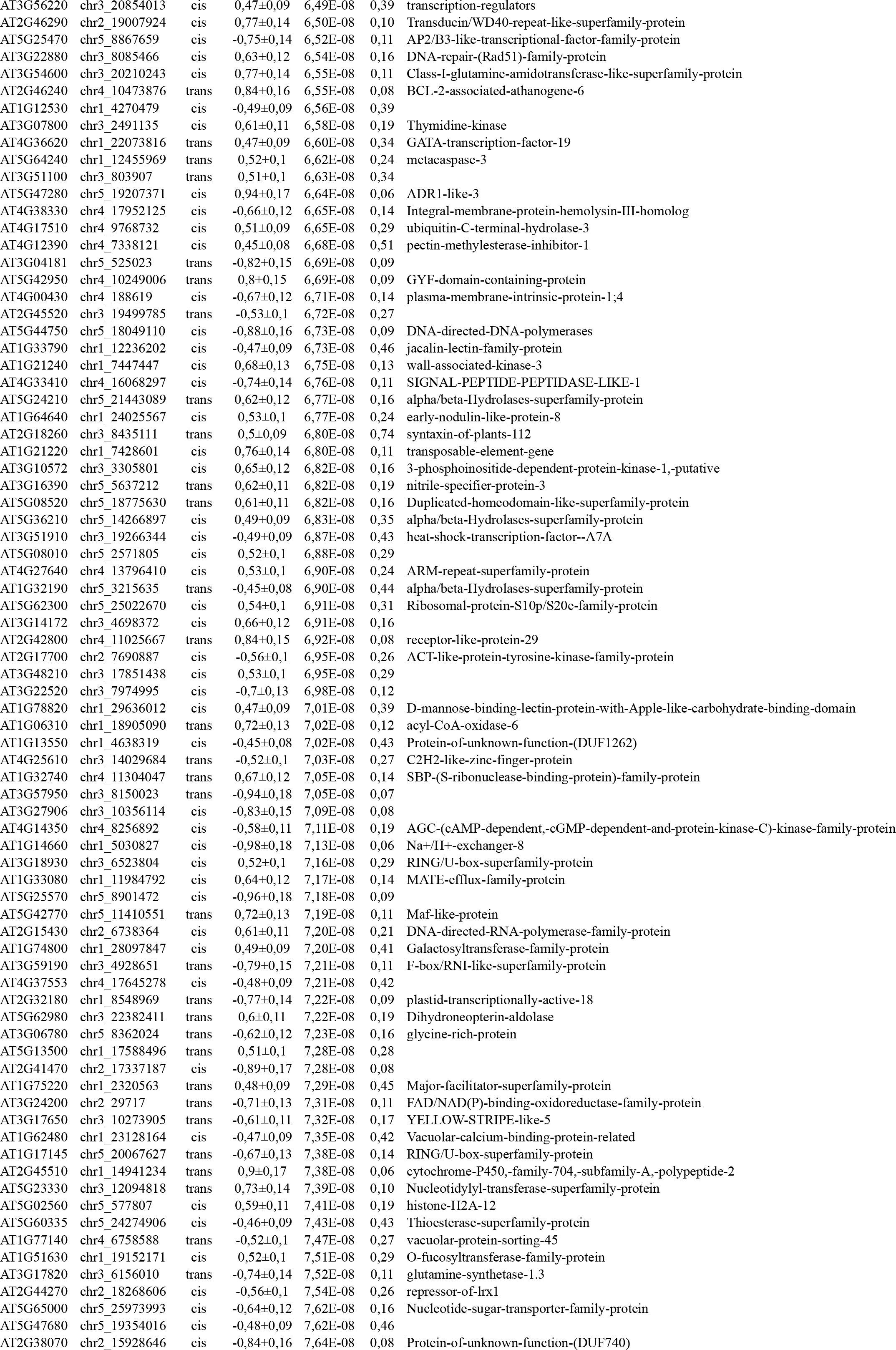
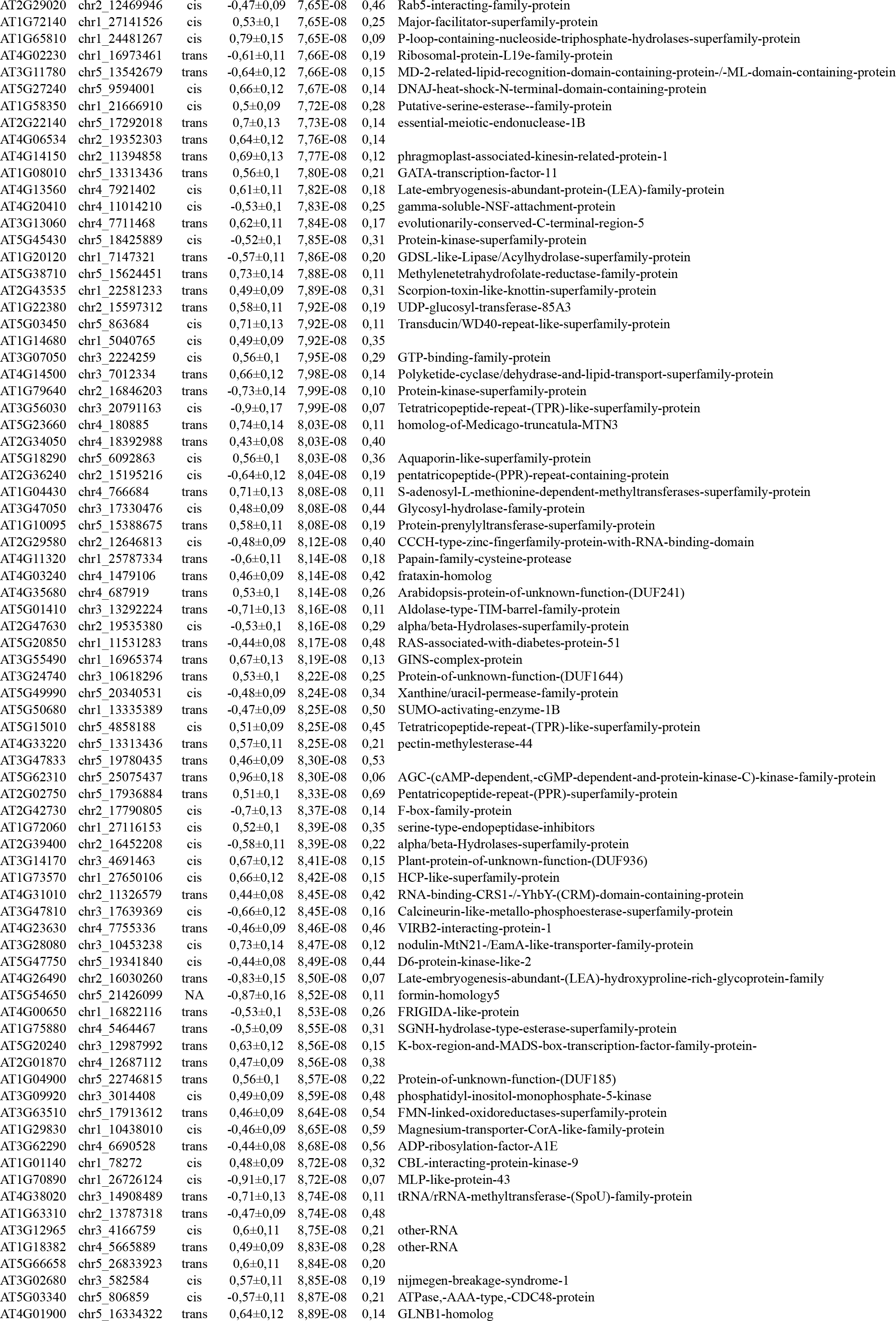
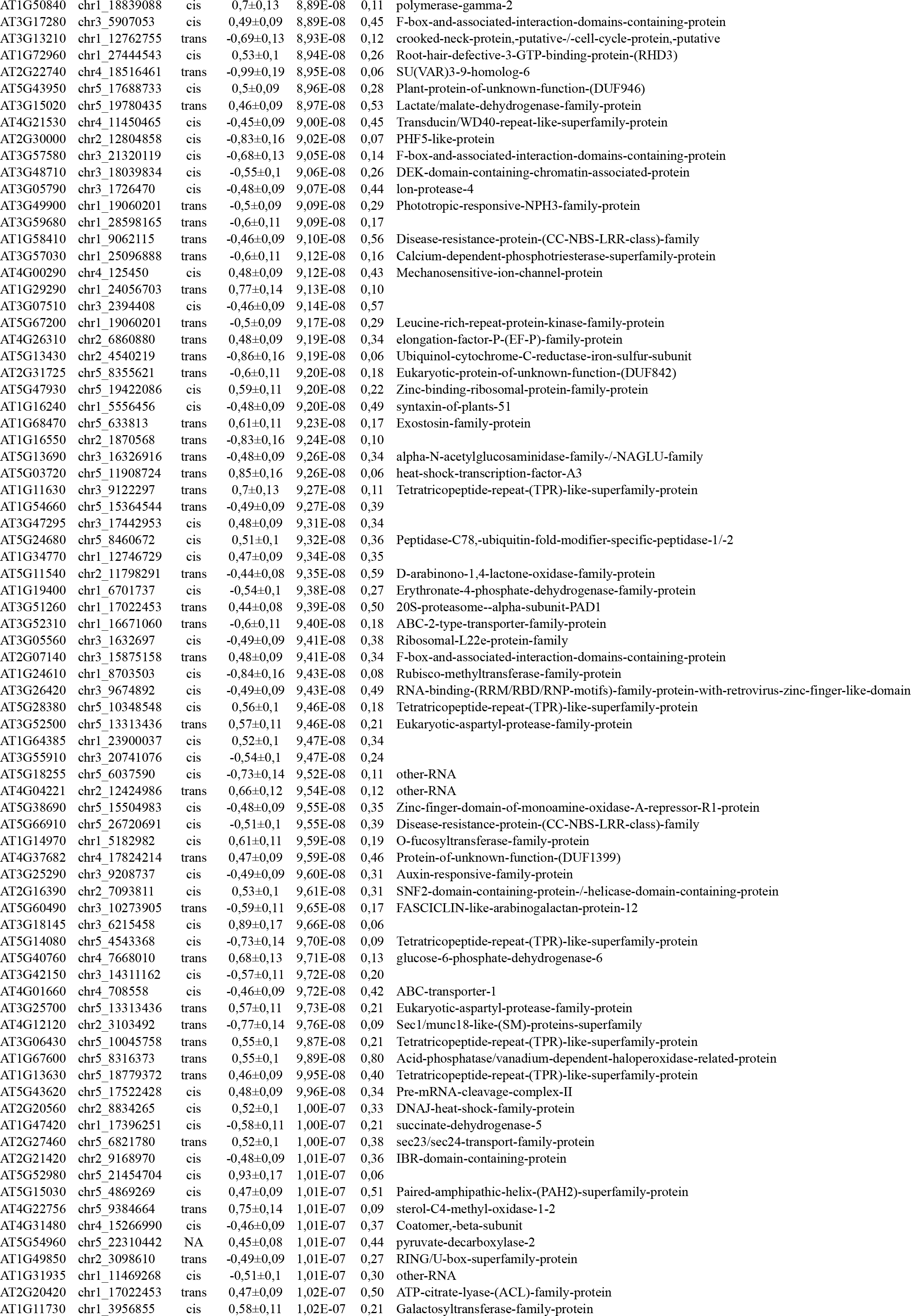
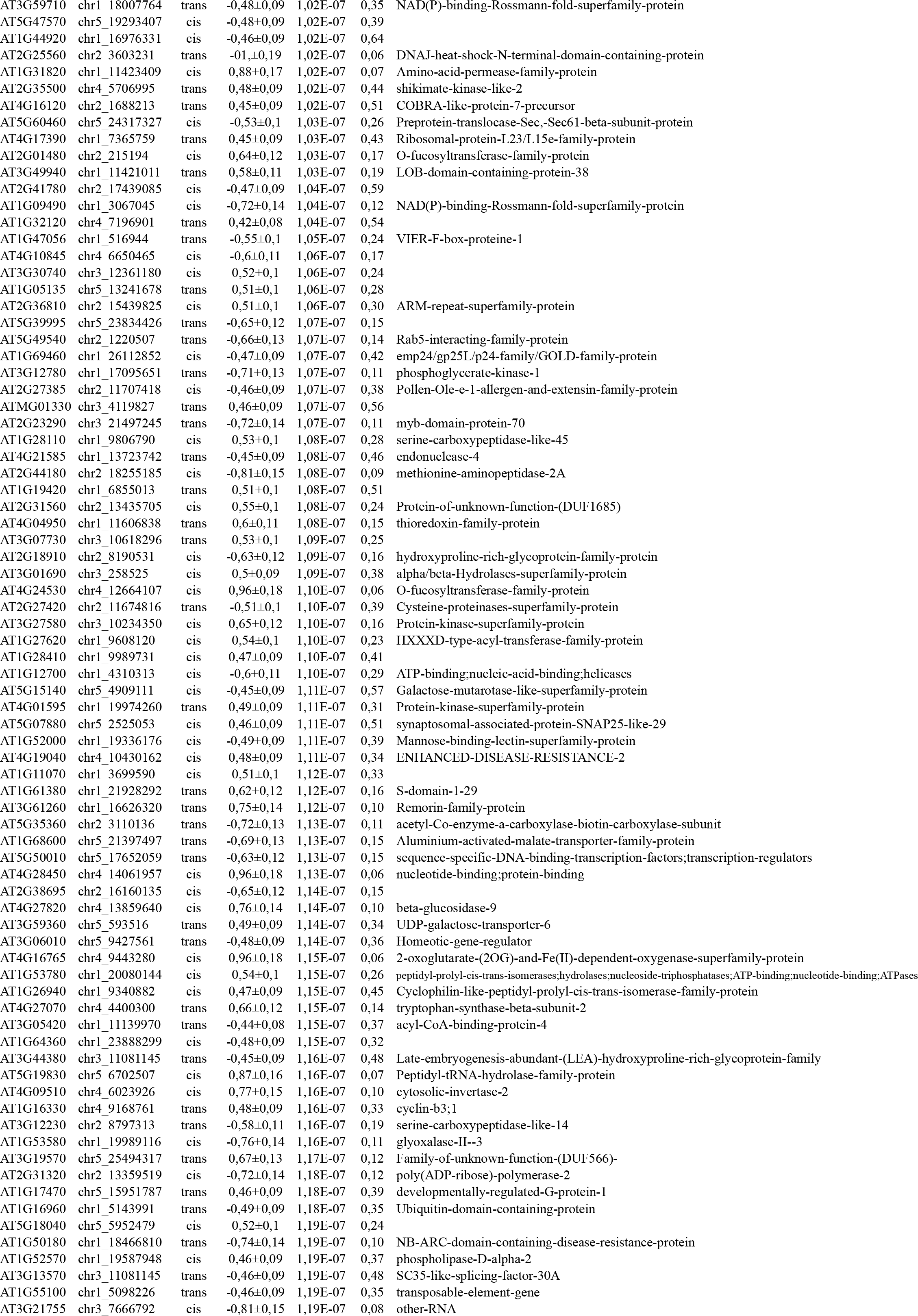
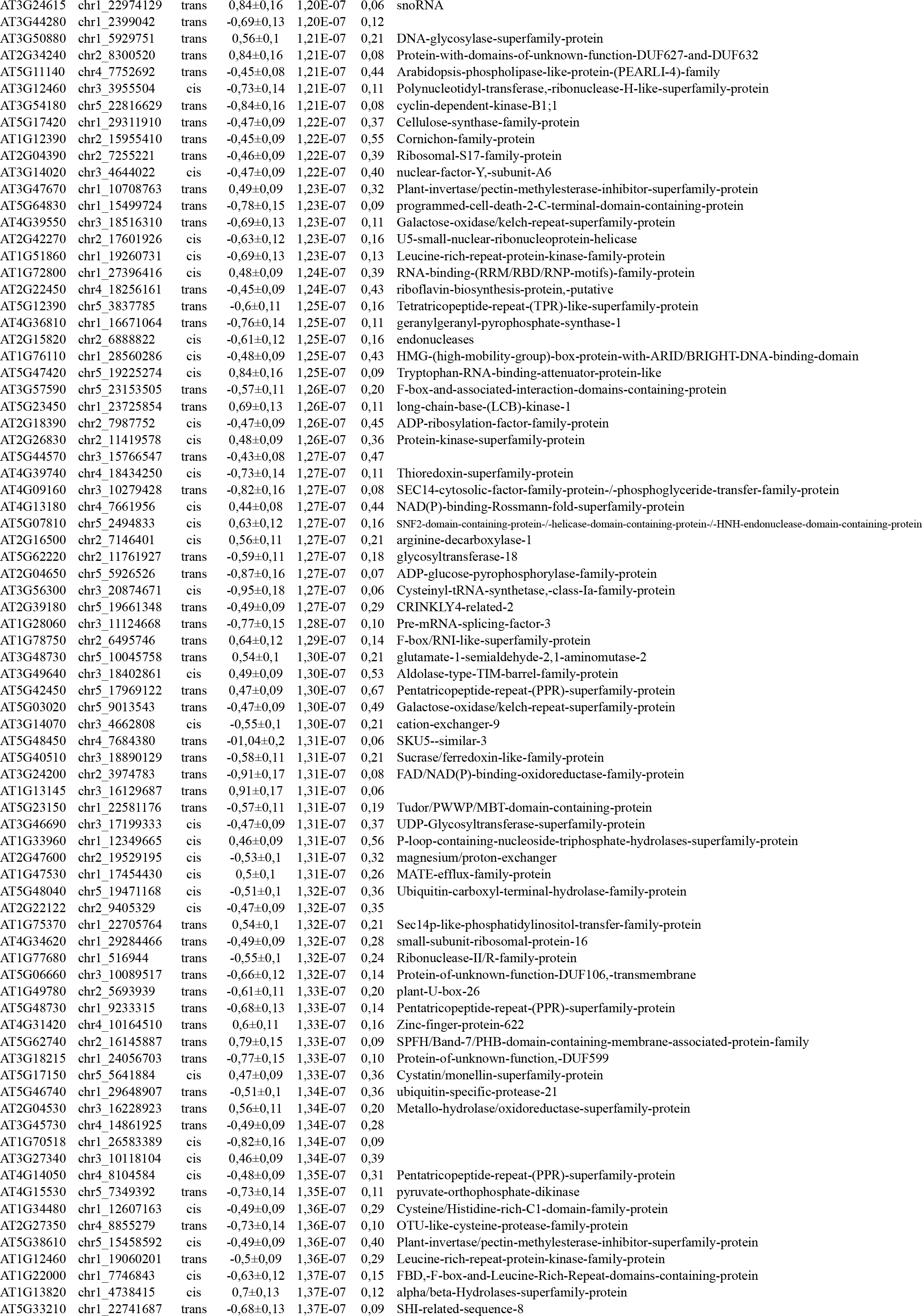
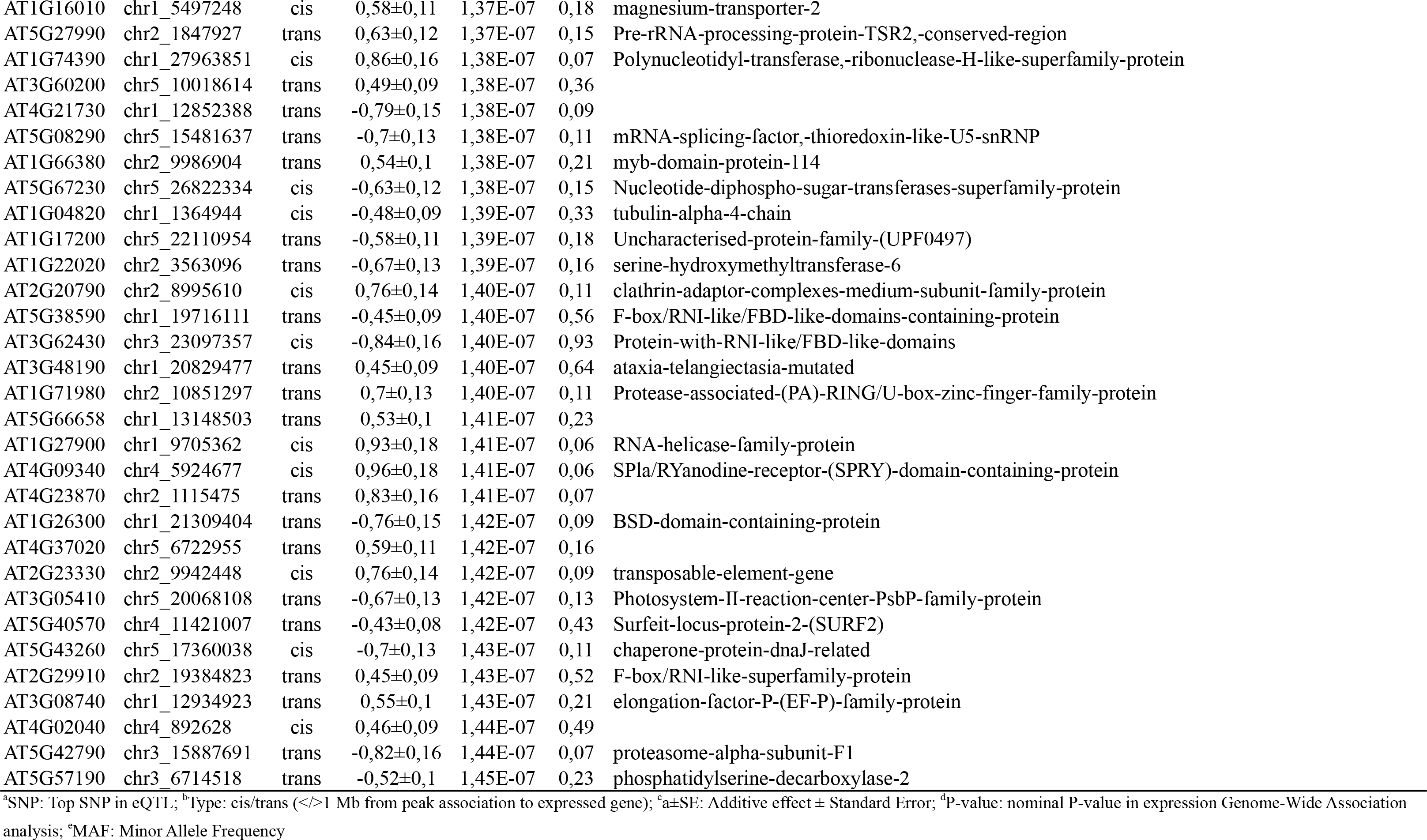
2,320 eQTL regulating the expression of 2,240 genes expressed in most of the 140 natural *A. thaliana* accessions

**Table S5.**
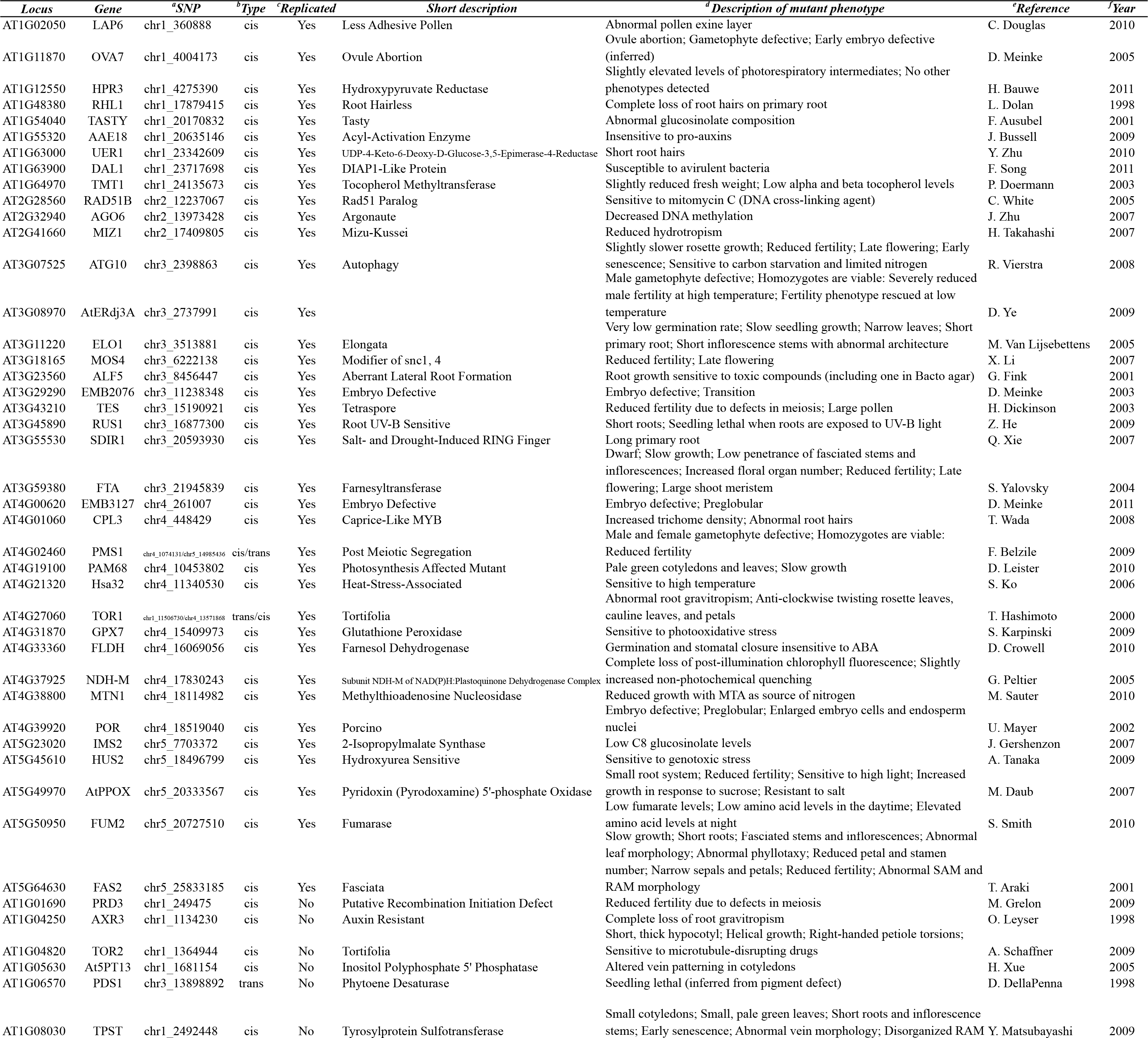
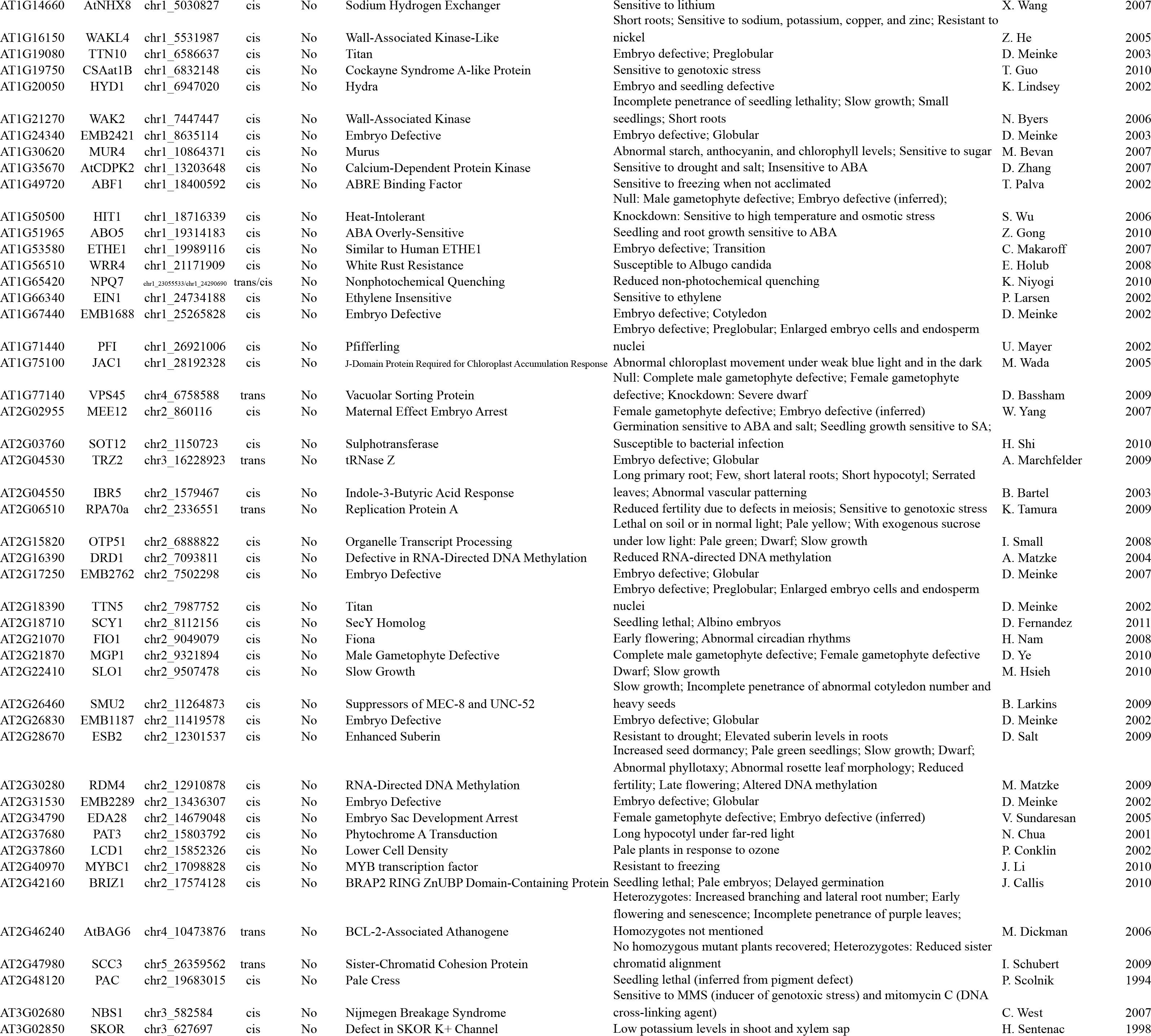
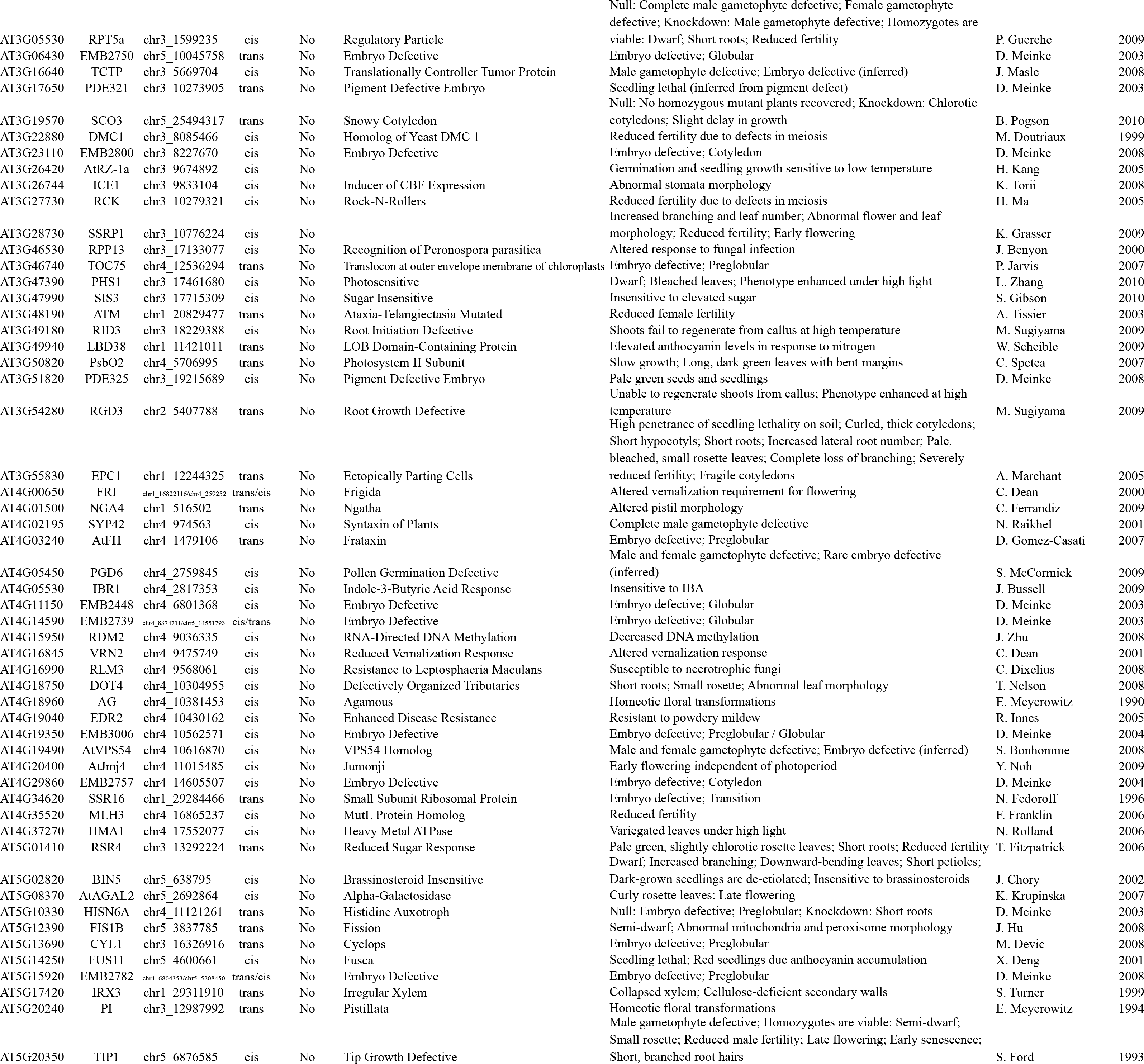
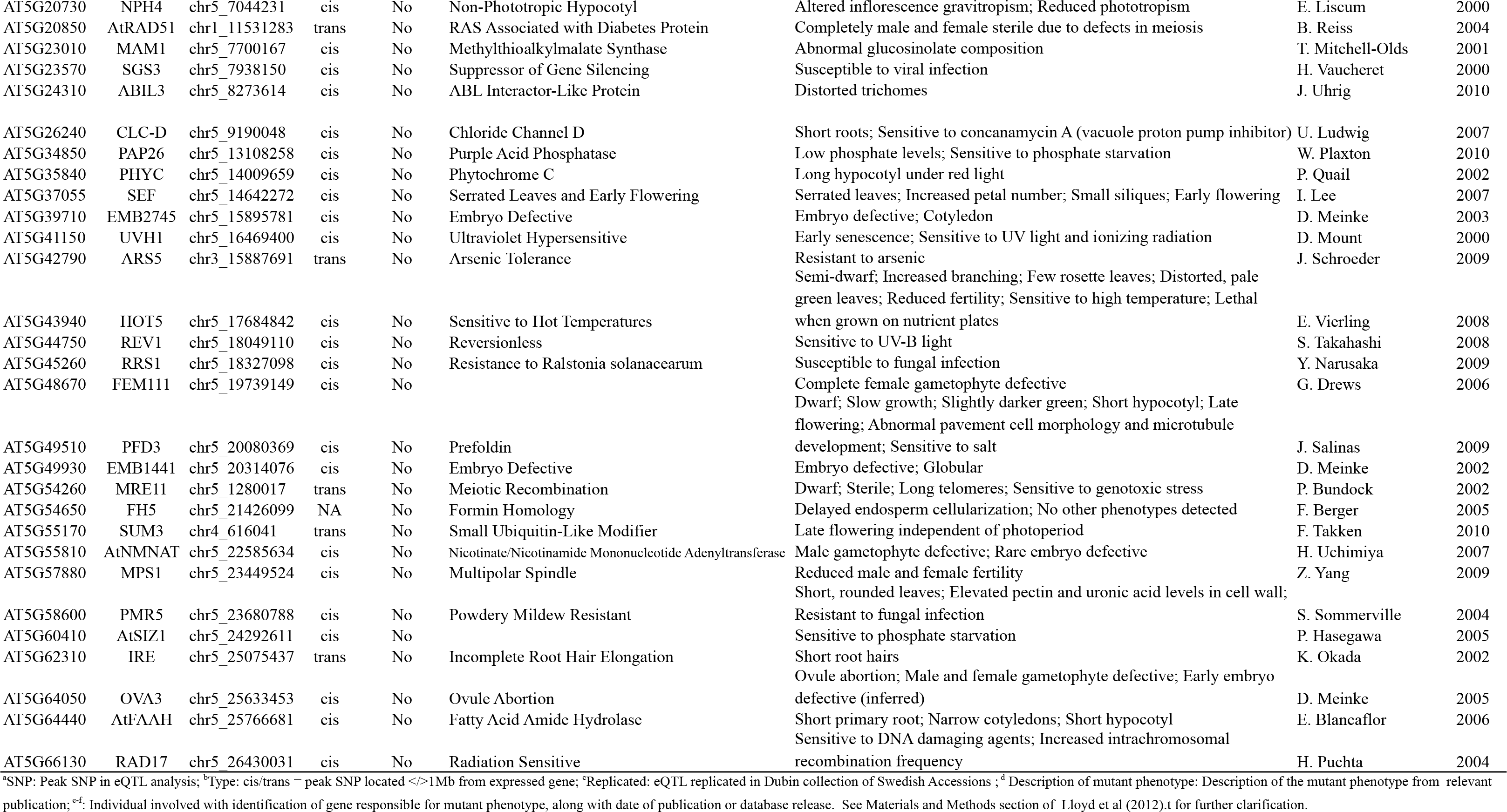
175 genes affected by eQTL in the population of 140 natural *A. thaliana* accessions for which strong phenotypic effect have already been described in Lloyd et al. [41].

**Table S6.**
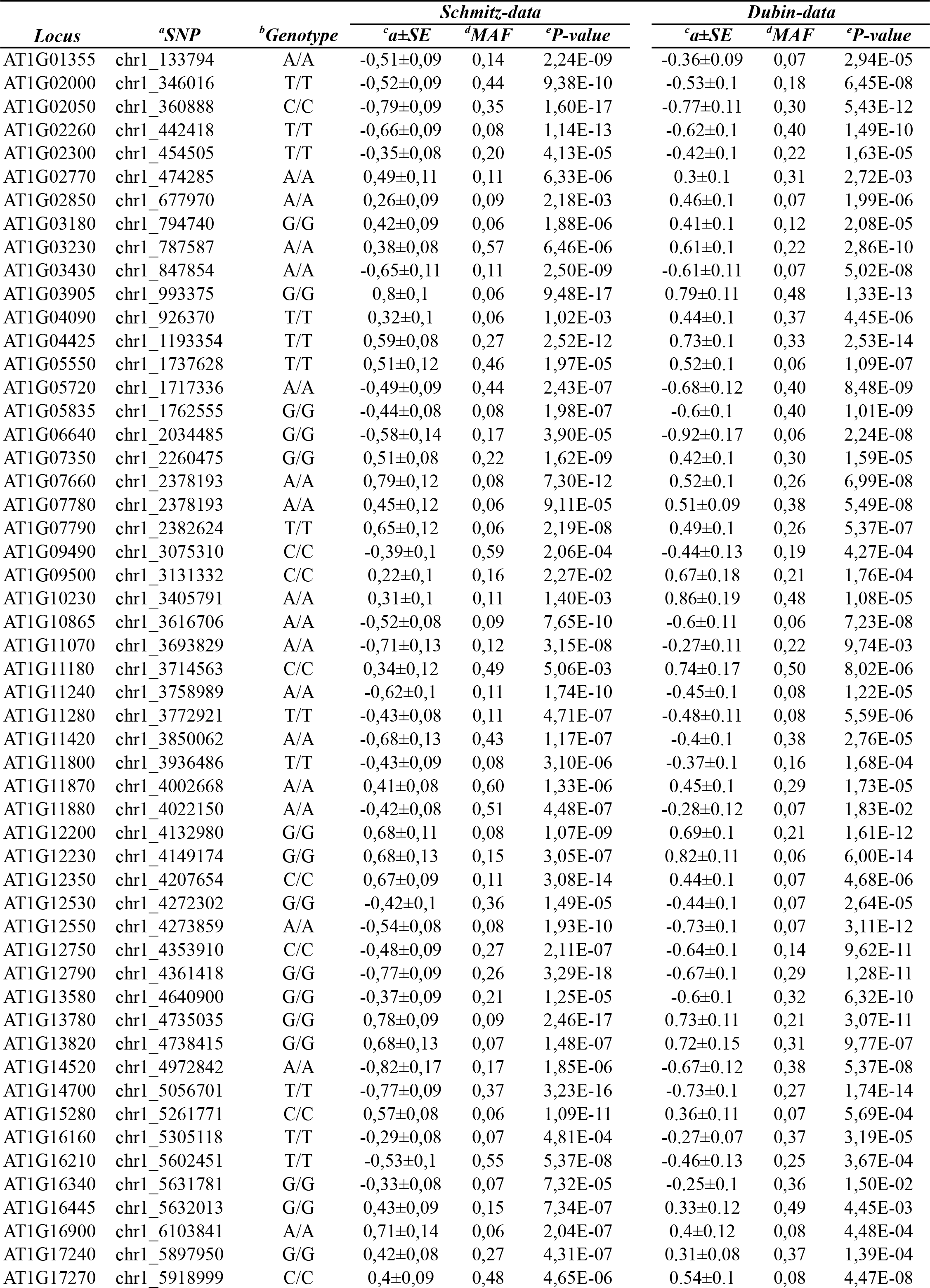
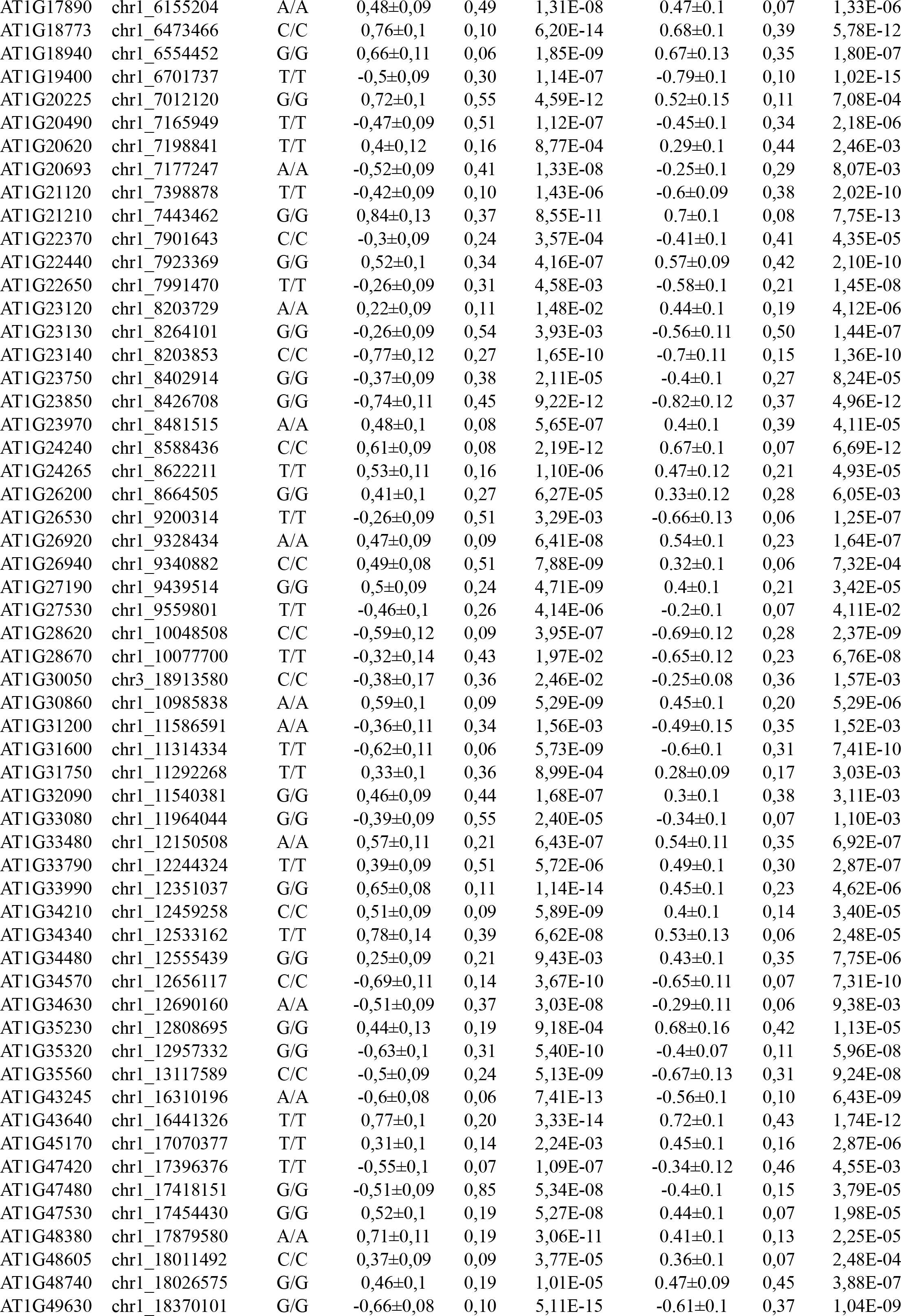
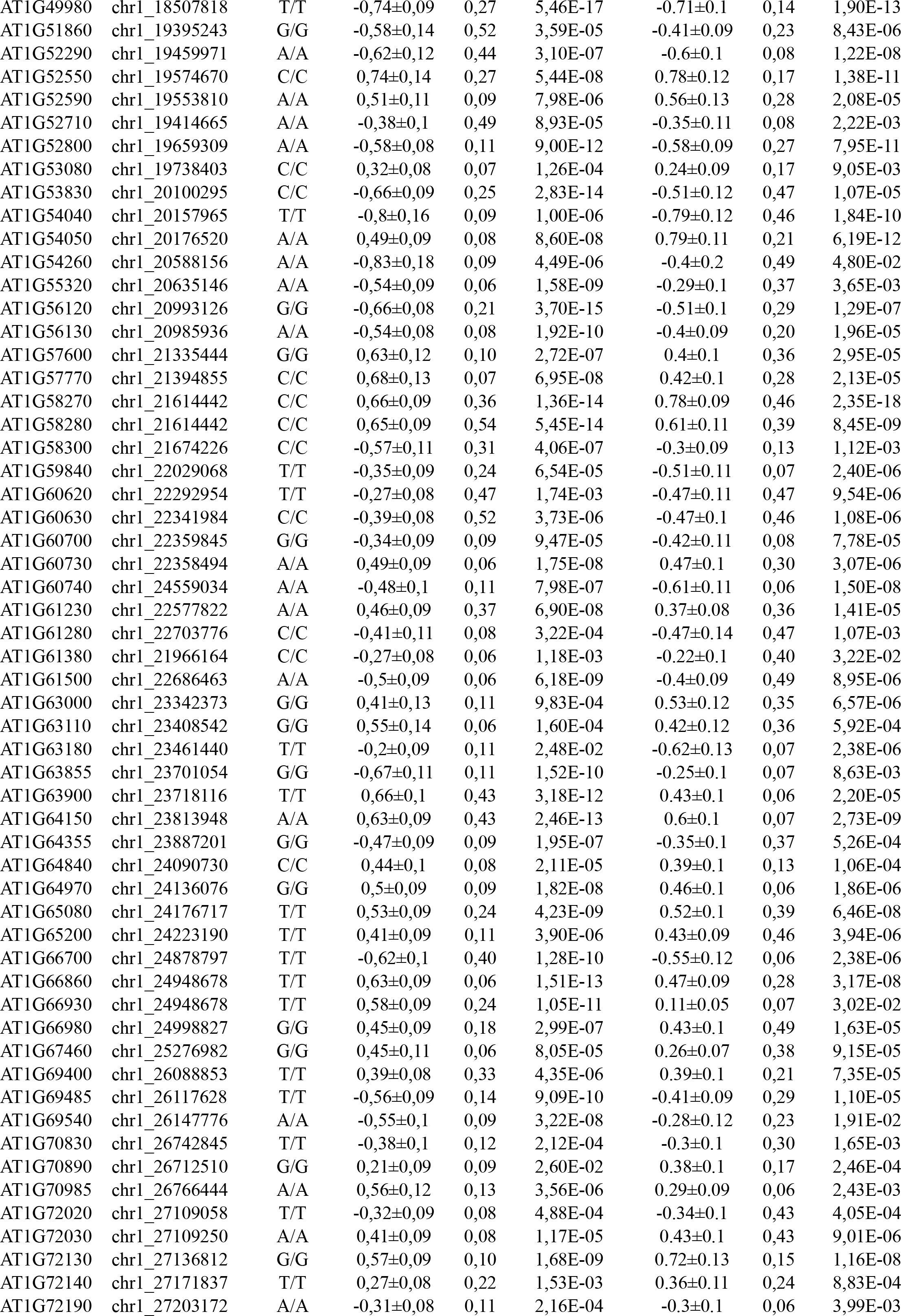
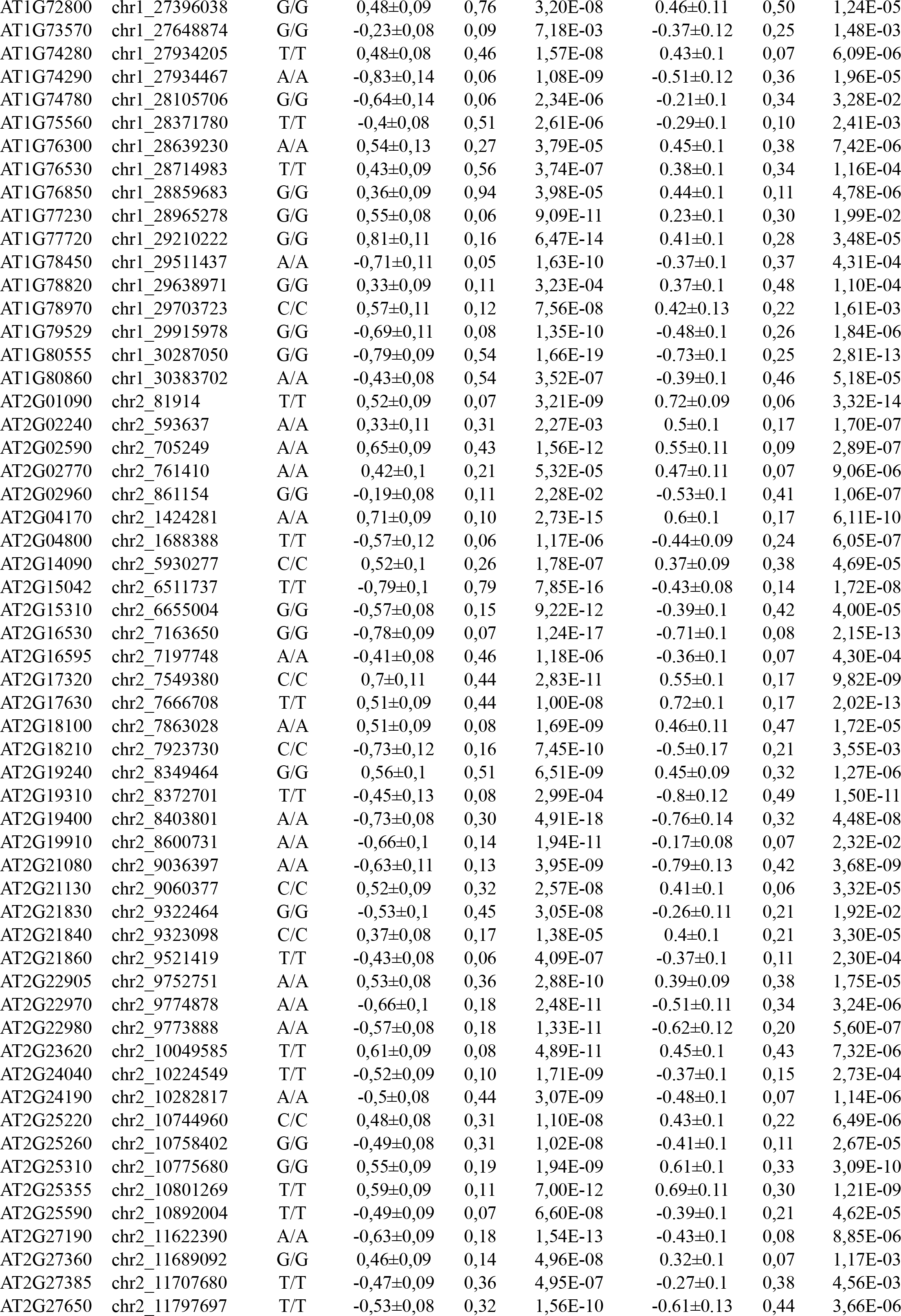
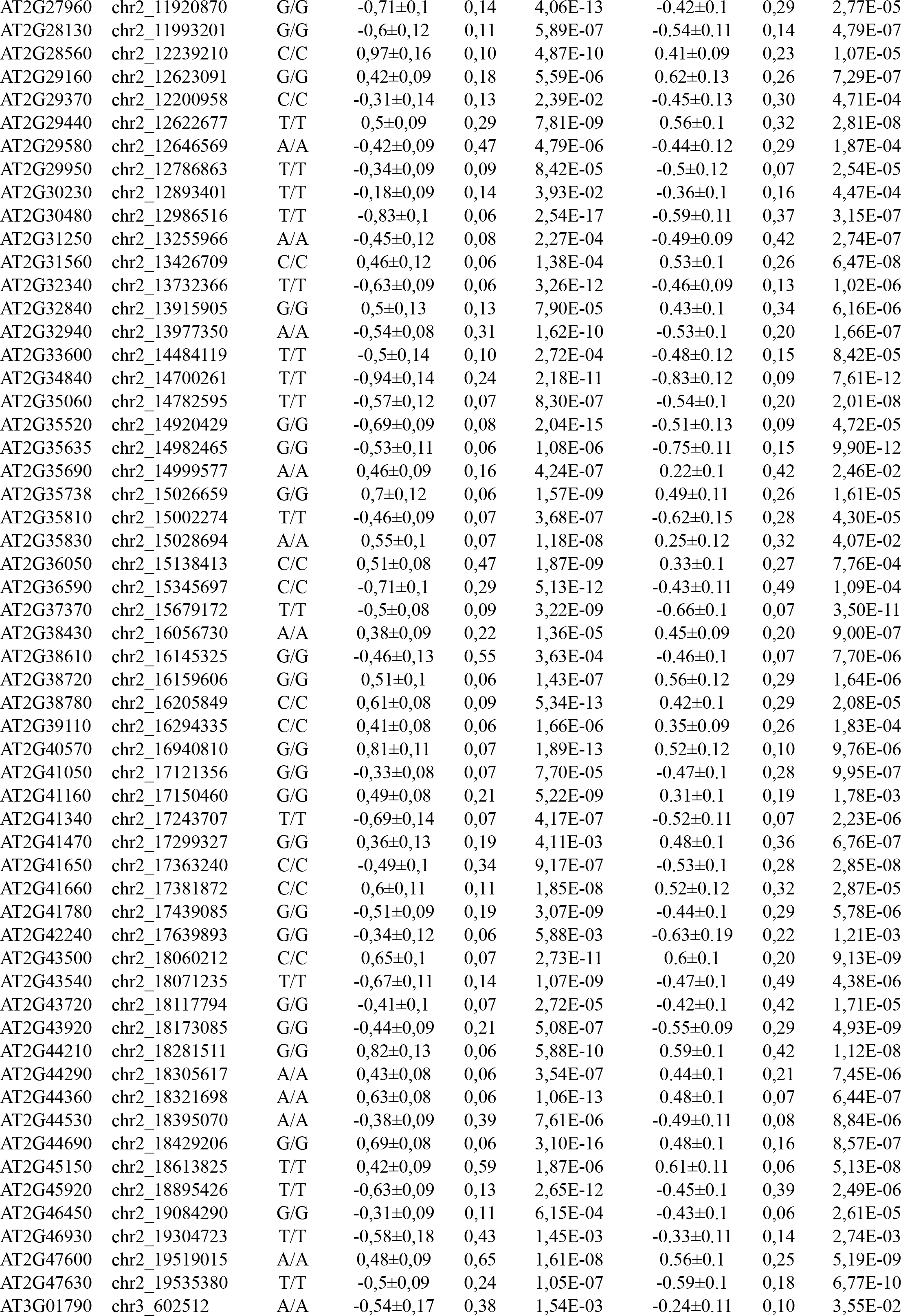
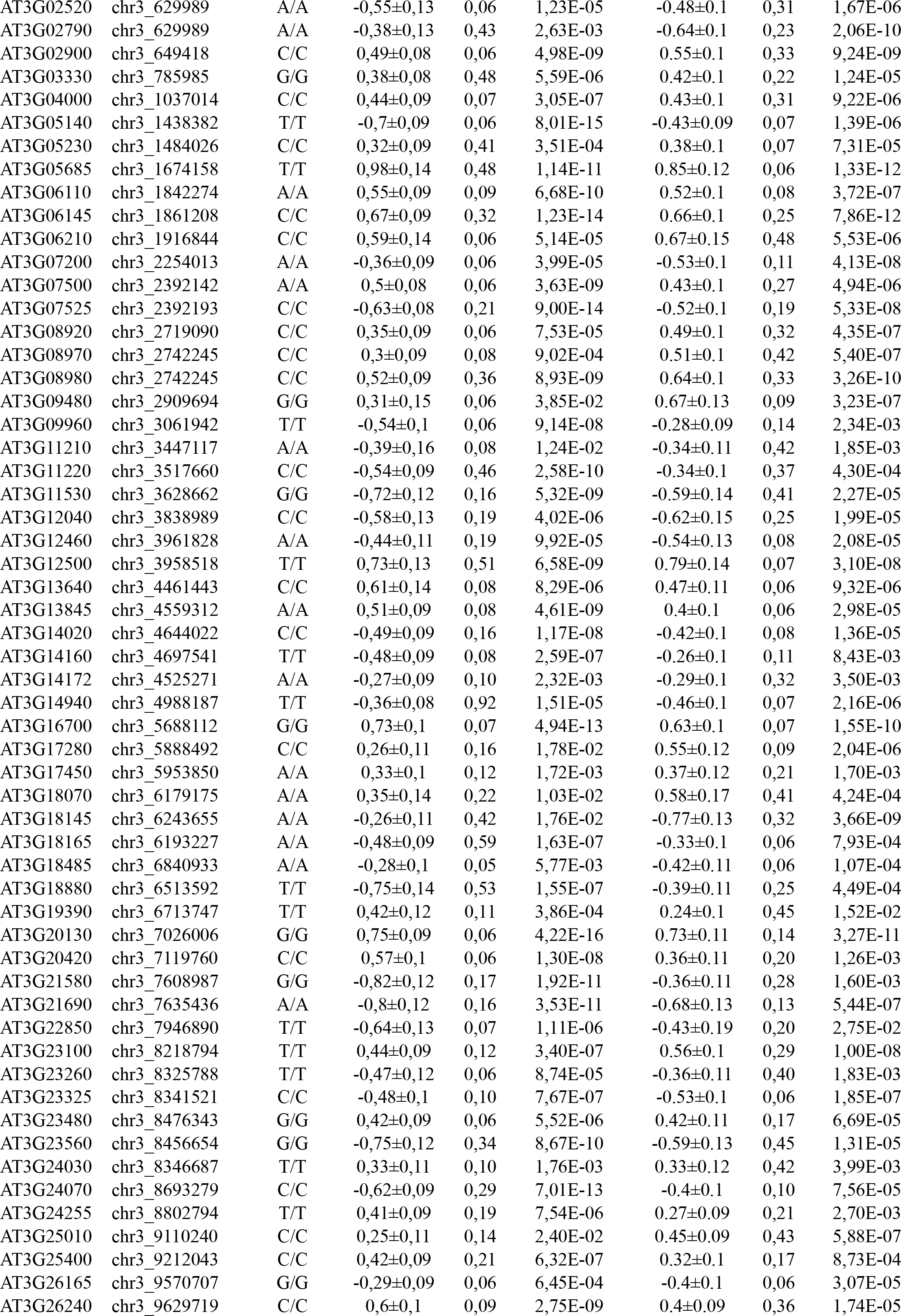
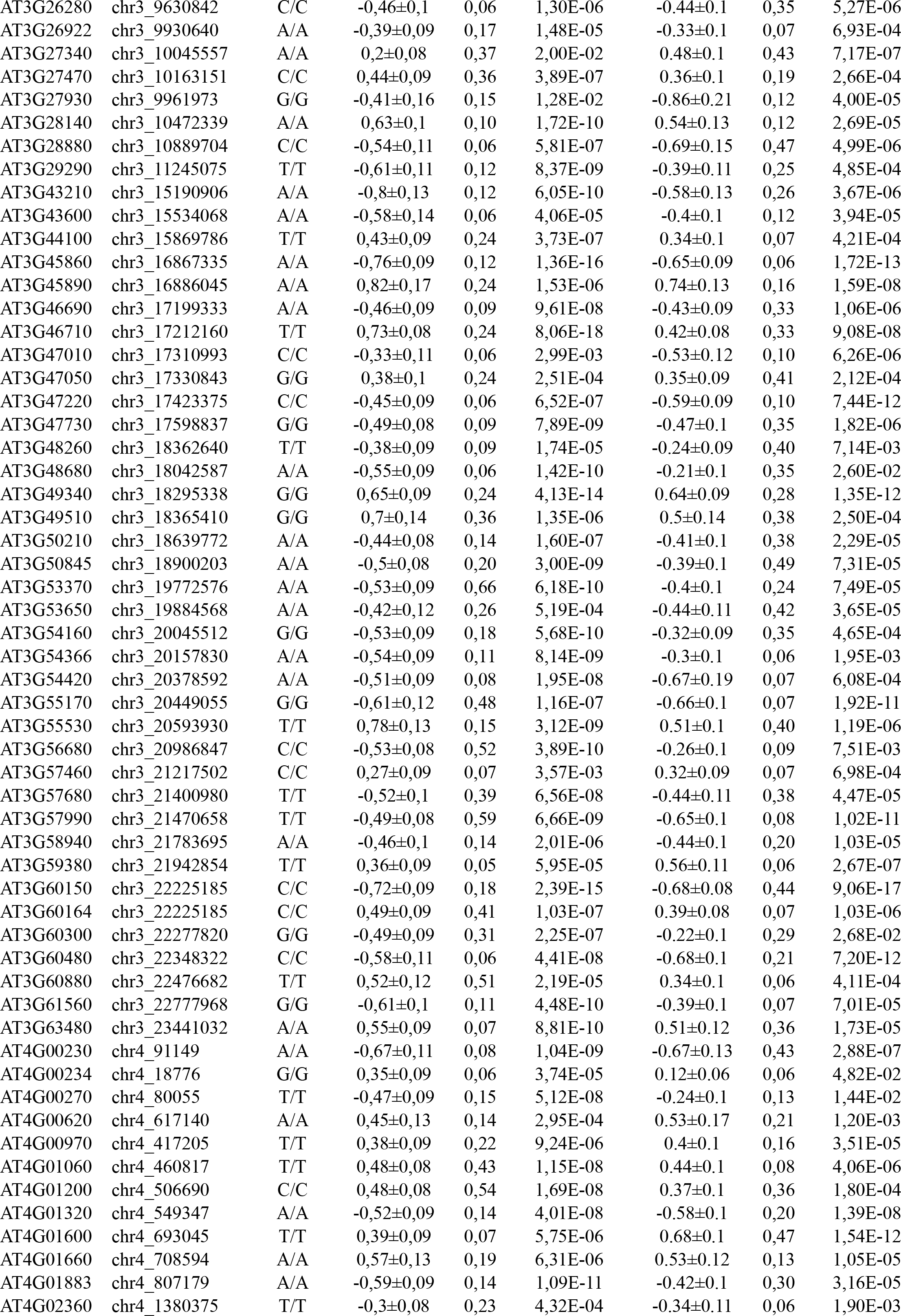
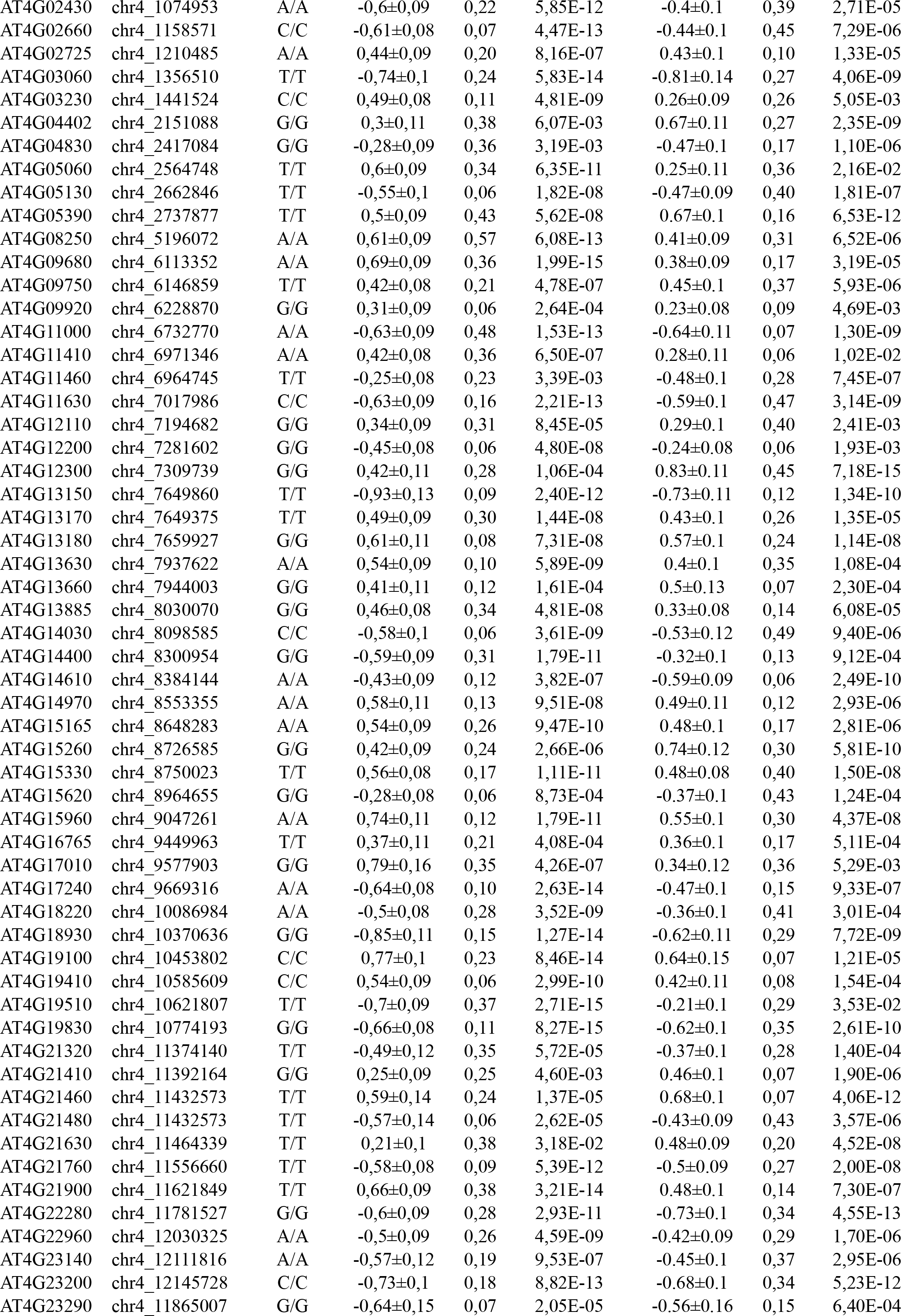
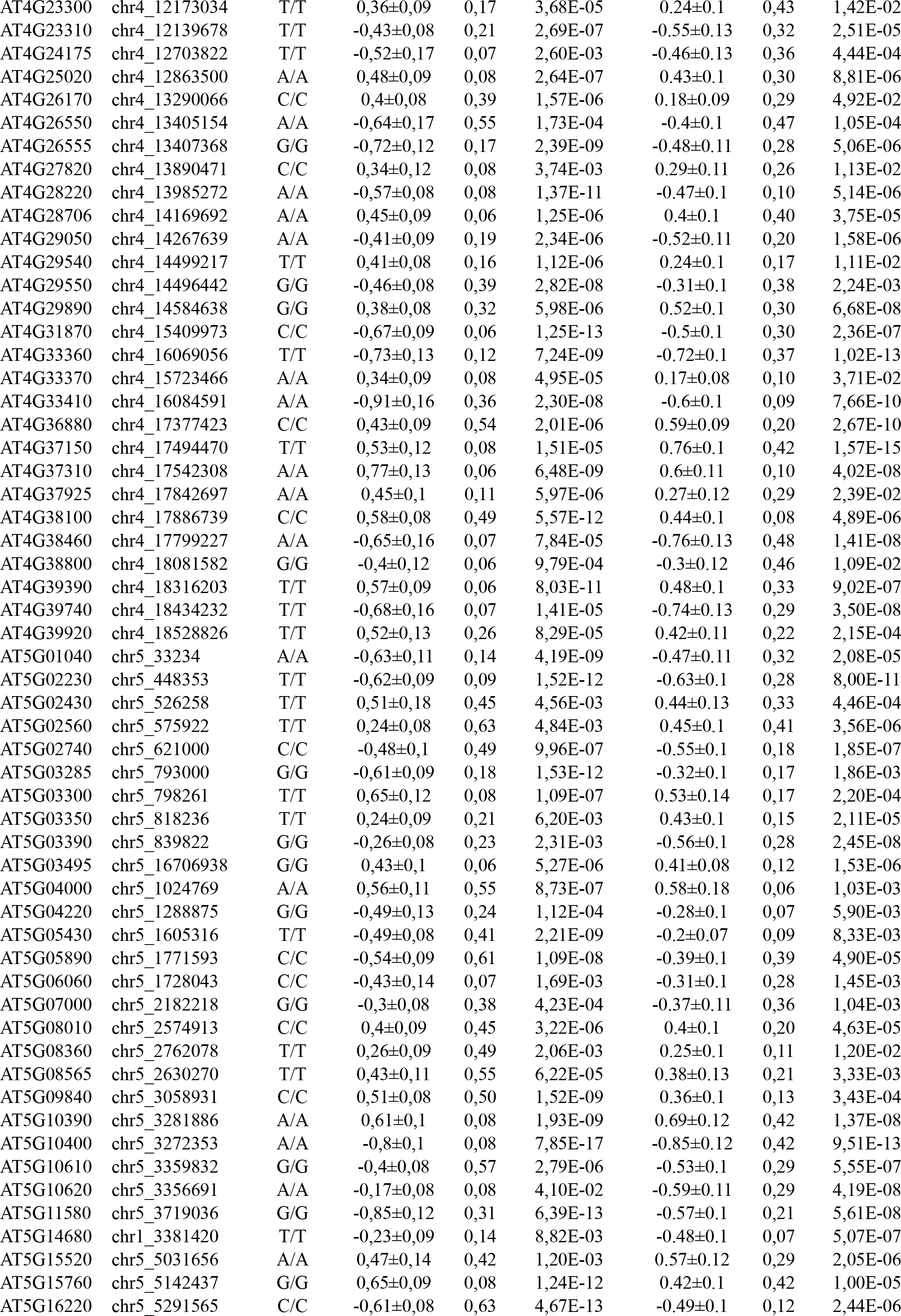
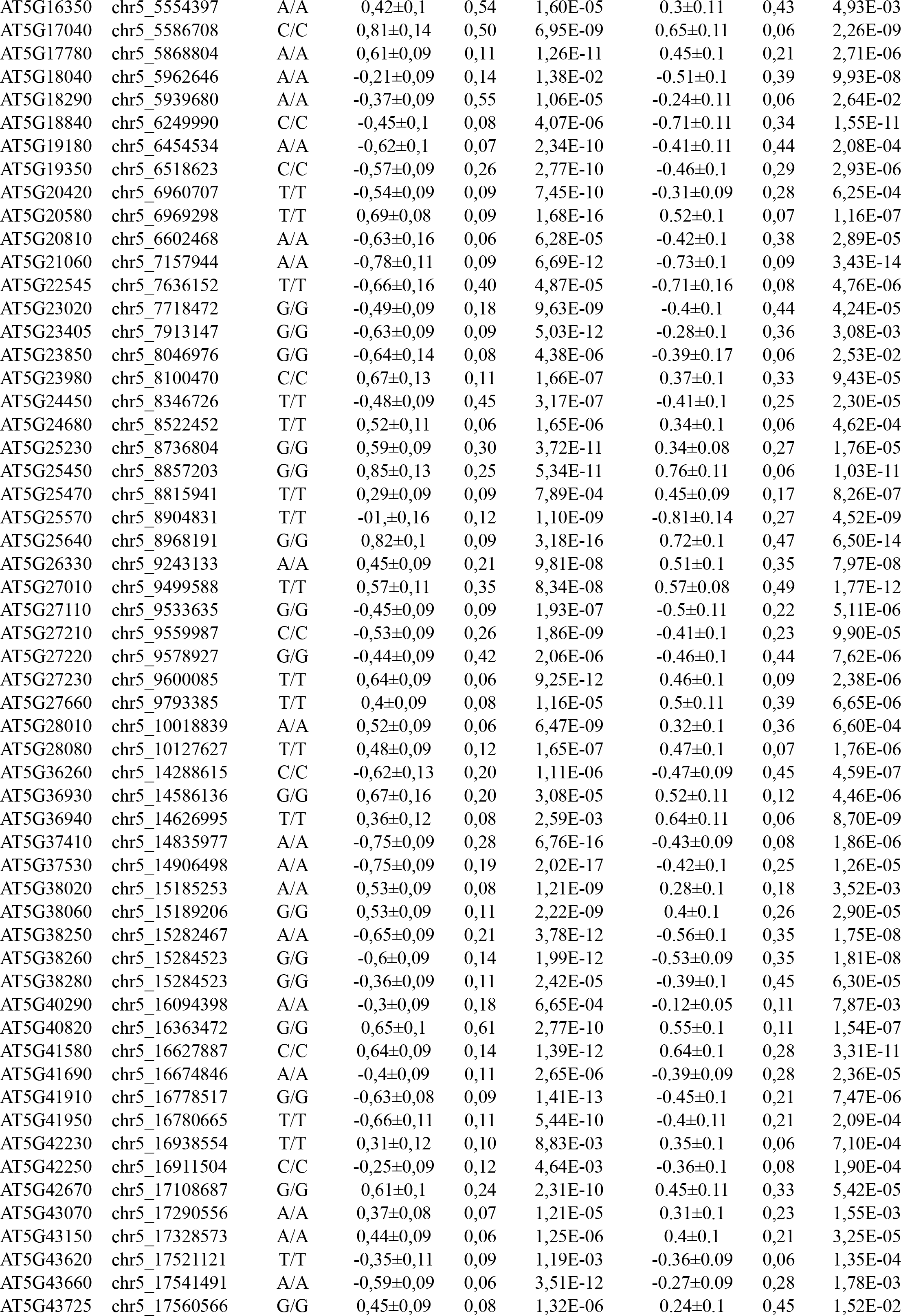
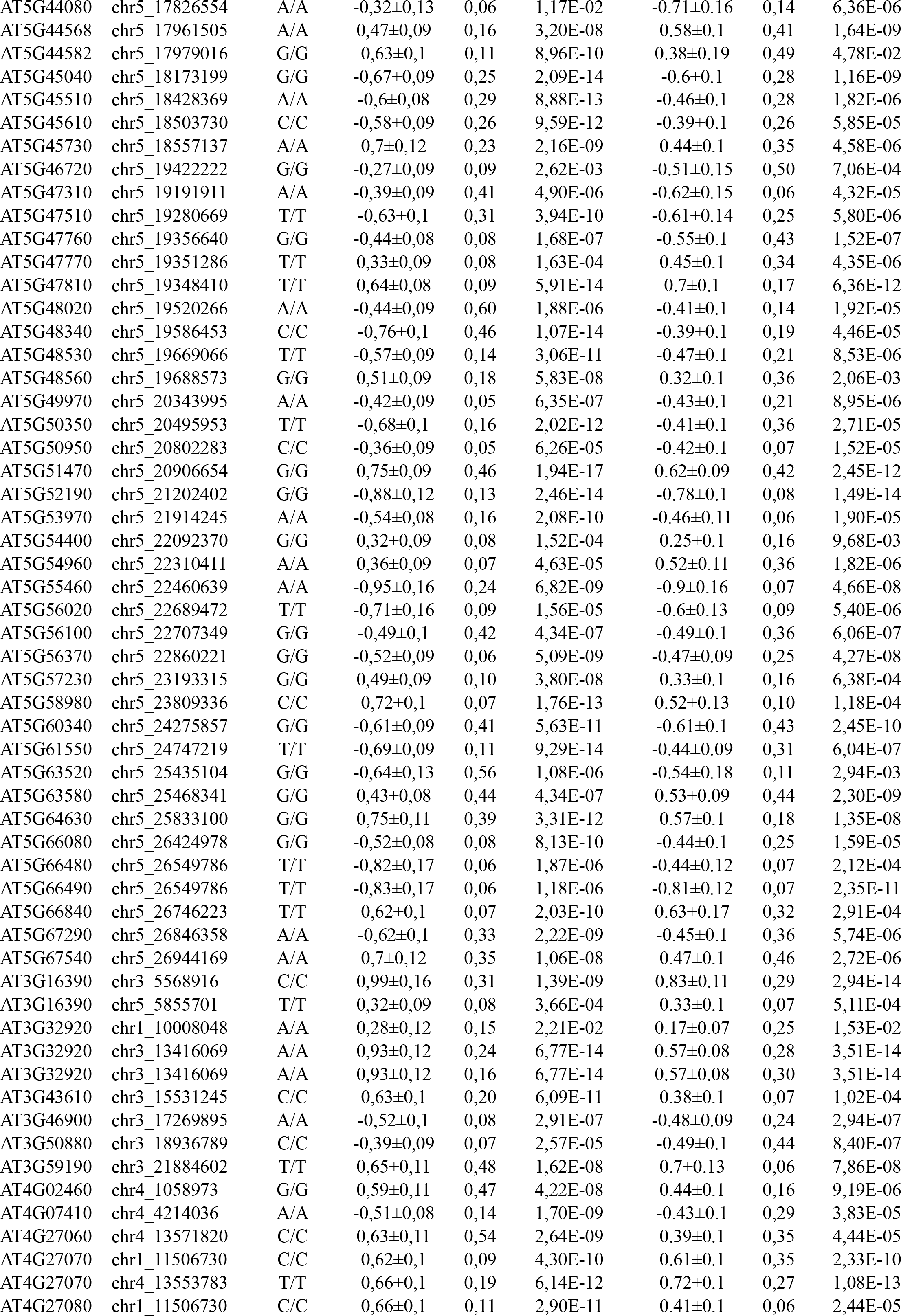
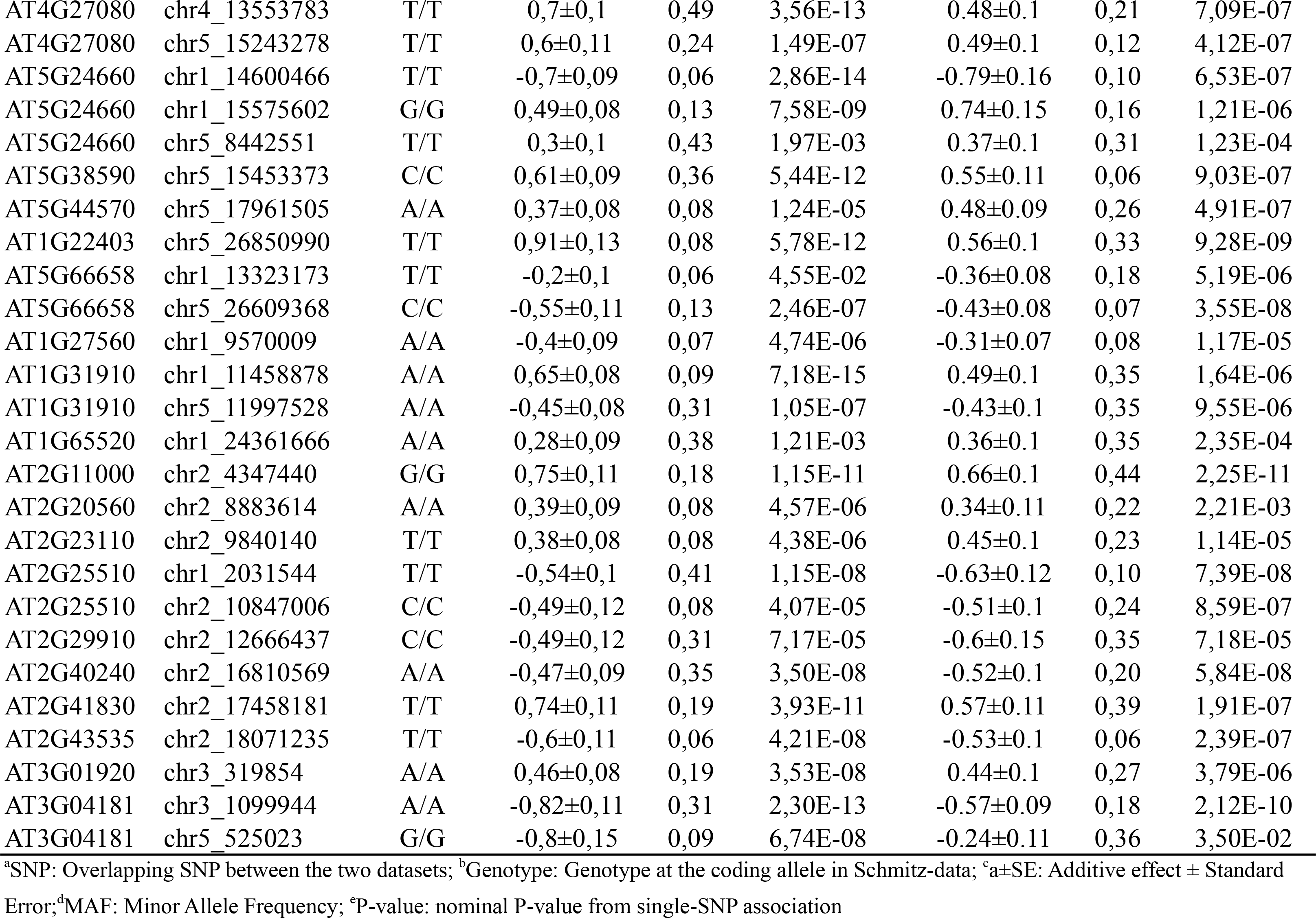
649 cis-eQTL detected in the population of 140 natural *A. thaliana* accessions that were replicated in the population of 107 Swedish natural *A. thaliana* accessions.

**Table S7.**
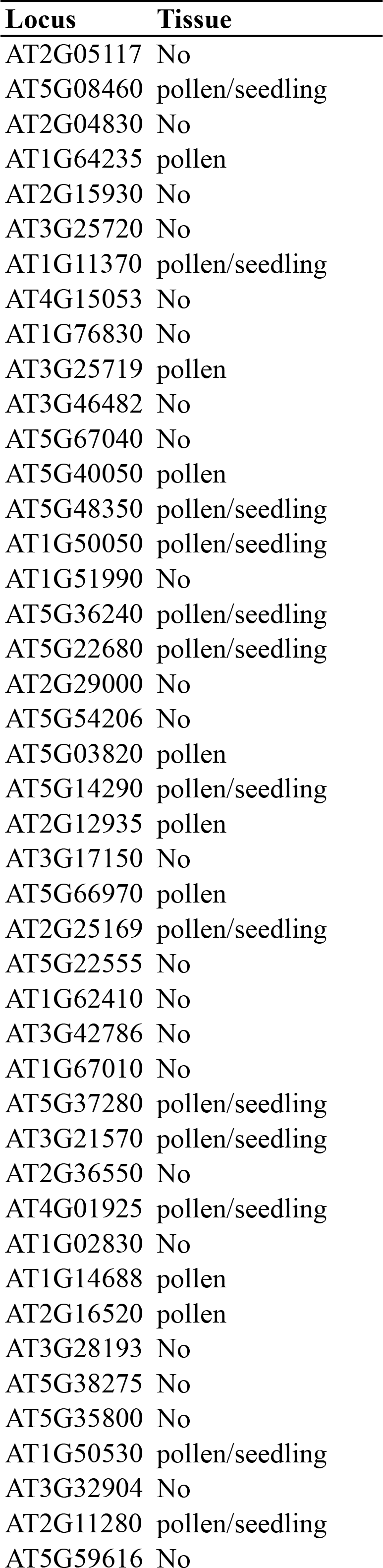
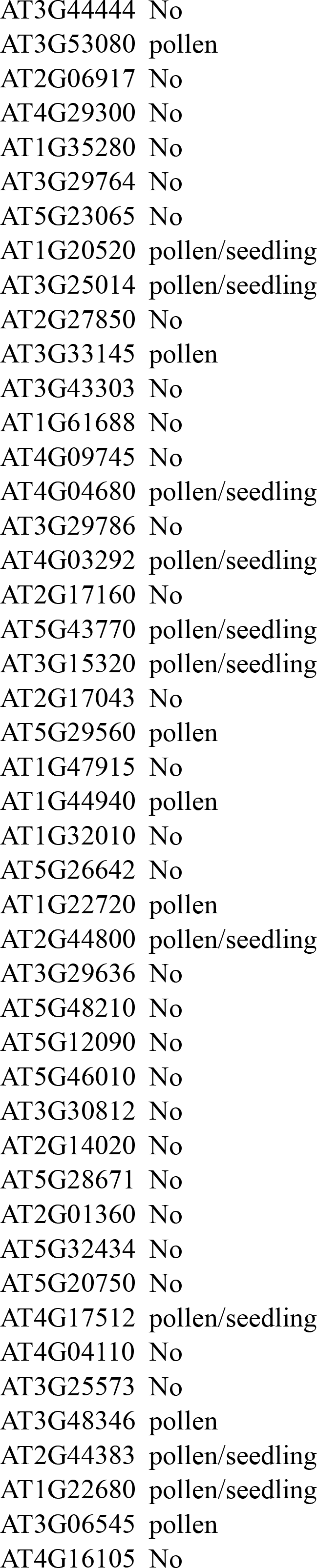
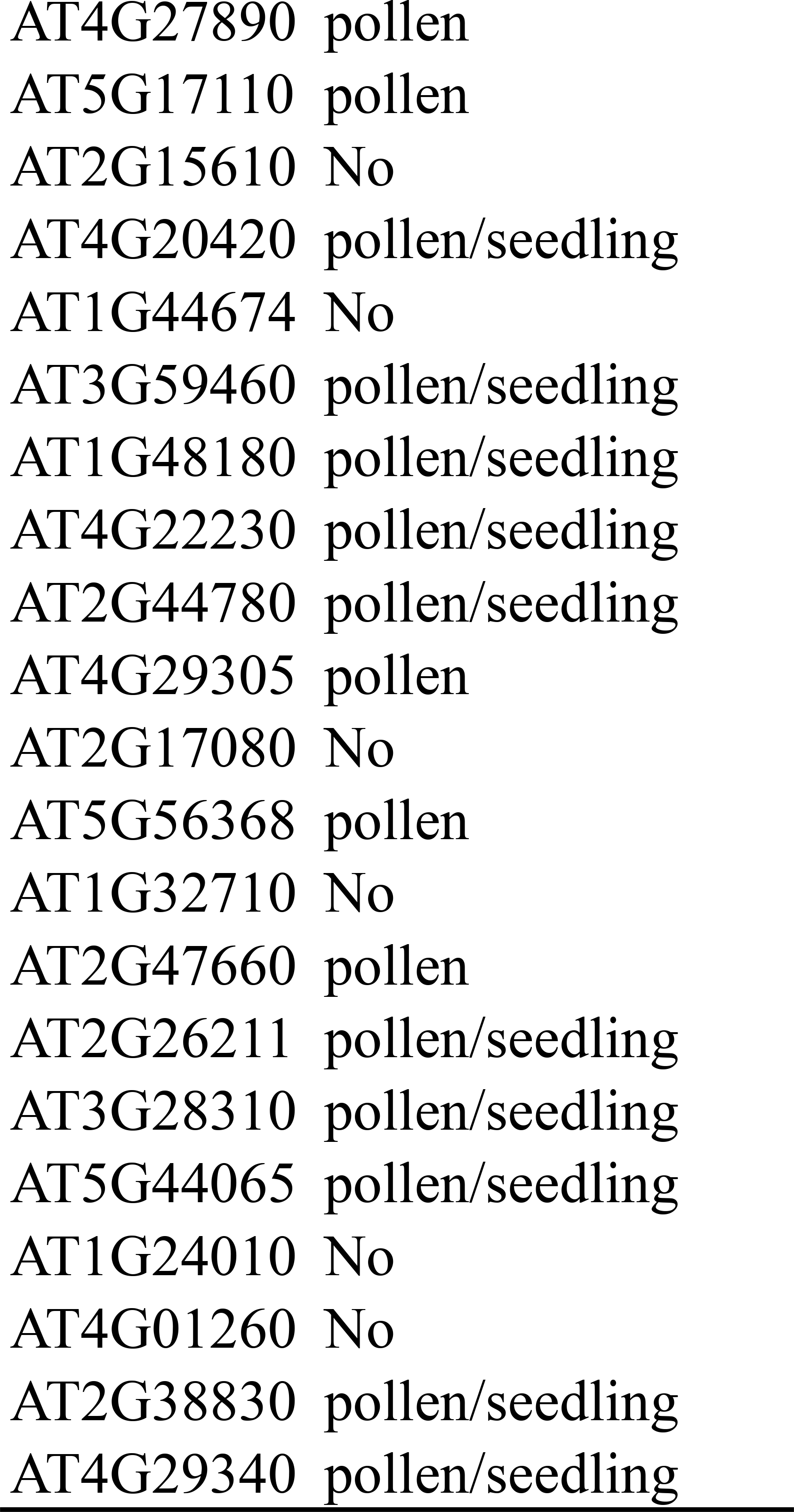
Tissue specific expression pattern for 111 genes with loss-of-expression cis-QTL and no transcripts in the leaf of *Col-0.*

#### Supplementary Figure legends

Geographical locations of 140 accessions in Schmitz-data

Illustration of 155 genes with detected cis-eQTLs and no mapped reads. A) Histogram of the number of genes with no mapped reads in genome sequencing for each accession. B) Histogram of the number of accessions for genes with no mapped reads.

## Acknowledgments

We thank Lars Hennig, David Salt, Daniel Kliebenstein and Detlef Weigel for valuable comments and suggestions during the analyses.

### Availability of data

Whole genome re-sequencing data for a population of 144 natural A. thaliana accessions (http://signal.salk.edu/atg1001/download.php) Whole genome RNA sequencing data for a population of 144 natural A. thaliana accessions http://www.ncbi.nlm.nih.gov/geo/query/acc.cgi?acc=GSE43858) Whole genome re-sequencing data for a Swedish population (https:/github.com/Gregor-Mendel-Institute/swedish-genomes) Whole genome RNA sequencing data for a Swedish population http://www.ncbi.nlm.nih.gov/geo/query/acc.cgi?acc=GSE54680)

### Abbreviations

ELPs: Expression Level Polymorphisms; eQTL: expression quantitative trait locus; FPKM: fragment per kilo base of exon per million fragments mapped; RPKM: Reads per kilobase per million reads mapped; HAC1: High Arsenic Content 1.

### Competing interests

The authors declare that they have no competing interests.

### Authors’ contributions

ÖC and XS initiated the study; ÖC and YZ developed the study, designed the analyses and summarised the results; YZ performed the analysis with support from SKGF and XS; ÖC and YZ wrote the manuscript. All authors read, commented and approved the final manuscript.

